# Multiplex genotyping method to validate the multiallelic genome editing outcomes using machine learning-assisted long-read sequencing

**DOI:** 10.1101/2020.12.14.422641

**Authors:** Akihiro Kuno, Yoshihisa Ikeda, Shinya Ayabe, Kanako Kato, Kotaro Sakamoto, Sayaka Suzuki, Kento Morimoto, Arata Wakimoto, Natsuki Mikami, Miyuki Ishida, Natsumi Iki, Yuko Hamada, Megumi Takemura, Yoko Daitoku, Yoko Tanimoto, Tra Thi Huong Dinh, Kazuya Murata, Michito Hamada, Masafumi Muratani, Atsushi Yoshiki, Fumihiro Sugiyama, Satoru Takahashi, Seiya Mizuno

**Affiliations:** Department of Anatomy and Embryology, Faculty of Medicine, University of Tsukuba, 1-1-1 Tennodai, Tsukuba, Ibaraki 305-8575, Japan; Ph.D Program in Human Biology, School of Integrative and Global Majors, University of Tsukuba, 1-1-1 Tennodai, Tsukuba, Ibaraki 305-8575, Japan; Doctoral program in Biomedical Sciences, Graduate School of Comprehensive Human Sciences, University of Tsukuba, 1-1-1 Tennodai, Tsukuba, Ibaraki 305-8575, Japan; Laboratory Animal Resource Center, Transborder Medical Research Center, Faculty of Medicine, University of Tsukuba, 1-1-1 Tennodai, Tsukuba, Ibaraki 305-8575, Japan; Experimental Animal Division, RIKEN BioResource Research Center, 3-1-1 Koyadai, Tsukuba, Ibaraki 305-0074, Japan; Department of Computer Science, University of Tsukuba, 1-1-1 Tennodai, Tsukuba, Ibaraki 305-8575, Japan; Bioinformatics Laboratory, Faculty of Medicine, University of Tsukuba, 1-1-1 Tennodai, Tsukuba, Ibaraki 305-8575, Japan; Doctoral program in Medical Sciences, Graduate School of Comprehensive Human Sciences, University of Tsukuba, 1-1-1 Tennodai, Tsukuba, Ibaraki 305-8575, Japan; Department of Genome Biology, Faculty of Medicine, University of Tsukuba, 1-1-1 Tennodai, Tsukuba, Ibaraki 305-8575, Japan

## Abstract

Genome editing can introduce designed mutations into a target genomic site. Recent research has revealed that it can also induce various unintended events such as structural variations, small indels, and substitutions at, and in some cases, away from the target site. These rearrangements may result in confounding phenotypes in biomedical research samples and cause a concern in clinical or agricultural applications. However, current genotyping methods do not allow a comprehensive analysis of diverse mutations for phasing and mosaic variant detection. Here, we developed a genotyping method with an on-target site analysis software named Determine Allele mutations and Judge Intended genotype by Nanopore sequencer (DAJIN) that can automatically identify and classify both intended and unintended diverse mutations, including point mutations, deletions, inversions, and *cis* double knock-in at single-nucleotide resolution. Our approach with DAJIN can handle approximately 100 samples under different editing conditions in a single run. With its high versatility, scalability, and convenience, DAJIN-assisted multiplex genotyping may become a new standard for validating genome editing outcomes.

## INTRODUCTION

The development of new technologies such as CRISPR-Cas has facilitated genome editing of any species or cell type. Nucleases such as Cas9 and FokI and deaminase fused with Cas9 have been used to introduce DNA double-strand breaks and perform base editing, respectively (**1-3**). However, as double-strand break repair pathways are regulated by host cells (4), verifying the result and selecting desired mutated alleles for precise genome editing are essential. Multiple alleles exist in a population of cells or individual animals that have undergone genome editing. In most cases, animals born following editing events at early embryonic stages are mosaic (5). Heterogeneous cell populations can be obtained by genome editing of cultured cells or delivering genome editing tools to somatic cells (6, 7).

Cell populations with incorrectly edited alleles need to be detected and excluded to ensure precise genome editing (8). Unintended alleles with similar genetic impact may be tolerated only in a specific purpose of genome editing, e.g. generation of null alleles through the deletion of critical exon(s) by using multiple guide RNAs, resulting in multiple patterns in the total deleted length (9). Recent studies have found that genome editing can induce various on-target events such as inversions, deletions, and endogenous and exogenous DNA insertions as well as indels and substitutions at, and in some cases, away from the target site (10-13). Furthermore, there is a possibility of gene conversion between homologous regions following genomic DNA cleavage (14-16).

The assessment of on-target editing outcomes and the selection of correct, precisely edited alleles lead to efficient production and breeding of founder animals and their offspring as well as efficient *in vivo* and *ex vivo* engineering. Demultiplexing of highly homologous mutated alleles is required to separate the signals of each allele from genetically engineered samples. However, the subcloning of amplified products is laborious, and short-range assessments with targeted PCR amplification and tracking of indels by decomposition analysis of Sanger sequencing data are likely to miss long-range mutation events, which may result in pathogenic phenotypes through unintended changes in gene expression (17,18). Moreover, short-range PCR analysis followed by NGS cannot identify multiple intended or unwanted mutations in *cis* or in *trans* (19,20). Long-read sequencing technologies enable a comprehensive analysis of the region of interest by providing longer sequence reads compared to the traditional strategy and make it possible to identify unexpected genome editing outcomes, including complex structural variations (10,13). Although targeted long-read sequencing allows the detection of complex on-target mutations over several kilobases (13,21), this method has instrumental limitations such as error rates and lack of tools for phasing and mosaic variant detection to validate multiple and diverse allelic variants to a single-base level (22). Thus, more accessible and high-throughput methods for routine assessment of genome editing outcomes are essential to detect the unpredictable editing events.

Herein, we describe a novel method for analysing genome editing outcomes, in which long-chain PCR products with barcodes obtained using two-step long-range PCR were used as samples, and allele validation was performed using our original software named Determine Allele mutations and Judge Intended genotype by Nanopore sequencer (DAJIN) that enables the comprehensive analysis of long reads generated using the nanopore long-read sequencing technology. DAJIN, a machine-learning-based model, identifies and quantifies allele numbers and their mutation patterns and reports consensus sequences to visualise mutations in alleles at single-nucleotide resolutions. Moreover, it allows multiple sample processing, and approximately 100 samples can be processed within a day. Because of these features, our strategy with DAJIN can validate the quality of genome-edited samples to select animals or clones with intended results efficiently, and as such has the potential to contribute to more precise genome editing.

## METHODS

### Animals

ICR and C57BL/6J mice were purchased from Charles River Laboratories Japan, Inc. (Yokohama, Japan). C57BL/6J-*Tyr*^*em2Utr*^mice were provided by RIKEN BRC (#RBRC06459). Mice were kept in plastic cages under specific pathogen-free conditions in a room maintained at 23.5 ± 2.5 °C and 52.5 ± 12.5 % relative humidity under a 14-h light:10-h dark cycle. Mice had free access to commercial chow (MF diet; Oriental Yeast Co., Ltd., Tokyo, Japan) and filtered water. All animal experiments were performed humanely with the approval from the Institutional Animal Experiment Committee of the University of Tsukuba following the Regulations for Animal Experiments of the University of Tsukuba and Fundamental Guidelines for Proper Conduct of Animal Experiments and Related Activities in Academic Research Institutions under the jurisdiction of the Ministry of Education, Culture, Sports, Science, and Technology of Japan.

### Genome editing in mouse zygotes

Mice with point mutations (PMs) and two-cut knockout (KO) were generated using the electroporation method (23). The guide RNA (gRNA) target sequences to induce each mutation are listed in Additional file 1: Table. S1. The gRNAs were synthesised and purified using a GeneArt™ Precision gRNA Synthesis Kit (Thermo Fisher Scientific, Waltham, MA, USA) and dissolved in Opti-MEM (Thermo Fisher Scientific). In addition, we designed three single-strand oligodeoxynucleotides (ssODN) donors for inducing PMs in *Tyr* (Additional file 1: Table. S1). These ssODN donors were ordered as Ultramer DNA oligos from Integrated DNA Technologies (Coralville, IA, USA) and dissolved in Opti-MEM. The mixtures of gRNA (5 ng/μL) and ssODNs (100 ng/μL) or mixtures of two gRNAs (25 ng/μL each) were used to generate point mutant mice or two-cut KO mice, respectively. GeneArt Platinum Cas9 Nuclease (100 ng/μL; Thermo Fisher Scientific) was added to these mixtures. Pregnant mare serum gonadotropin (5 units) and human chorionic gonadotropin (5 units) were intraperitoneally injected into female C57BL/6J mice (Charles River Laboratories) with a 48-h interval. Next, unfertilised oocytes were collected from their oviducts. Then, according to standard protocols, we performed *in vitro* fertilisation with these oocytes and sperm from male C57BL/6J mice (Charles River Laboratories). After 5 h, the above-mentioned gRNA/ssODN/Cas9 or two gRNAs/Cas9 mixtures were electroplated into the mouse zygotes using a NEPA 21 electroplater (NEPAGNENE; Chiba, Japan), under previously reported conditions (24). The electroporated embryos that developed into the two-cell stage were transferred to oviducts of pseudopregnant ICR female mice. The floxed mice were generated using the microinjection method (25). Each gRNA target sequence (Additional file 1: Table. S5) was inserted into the entry site of pX330-mC carrying both the gRNA and Cas9 expression units. These pX330-mC plasmid DNAs and donor DNA plasmid were isolated using FastGene Plasmid Mini kit (Nippon Genetics, Tokyo, Japan) and filtered using MILLEX-GV® 0.22 μm filter unit (Merck Millipore, Darmstadt, Germany) for microinjection. Next, C57BL/6J female mice superovulated using the method described above were naturally mated with male C57BL/6J mice, and zygotes were collected from the oviducts of the mated female mice. For each gene, a mixture of two pX330-mC (circular, 5 ng/μL each) and a donor (circular, 10 ng/μL) was microinjected into the zygote. The zygotes that survived were then transferred into the oviducts of pseudopregnant ICR female mice. When the newborns were around 3 weeks of age (Additional file 1: Table. S2), the tail was sampled to obtain genomic DNA.

### Library preparation and nanopore sequencing

We used PI-200 (KURABO INDUSTRIES LTD., Osaka, Japan), according to the manufacturer’s protocol, for the extraction and purification of genomic DNA obtained from the tail of mice. The purified genomic DNA was amplified using PCR using KOD multi & Epi (TOYOBO, Osaka, Japan) and target amplicon primers (Additional file 1: Table. S3). In the target amplicon primer, the universal sequence is located on the 5′ side, and the sequence for target gene amplification is on the 3′ side. Five-fold dilutions of the PCR products were used as templates for nested PCR performed using KOD multi & Epi and barcode attachment primers (Additional file 1: Table. S4). The 5′ side of the barcode attachment primer has a barcode sequence, and the 3′ sequence is annealed to the universal sequence of the target amplicon primer. The barcoded PCR products were mixed in equal amounts and then purified using FastGene Gel/PCR Extraction Kit (Nippon Genetics, Germany). The volume of the mixed and purified PCR products was adjusted to 20–30 ng/μL. The library was prepared using Ligation Sequencing 1D kit (SQK-LSK108_109; ONT, Oxford, UK) and NEBNext End repair/dA-tailing Module NEB Blunt/TA Ligase Master Mix (New England Biolabs, Ipswich, MA, USA) according to the manufacturer’s instructions. The prepared library was loaded onto a primed R9.4 Spot-On Flow cell (FLO-MIN106; ONT, Oxford, UK). The 24 h or 36 h run time calling sequencing protocol was selected in the MinKNOW GUI (version 4.0.20), and base calling was allowed to complete after the sequencing run was completed. After base calling, we demultiplexed the barcoding libraries using qcat (version 1.1.0) with default parameter settings. Total nanopore sequencing reads per sample are listed in Additional file 1: Table. S5.

### Conventional genotyping analysis

To evaluate the validity of DAJIN’s genotyping results, we used conventional genotyping methods, including short-amplicon PCR, PCR-RFLP, and Sanger sequencing. For the genotyping of the two-cut KO and PM lines, genomic PCR was performed using AmpliTaq Gold 360 DNA Polymerase (Thermo Fisher Scientific) and the relevant primers (Additional file 1: Table. S6). Agarose gel electrophoresis was performed to confirm the size of the PCR products. In the flox knock-in design, genomic PCR was performed using KOD FX (TOYOBO) and the relevant primers (Additional file 1: Table. S6). The PCR products were digested with restriction enzymes *Asc*I (New England Biolabs) and *Eco*RV (New England Biolabs) for 2 h to check LoxP insertion on the left and right side, respectively. Agarose gel electrophoresis was performed to confirm the size of the PCR fragments. PCR products with mutant sequences were identified using Sanger sequencing using the BigDye™ Terminator v3.1 Cycle Sequencing Kit (Thermo Fisher Scientific).

### Targeted next-generation sequencing

Genomic PCR was performed using AmpliTaq Gold 360 DNA Polymerase (Thermo Fisher Scientific) and the relevant primers whose barcode sequences were added to the 5′ end (Additional file1: Table. S7). The PCR amplicons in 226 bp (Tyr.c140 G>C) and 203 bp (*Tyr* c.316 G>C, *Tyr* c.308 G>C) lengths were purified using FastGene Gel/PCR Extraction Kit (Nippon Genetics, Düren, Germany). Paired-end sequencing (2 × 151 bases) with these purified amplicons was performed using MiSeq (Illumina, San Diego, CA, USA) at Tsukuba i-Laboratory LLP (Tsukuba, Ibaraki, Japan). Paired-end reads were mapped against chromosome 7 of mouse genome assembly mm10 using STAR (version2.7.0a) with default settings (26). Mapped reads were visualised using IGV (version 2.9.4) (27). The samples carrying the intended point mutation at frequencies >10% were considered as positive.

### Nanopore read simulation

To prepare training data for deep neural network (DNN) models, we generated simulation reads of the possible alleles using NanoSim (version 2.5.0) (28). We trained NanoSim to obtain an error profile using nanopore sequencing reads from a WT control. Next, we applied the error profile to generate 10,000 simulation reads per each possible allele that could be caused by genome editing (Additional file 2: Fig. S1). In the PM design, we generated simulation reads with a deletion or random nucleotide insertion of the gRNA length at the Cas-cutting site.

### Pre-processing

We performed pre-processing to exclude reads without target loci and to perform MIDS conversion. First, the genome-edited sequence was aligned to the user-provided WT sequence using minimap2 (version 2.17) (29) with the ‘--cs=long’ option, and the position of the target mutant base was detected according to the CS-tag in the SAM file. Simulated and nanopore sequencing reads were then aligned using minimap2 to the WT sequence. Reads with lengths more than 1.1 times longer than the maximum length among possible alleles were excluded. For the remaining reads, we detected the start and end positions of each read relative to the WT sequence based on CIGAR information and extracted the reads containing the mutant region of interest (Additional file 2: Fig. S4a).

The extracted reads were subjected to MIDS conversion (Additional file 2: Fig. S4b). The matched, inserted, deleted, and substituted bases compared to the control sequence were converted to M (Match), I (Insertion), D (Deletion), and S (Substitution), respectively. Next, the read lengths were trimmed or padded with ‘=’ to equalise their sequence length. Then one-hot encoding was performed on the MIDS sequence.

### Deep learning model

We constructed a deep neural network (DNN) model to classify alleles. The structure of the deep learning model is shown in Additional file 2: Fig. S5. The architecture comprises three layers of convolutional and max-pooling layers and a fully connected layer, and a softmax function to predict the allele types. The batch size was 32. The maximum number of training epochs was 200, and the training was stopped when validation loss was not improved during 20 epochs. To detect SV reads, we extracted the outputs from the fully connected (FC) layer. Then we trained the Local Outlier Factor (LOF) (30) using the output of the simulated reads. Subsequently, the output of the nanopore sequence reads was placed in the LOF; it annotated unexpected mutation reads as “SV”. We assessed the accuracy of the classification using simulation reads, and it was able to accurately classify alleles in all genome editing designs conducted in this study (Additional file 1: Table. S8).

### Allele clustering

In order to distinguish each allele precisely, DAJIN conducts compressed MIDS conversion and clustering. To generate fixed-length sequences, we performed compressed MIDS conversion, which replaces successive insertions with a character corresponding to the number of insertions and then substitutes the insertion (Additional file 2: Fig. S6). A character is assigned to the number from 1 to 9 or a letter from a to z. If the number of consecutive insertions is in the range 1–9, the character is the corresponding number. If the number of consecutive insertions is in the range 10–35, the character is ‘a’ (=10) to ‘y’ (=35). If the number is greater than 35, the character is ‘z’ (>35).

To mitigate nanopore sequencing errors, the MIDS’s relative frequencies of the sample were subtracted from the control applied to each read, called MIDS score (Additional file 2: Fig. S7). The MIDS score was reduced into 10 dimensions using principal component analysis. Then Hierarchical Density-based Spatial Clustering of Applications with Noise (HDBSCAN) (31) was performed for the clustering. For parameters, we set ‘min_samples’, which specifies the minimum size of each cluster formed, as ‘1’ to reduce noise points. Furthermore, we tuned ‘min_cluster_size’, which defines the minimum number of samples in each cluster. We set the value as 50 equal intervals between 1/5 and 2/5 of the total number of reads and then selected the ‘min_cluster_size’, which outputs the mode of cluster numbers.

### Filter minor alleles

To improve the interpretability, DAJIN has a default setup to remove minor alleles. Minor alleles were defined as those in which the number of reads was 1% or less of the total number of reads of a sample. DAJIN was able to report all allele information using the ‘filter=off’ option.

### Consensus sequence

The consensus sequence for each allele was output as a FASTA file and an HTML file. In the HTML file, the mutated nucleotides are coloured. To generate the consensus sequence, we compared FASTA alleles and compressed MIDS sequences.

### Generation and Visualisation of BAM files

DAJIN generates BAM files to visualise the DAJIN-reported alleles in a genome browser. First, DAJIN uses minimap2 to map the nanopore sequence reads to the WT sequences described in the user-inputted FASTA file, then samtools (version 1.10) (32) generates sorted BAM files. Next, the target genome coordinates and chromosome length are obtained from the UCSC Table Browser (33) according to the user-inputted FASTA file and genome assembly ID. Then DAJIN replaces the chromosome number and chromosome length in SN and LN headers of BAM files.

### Single-nucleotide variant (SNV) and structural variation (SV) callers

We installed Medaka (version 1.2.1) (34), Clair (version 2.1.1) (35), NanoSV (version 1.2.4) (36), NGMLR (version 0.2.7) (37), Sniffles (version 1.0.12) (37) via Bioconda (38). For Sniffles, the parameters of ‘—cluster’ and ‘-n -1’ were provided to phase SVs in all sequencing reads, otherwise the default parameters were chosen.

## RESULTS

### Workflow of DAJIN

We designed DAJIN to genotype genome-edited samples by capturing diverse mutations from single-nucleotide variants (SNV) to structural variations (SV). The overall workflow of DAJIN is presented in Fig. 1a. DAJIN requires: 1) a FASTA file describing possible alleles, which must include the DNA sequence before and after genome editing; 2) FASTQ files from nanopore sequencing, which include a control sample; 3) gRNA sequence including the protospacer adjacent motif (PAM); and 4) a genome assembly ID such as hg38 and mm10. Next, DAJIN generates simulation reads using NanoSim (28) according to the user-inputted FASTA file. The sequence reads are pre-processed, and one-hot encoded. Subsequently, the simulated reads are used to train a deep neural network (DNN) model to detect SV reads and classify allele types. DAJIN defines SV alleles as a different sequence from the user-inputted FASTA file. Next, DAJIN conducts clustering to estimate the alleles. Finally, it reports the consensus sequence to visualise the mutations in each allele and labels the alleles. The details are described in Methods and Additional file 2: Figs. S1,4–7. The outputs of DAJIN are shown in Fig. 1b. DAJIN reports allele frequencies in each sample, the consensus sequences, and BAM files for each allele. In this study, DAJIN was evaluated on nine mouse strains of three types of genome editing design: point mutation (PM), 2-cut knockout (KO), and flox knock-in (KI). The performance evaluations are described in detail below.

**Figure 1.**
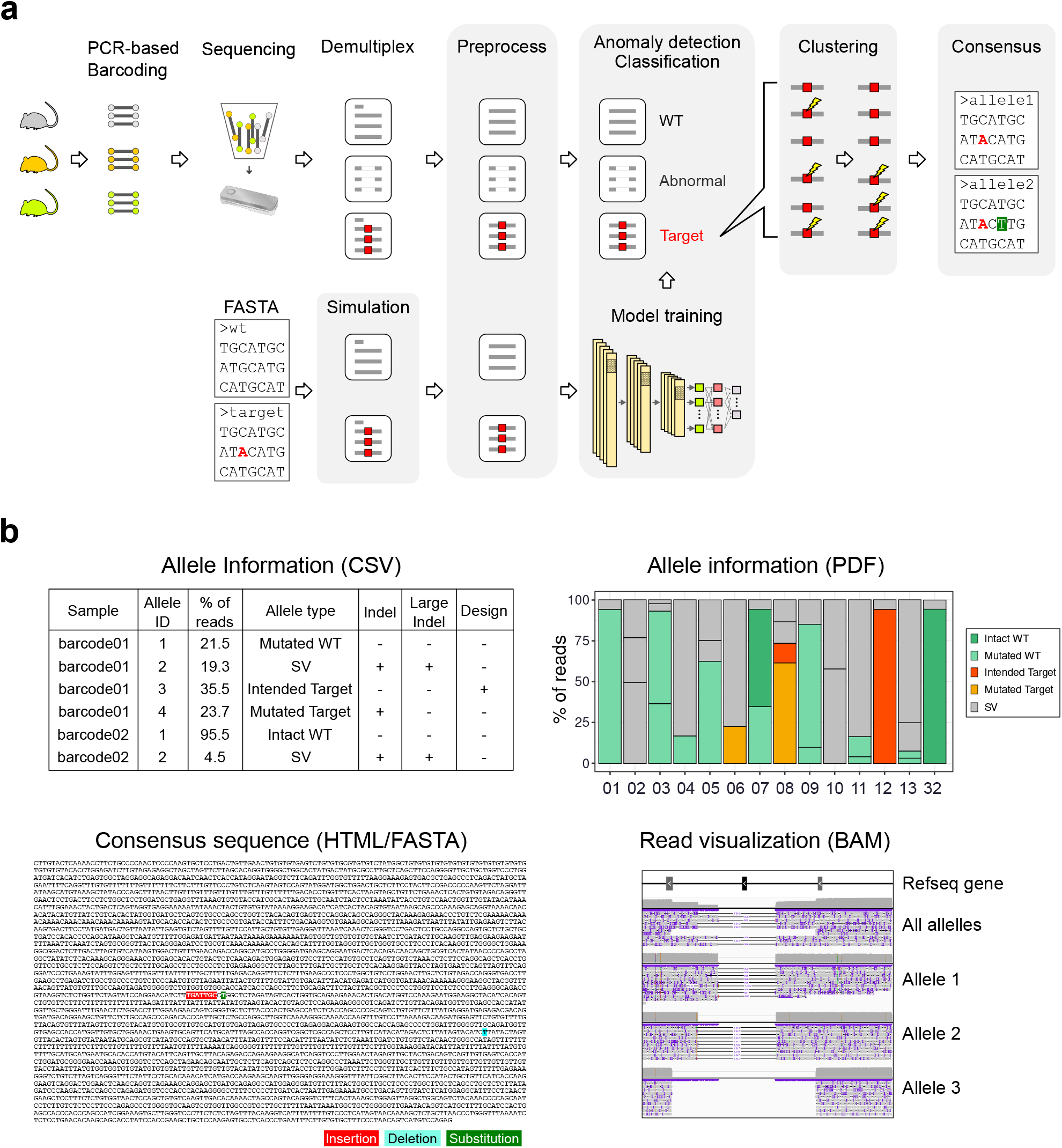
Overview of the methods. **a** The schematic of DAJIN’s workflow. DAJIN automates the procedures highlighted in grey. Red-coloured nucleotides represent intended point mutation. A green highlighted nucleotide represents unintended substitution. **b** The outputs of DAJIN. The file formats are described in parentheses. SV: structural variations.

### Performance of SV detection

CRISPR-Cas genome editing has been reported to induce unexpectedly large indels, which might be overlooked by conventional short PCR-based genotyping methods. Conversely, a nanopore long-read sequencer can capture large indels within its amplicon size, which allows the detection of SV alleles. Thus, we implemented an SV detection using DNN (See Methods in detail) into DAJIN and evaluated its performance. We simulated nanopore reads with a deletion in the range of 10 bp to 200 bp at the *Cables2* genome locus (Additional file 2: Fig. S2a). Next, we pre-processed the simulated reads with or without MIDS conversion and conducted Uniform Manifold Approximation and Projection (UMAP) (39) and Local Outlier Factor (LOF) (30) using outputs from the last fully connected (FC) layer (Additional file 2: Fig. S2b). The UMAP revealed the cluster of 50 bp deletion with MIDS conversion, which was unclear without MIDS conversion (Additional file 2: Fig. S2c). We next investigated the accuracy of SV allele detection. The results showed that DAJIN labelled more than 50 bp deletion as SV with MIDS conversion, but not without, which indicates that the MIDS conversion improves SV allele detection (Additional file 2: Fig. S2d). Since several SV callers using long-read sequencing have been developed, we next compared DAJIN to NanoSV (36) and Sniffles (37). We prepared 1000 simulated reads containing deletions of 50, 100, and 200 bases for each allele and mixed them to imitate genome-edited samples with the three alleles (Additional file 2: Fig. S3a). Then we evaluated the samples using DAJIN, NanoSV, and Sniffles. The results showed that DAJIN discriminated each allele according to the deletion sizes; however, NanoSV and Sniffles did not (Additional file 2: Fig. S3b).

### DAJIN captures point mutation alleles

Next, we evaluated DAJIN’s performance using genome-edited mice. We induced *Tyr* c.140G>C point mutation (PM) using CRISPR-Cas9 genome editing in C57BL/6J mouse fertilised eggs, and obtained 13 founder mice. Next, we amplified a 2845 bp DNA sequence at the *Tyr* loci of the founder mice (barcode (BC) 01–13) and a WT control mouse (BC32) (Fig. 2a). Then, DAJIN was used to analyse the PCR amplicons of the 14 mice (Fig. 2b). Because the PM’s genome-editing design potentially generates WT, PM, and unexpected SV alleles, DAJIN annotated ‘WT’, ‘PM’, and ‘SV’ allele types.Besides, when DAJIN’s consensus sequence perfectly matched the sequences of ‘WT’ and ‘PM’ described in the user-inputted FASTA file, these alleles were labelled as ‘Intact WT’ and ‘Intended PM’, respectively, whereas when there was a mismatch between DAJIN’s consensus sequence and FASTA sequence, it was labelled as ‘Mutated WT’ and ‘Mutated PM’.

**Figure 2.**
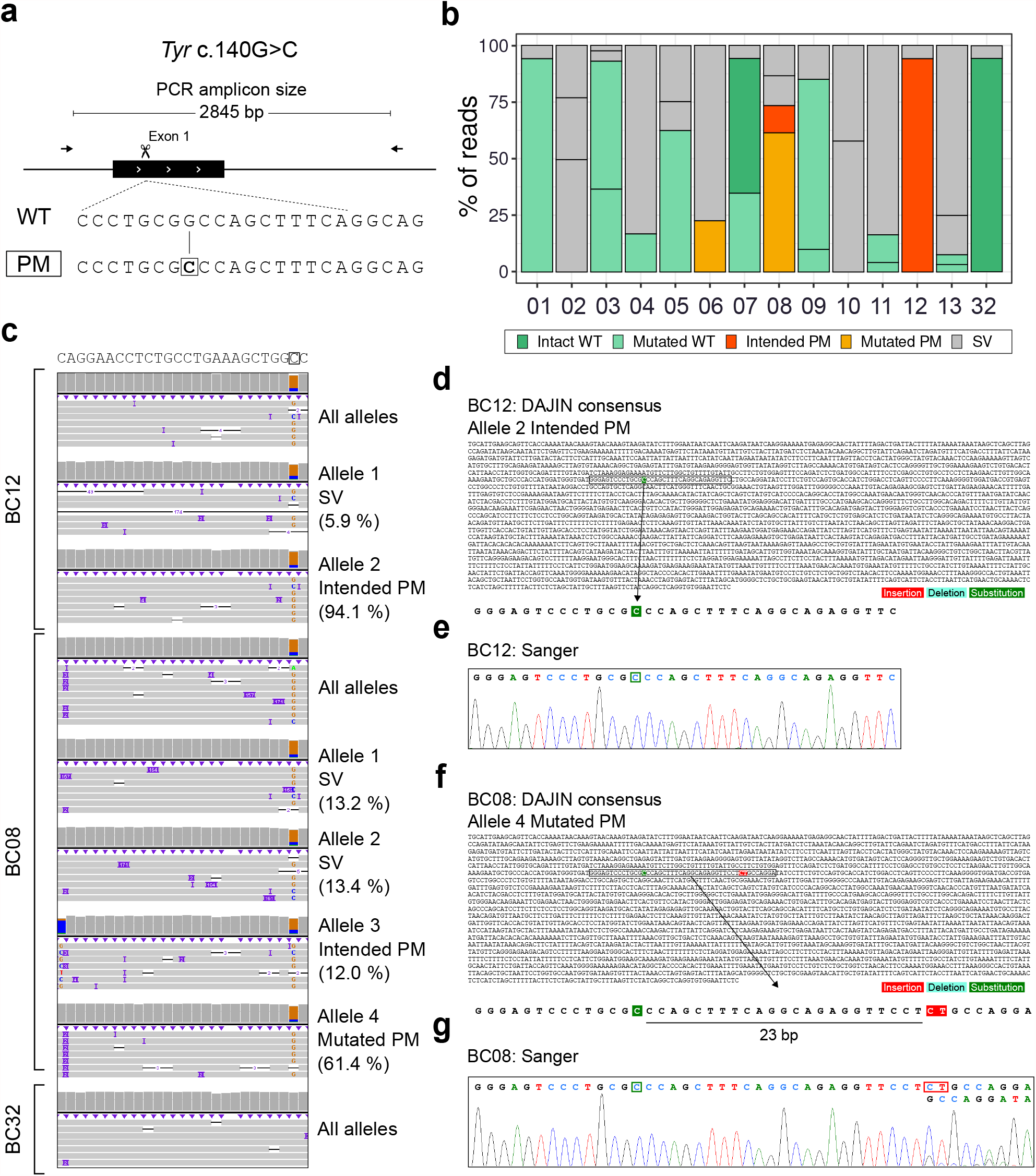
Application of DAJIN for point mutation design. **a** Genome editing design for *Tyr* c.140G>C point mutation. The scissors represent a Cas9-cutting site. The arrows represent PCR primers, including the PCR amplicon size. The boxed allele type represents the target allele, and the boxed nucleotide represents a targeted point mutation. **b** DAJIN’s report of the allele percentage. The barcode numbers on the x-axis represent mouse IDs. BC32 is a WT control. The y-axis represents the percentage of DAJIN-reported alleles. The colours of the bars represent DAJIN-reported allele types. The compartments partitioned off by horizontal lines in a bar represent the DAJIN-reported alleles. **c** Visualisation of nanopore sequencing reads at *Tyr* target locus. BC12 and BC08 contain target alleles. BC32 is a WT control. The ‘All alleles’ track represents all reads of each sample. The ‘Allele’ track represents DAJIN-reported alleles. **d** Comparison between DAJIN’s consensus sequence and Sanger sequencing. The sequence represents the consensus sequence of a dominant allele of BC12 and BC08. The colours on the nucleotides represent mutation types, including insertion (red), deletion (sky blue), and substitution (green). The coloured boxes in the Sanger sequence represent mutated nucleotides, including insertion (red) and substitution (green).

DAJIN reported the percentages of the predicted allele types and identified two mice (BC08, BC12) having the intended PM (Fig. 2b). Visualisation using IGV showed that DAJIN accurately captured the c.140G>C PM in BC08 and BC12 (Fig. 2c). Moreover, DAJIN detected an unexpected 2 bp insertion in BC08 at 23 bp downstream from the PM and labelled the allele as ‘Mutated PM’ (Fig. 2c). Next, DAJIN’s consensus sequence reported the BC12 included in the ‘Intended PM’ allele (Fig. 2d).

Sanger sequencing of BC12 at the PM locus supported the target PM’s induction (Fig. 2e). Furthermore, DAJIN’s consensus sequence of ‘Mutated PM’ allele in BC08 included an unexpected 2 bp (CT) insertion as well as the PM (Fig. 2f). The same ‘CT’ insertion was validated via Sanger sequencing (Fig. 2g). Besides, DAJIN reported the ratio of ‘CT’ inserted alleles and the other alleles was approximately 5:1 (Fig. 2c). Sanger sequencing also showed a waveform intensity ratio of 5:1 between the PM with CT insertion allele and that without the insertion allele (Fig. 2g). This consistency indicated that DAJIN accurately quantifies the allele frequencies.

To further evaluate DAJIN’s performance, we generated two more PM mice, *Tyr* c.316G>C and *Tyr* c.308G>C (Additional file 2: Fig. S8). Besides, a C57BL/6J-*Tyr*^*em2Utr*^ mouse, which carries the *Tyr* c.230G>T PM (5), was added to BC31 as a control for ‘Mutated WT’. For *Tyr* c.316G>C and c.308G>C projects, DAJIN reported that 1 out of 6 mice (BC18) and 8 out of 11 mice (BC21, BC22, BC23, BC24, BC26, BC29, and BC30) had the ‘Intended PM’ (Fig. 3a). As with BC12, DAJIN annotated almost all (93.6%) nanopore sequencing reads in BC21 as ‘Intended PM’. Subsequently, DAJIN’s consensus sequence in BC21 reported the intended c.308G>C PM, and we detected a single waveform of the PM using Sanger sequence analysis (Additional file 2: Fig. S9). In addition, DAJIN correctly identified *Tyr* c.230G>T PM in BC31, which was used as the positive control of ‘Mutated PM’ (Additional file 2: Fig. S10).

**Figure 3.**
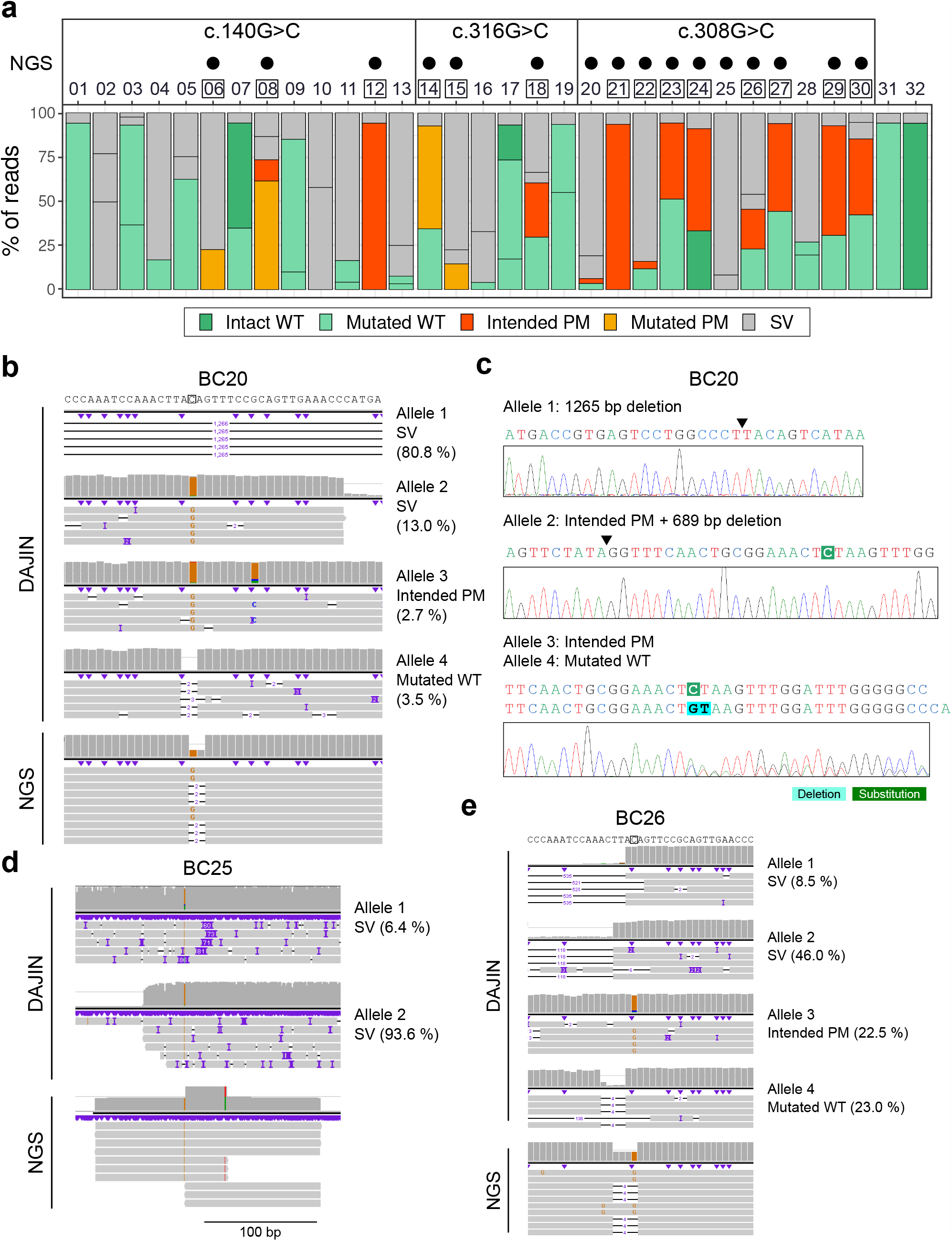
Comparison to NGS for point mutation design. **a** The genotyping results obtained using DAJIN and NGS. The numbers on the x-axis represent barcode IDs. The genome editing of BC01–BC13, BC14–BC19, and BC20–BC30 aims to induce c.140G>C, c.316G>C, and c.308G>C, respectively. The black dots represent PM positive samples detected using NGS. The boxed numbers represent PM positive samples detected using DAJIN. The barplot of c.140G>C samples is the same plot as shown in Fig. 2b. The BC31 and BC32 are albino (c.230G>T) and WT mice, respectively. The y-axis represents the percentage of DAJIN-reported alleles. The colours of the bar represent DAJIN-reported allele types. The horizontal lines in a bar represent the DAJIN-reported alleles. **b** Comparison to NGS in BC20. **c** Sanger verification of DAJIN detected alleles. The arrowheads represent the junction sites of DNA. **d** Comparison to NGS in BC25. **e** Comparison to NGS in BC26.

Next, we genotyped the PM mice using next-generation sequencing (NGS) and compared them to DAJIN’s results. NGS reported that a total of 16 mice (BC06, BC08, and BC12 in c.140G>C; BC14, BC15, and BC18 in c.316G>C; BC20, BC21, BC22, BC23, BC24, BC25, BC26, BC27, BC29, and BC30 in c.316G>C) had the intended PM (Fig. 3a). Notably, we found that the genotyping of NGS can be misleading when the samples contain SV alleles. For instance, DAJIN reported BC20 as a mosaic, including two SVs, one intended PM, and one mutated WT, and we confirmed the alleles via Sanger sequencing (Fig. 3b,c). In contrast, because NGS could not capture SVs, the genotyping result seemed heterozygous (Fig. 3c). Next, DAJIN showed BC25 included two SV alleles (∼70 bp insertion and large deletion). On the other hand, NGS showed one allele, which may be derived from the ∼70 bp inserted allele (Fig. 3d). Furthermore, as with BC20, DAJIN reported BC26 as a mosaic including SV alleles, while NGS reported it as heterozygous (Fig. 3e). These results indicate that although NGS can capture PM with high sensitivity and specificity, excluding SV alleles, including the intended PM by long-read sequencing is essential for accurate genotyping.

To compare DAJIN to existing long-read based SNV callers, we performed Medaka and Clair (35) on the 16 mice reported as PM positive by NGS. The results showed that both Medaka and Clair were prone to overlook the minor PM alleles (Additional file 1: Table. S9), which may be because they were designed to handle diploid genomes, while DAJIN can treat multiallelic mutations.

We next evaluated whether DAJIN correctly captured SV alleles. We conducted short and long PCRs for detecting small and large indel mutations (Additional file 2: Fig. S11a). The PCRs revealed 17 samples with aberrant PCR bands, which were consistent with the samples with SV alleles reported by DAJIN (Additional file 2: Fig. S11b,c). We further analysed BC02 and BC10, the alleles reported as ‘SV’ by DAJIN. Visualisation of the reads revealed that BC02 and BC10 had approximately 50 bp and 40 bp insertions, respectively (Additional file 2: Fig. S12a). We conducted PCR and validated that BC02 and BC10 had 50 bp and 40 bp larger bands, respectively, than that in WT control (Additional file 2: Fig. S12b,c). This result indicated that DAJIN was able to correctly annotate alleles with 40–50 bp insertion as ‘SV’ alleles. Taken together, these results indicated that DAJIN’s genotyping outperforms the conventional methods in accurately identifying the PMs owing to SV allele detection ability.

### DAJIN identifies knockout alleles

We next applied DAJIN to the knockout (KO) design. We designed to remove exon 6 of *Prdm14* by two-cut strategy with CRISPR-Cas9 system (25) (Fig. 4a). The predicted deletion size was 1043 bp length and may yield an inverted allele as a by-product. Thus, DAJIN annotated ‘WT’, ‘Deletion (Del)’, ‘Inversion’, and ‘SV’ alleles. We generated 10 *Prdm14* deletion founder mice (BC16–25) and analysed them using DAJIN with a WT mouse as a control (BC26); of the 10 mice, five (BC16, BC18, BC20, BC23, and BC24) contained ‘Mutated Del’ allele (Fig. 4b). Next, we evaluated BC18 and BC23 as DAJIN predicted that they contained the ‘Mutated Del’ allele. Read Visualisation showed that DAJIN discriminated ‘SV’ alleles with 100–200 bp larger deletion than the intended deletion (Fig. 4c). Furthermore, DAJIN’s consensus of ‘Mutated Del’ alleles showed that BC18 had a 1-bp deletion and BC23 included the 7-bp insertion and 1-bp substitution at the joint site, respectively. The same mutations were validated using Sanger sequencing (Fig. 4d,e).

**Figure 4.**
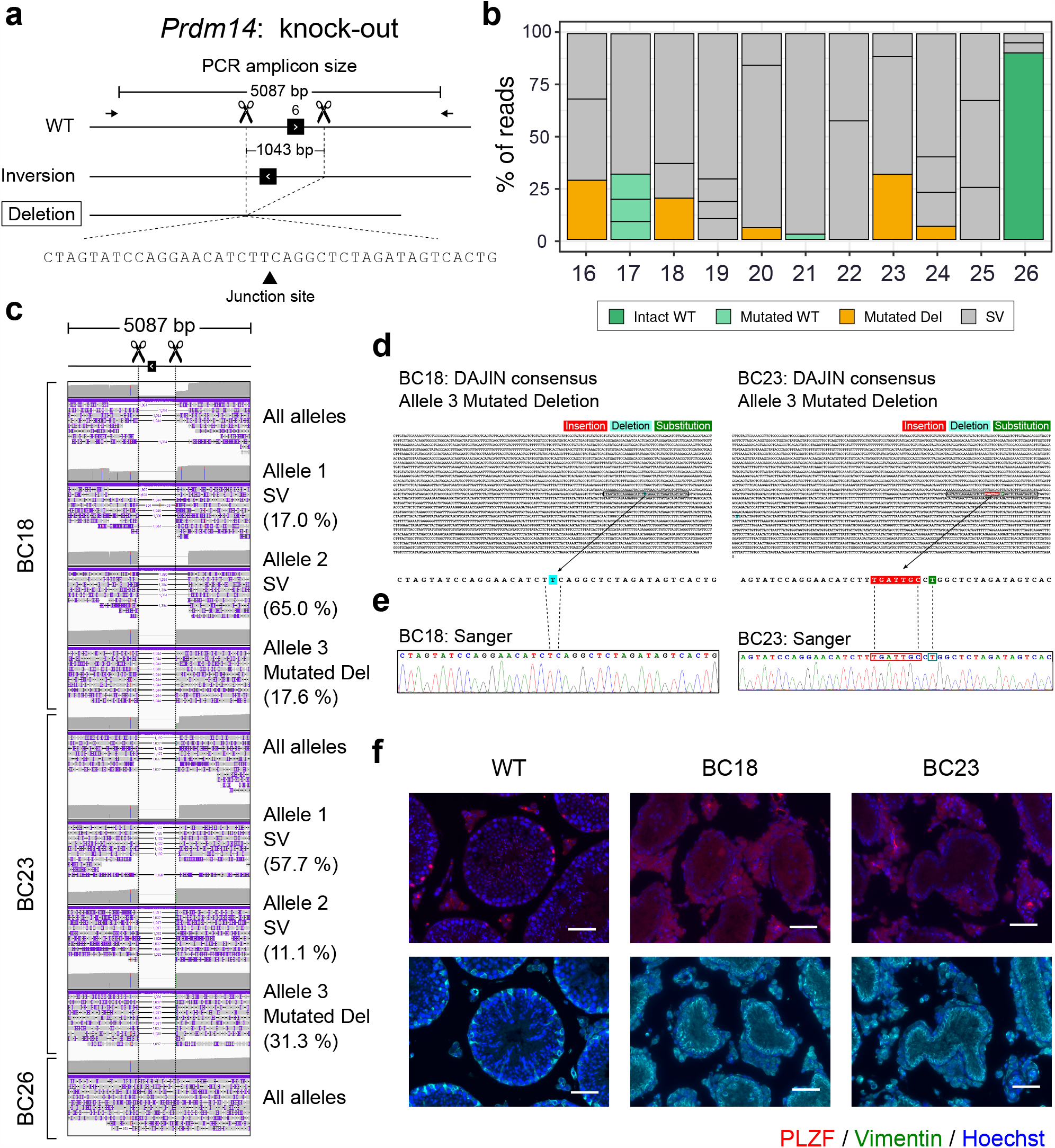
Application of DAJIN for knockout design. **a** Genome editing design for *Prdm14* knockout. The scissors and dotted lines represent Cas9-cutting sites. The arrows represent PCR primers, including the size of the PCR amplicon. The boxed allele type represents the target allele. The inversion allele represents a possible by-product. The triangle on the nucleotides represents a junction site of two DNA fragments. **b** DAJIN’s report of the allele percentage. The barcode numbers on the x-axis represent mouse IDs. BC26 is a WT control. The y-axis represents the percentage of DAJIN-reported alleles. The colours of the bar represent DAJIN-reported allele types. The compartments partitioned off by horizontal lines in a bar represent the DAJIN-reported alleles. **c** Visualisation of nanopore sequence reads at *Prdm14* target locus. BC18 and BC23 contain target alleles. BC26 is a WT control. The ‘All alleles’ track represents all reads of each sample. The ‘Allele’ track represents DAJIN-reported alleles. **d** DAJIN’s consensus sequences of the target allele. The top sequence represents the consensus sequence of target alleles of BC18 Allele 3 and BC23 Allele 3. The bottom sequences enlarge the boxed sequence of the consensus sequence. The colours on the nucleotides represent mutation types, including insertion (red), deletion (sky blue), and substitution (green). **e** Validation by Sanger sequencing. The dotted lines represent corresponding nucleotides between Sanger and DAJIN’s consensus sequences. **f** PLZF (red), Vimentin (green), and Hoechst (blue) staining of the testis section of WT (left), BC18 (middle), and BC23 (right). Upper panels show co-staining of PLZF (red) and Hoechst (blue). Lower panels show co-staining of Vimentin (green) and Hoechst (blue). PLZF and Vimentin are markers of undifferentiated spermatogonia and Sertoli cells, respectively, along the seminiferous tubules’ basal lamina. Scale bar: 100 µm

We evaluated the phenotypes of BC18 and BC23 mice. The deletion of *Prdm14* inhibits primordial germ cell differentiation and causes the complete depletion of germ cells in adult female and male mice (40). We performed immunostaining of testis sections for PLZF1 (spermatogonia marker) and Vimentin (Sertoli cell marker). Spermatogonia were not detected in BC18 and BC23 (Fig. 4f).

To confirm whether DAJIN is applicable to the CRISPR-Cas12a system (41), we generated *Prdm14* KO mice using Cas12a (Additional file 2: Fig. S13a). In all, 15 founder mice were obtained, and DAJIN analysis revealed that four of them (BC10, BC11, BC12, and BC13) had ‘Mutated Del’ (Additional file 2: Fig. S13b). We validated the mice carrying the deletion allele using conventional PCR and electrophoresis analysis. The electrophoresis analysis of the Cas9 group showed that five mice (BC16, BC18, BC20, BC23, and BC24) had the deletion alleles, similar to DAJIN’s report (Additional file 2: Fig. S13c,d).

To evaluate the versatility of DAJIN at other genomic loci and different cleavage widths, we established KO mice for the *Ddx4* gene using the two-cut strategy with Cas9 and Cas12a system. *Ddx4* KO was designed to cleave 3377 bp, including exons 11–15. We obtained 21 founder mice and analysed the 5221 bp PCR amplicon containing the target region (Additional file 2: Fig. S14a). DAJIN reported that one mouse (BC27) subjected to Cas12a-based genome editing and four mice (BC36, BC39, BC44, and BC46) subjected to Cas9-based genome editing carried the ‘Mutated Del’ allele (Additional file 2: Fig. S14b). The presence of the deletion alleles was confirmed via electrophoresis of the PCR products (Additional file 2: Fig. S14c,d). Furthermore, we generated KO mice for the *Stx2* gene using the two-cut strategy to cleave 727 bp, including exon 5 of *Stx2* (42). DAJIN reported that 13 of the 29 founder mice (BC01, BC03, BC04, BC05, BC07, BC09, BC14, BC15, BC20, BC21, BC22, BC23, and BC24) had the ‘Mutated Del’ allele (Additional file 2: Fig. S15a,b), and DAJIN’s results were consistent with the PCR-based genotyping (Additional file 2: Fig. S15c,d). In the *Stx2* analysis, DAJIN detected the ‘Inversion’ allele in three mice (BC08, BC16, and BC17). To verify the inversion alleles, we performed PCR for amplifying the genome region containing the inversion junction sites (Additional file 2: Fig. S15e). The inversion band was found in all three mice (Additional file 2: Fig. S15f). Besides, the 1 bp (A) insertion at the inversion junction site was found in DAJIN’s consensus sequence of BC17. This insertion was also confirmed via Sanger sequencing (Additional file 2: Fig. S15g). These results indicated that DAJIN could accurately identify single-nucleotide variants in inversion alleles. Next, DAJIN reported three SVs in BC25 (Additional file 2: Fig. S15b). The PCR electrophoresis validated the three alleles (Additional file 2: Fig. S16a,b). Moreover, DAJIN’s consensus sequence of BC25 reported that Allele 1, 2, and 3 included 986 bp deletion, 2477 bp deletion plus 1 bp substitution, and 1345 bp deletion, respectively (Additional file 3). Then we performed Sanger sequencing, and the results perfectly matched with that of DAJIN’s report at a single-nucleotide resolution (Additional file 2: Fig. S16c). Lastly, to compare DAJIN’s SV detection ability to the previous SV callers, we tested NanoSV and Sniffles for the alleles of BC25. Although NanoSV and Sniffles annotated SVs, they were not able to discriminate the three alleles correctly (Additional file 2: Fig. S17). We also performed NanoSV and Sniffles on BC18 and BC23 in order to test whether they can detect KO alleles. They captured the one cutting site (chr1:13118480) but could not differentiate the intended deletion and SV alleles with 100–200 bp larger deletion than the intended deletion (Additional file 4). Taken together, these results demonstrated DAJIN’s ability to accurately genotype the KO design. DAJIN’s genotyping for KO alleles was perfectly matched with Sanger sequencing. Furthermore, it correctly identifies multiallelic SVs, which current tools could not.

### DAJIN identifies flox knock-in alleles

Cre-LoxP-based conditional KO experiments are mostly performed to analyse gene function under specific conditions. Genome editing for generating floxed alleles requires *cis* knock-in at two loci simultaneously, which lowers the generation efficiency. Moreover, genotyping of the *cis* knock-in is difficult and error-prone owing to the need to identify *cis* mutations at several kilobases of the DNA region. Besides, the generation of floxed alleles using ssODNs as the donor of the knock-in sequence occasionally leads to the introduction of unintended mutations in a critical LoxP sequence because of the error in the synthesis process and its secondary structure (43). Because of these difficulties, no standard genotyping method is currently available to comprehensively and accurately evaluate flox mutations induced by genome editing in one step. Therefore, we evaluated whether DAJIN can correctly genotype floxed alleles at single-nucleotide resolution.

We performed validation experiments using plasmid vectors with completely defined sequences. We generated six types of plasmids with LoxP sequences: 1) ‘Intended flox’, 2) 1-bp insertion in left LoxP, 3) 1-bp deletion in left LoxP, 4) 1-bp substitution in left LoxP, 5) 1-bp substitution in right LoxP, and 6) 1-bp substitution in both LoxPs (Fig. 5a). We mixed the WT genomic DNA with each plasmid to imitate the heterozygous genotype. The mixed DNA samples were used as a PCR template. Then DAJIN analysed the PCR products that were 2724 bp in length (Fig. 5b). The results showed that DAJIN correctly discriminated between ‘WT’, ‘Intended flox’, and ‘Mutated flox’. Furthermore, the proportion of WT and LoxP alleles reflected the designed allele frequency (Fig. 5c). On the other hand,

**Figure 5.**
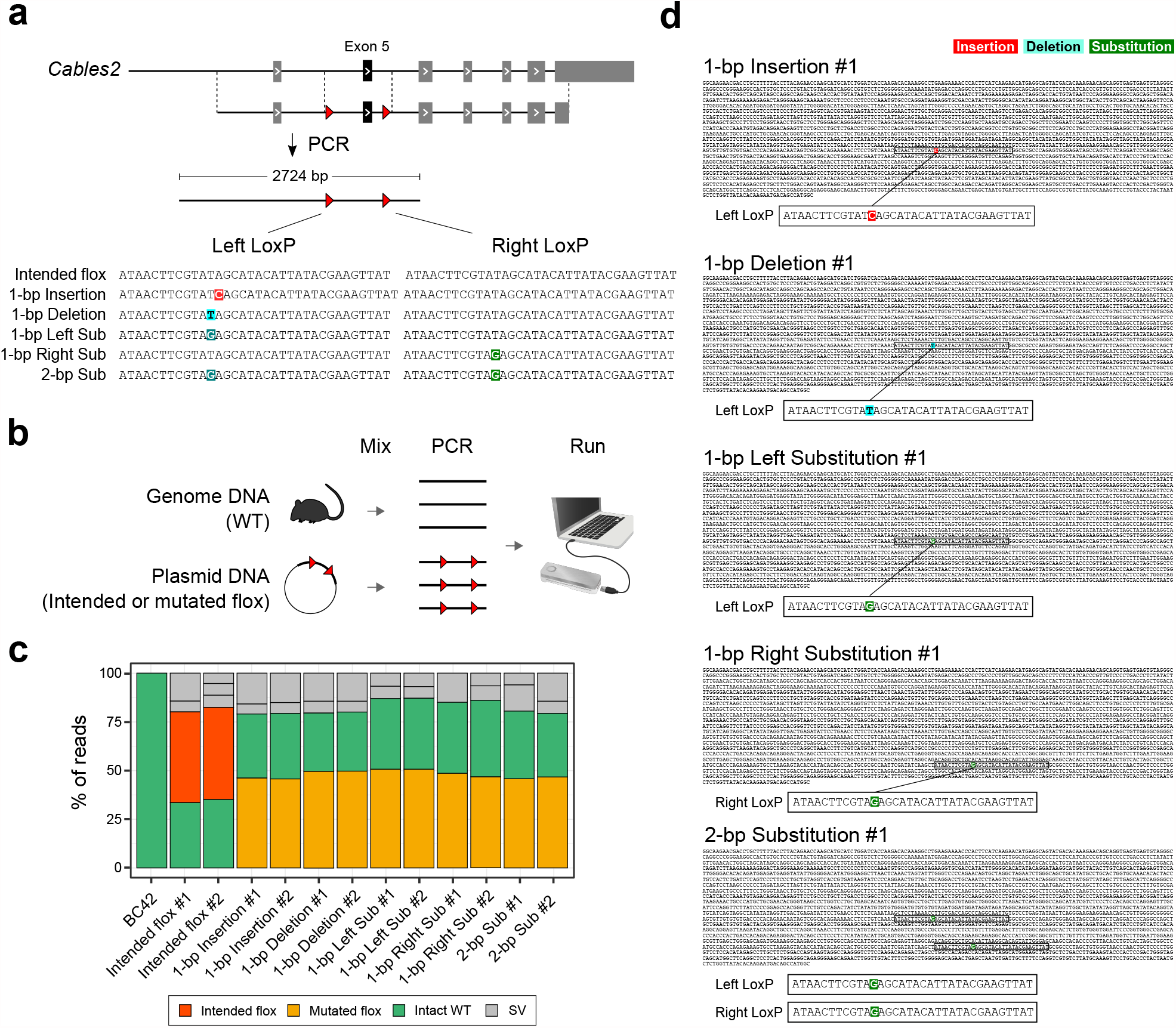
Precise SNV detection in flox knock-in allele. **a** Genome editing design. The red arrowheads represent LoxPs. The colours on the nucleotides represent the types of mutations, including insertion (red), deletion (sky blue), and substitution (green). ‘Sub’: substitution. **b** Experimental design. **c** DAJIN’s report of the allele percentage. The barcode numbers on the x-axis represent sample IDs. Barcode42 is a WT control. The y-axis represents the percentage of DAJIN-reported alleles. The colours of the bar represent DAJIN-reported allele types. The compartments partitioned off by horizontal lines in a bar represent the DAJIN-reported alleles. ‘Sub’: substitution. **d** DAJIN’s consensus sequences of a floxed allele in each sample. The colours on the nucleotides represent mutation types, including insertion (red), deletion (sky blue), and substitution (green). The boxed sequences in the consensus sequences are LoxP sites.

DAJIN reported there were about 20% of SV alleles in every sample. Thus we examined the SV alleles and found the SV alleles included either the left-side LoxP or the right-side LoxP (Additional file 2: Fig. S18). These ‘pseudo-LoxP’ alleles could be induced by PCR recombination. Next, DAJIN’s consensus sequences discovered all types of variants that we induced in the LoxPs (Fig. 5d). These results indicated that DAJIN could discriminate knock-in sequences according to their variants.

We next assessed DAJIN’s genotyping performance using genome-edited flox KI mice. We targeted *Cables2* exon 5, and to induce it, we simultaneously cut introns 5 and 6 and knocked in the two LoxPs via homology-directed repair using a single-strand plasmid DNA donor. In this design, genome editing potentially generates seven types of alleles, such as WT, flox, Left LoxP, Right LoxP, Inversion, Deletion, and Unexpected SV alleles (Fig. 6a). We obtained 20 founder mice (BC01–BC20) and DAJIN reported that 9 mice (BC06, BC10, BC11, BC12, BC13, BC14, BC17, BC18, and BC20) contained the ‘Intended flox’ allele, and 11 mice (BC01, BC05, BC06, BC09, BC11, BC12, BC13, BC17, BC18, BC19, and BC20) contained ‘Deletion’ alleles (Fig. 6b). Since the *Asc*I or *Eco*RV recognition sites were knocked in next to the LoxP sequence, PCR-RFLP digestion of *Asc*I or *Eco*RV can reveal LoxP insertion (Fig. 6c). The PCR-RFLP results were consistent with DAJIN (Fig. 6d). We also evaluated the ‘Deletion’ alleles using standard PCR (Fig. 6e), and the results were compatible with DAJIN’s reports (Fig. 6f). Moreover, the consensus sequence of DAJIN for BC14 showed that the entire 2724 bp was intact, including the left and right LoxP sites (Fig. 6g). Sanger sequencing also revealed that both left and right knock-in sequences were intact, corresponding to DAJIN’s consensus sequence (Fig. 6h). These results indicated that DAJIN correctly identified the intended floxed mice.

**Figure 6.**
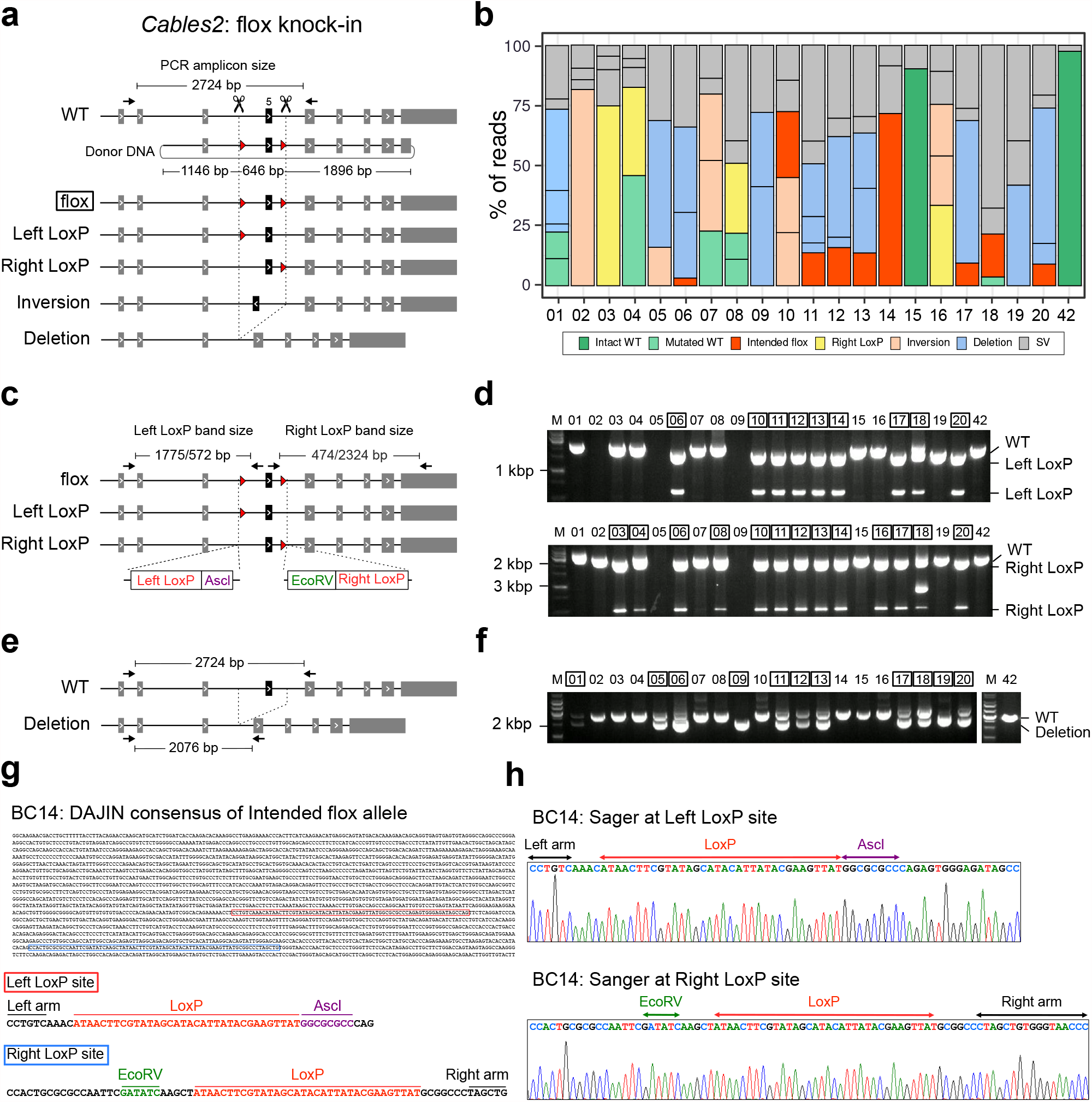
Application of DAJIN for flox knock-in design. **a** Genome editing design for flox knock-in into the *Cables2* locus. The scissors represent Cas9-cutting sites. The arrows represent PCR primers, including the size of the PCR amplicon. The circular DNA represents the donor DNA. The base numbers on the donor DNA describe the left, central, and right arm sizes. The red arrowheads represent LoxPs. The boxed allele type represents the target allele. The other allele types include Left LoxP and Right LoxP. Inversion and Deletion represent possible by-products. **b** DAJIN’s report of the allele percentage. The barcode numbers on the x-axis represent mouse IDs. BC42 is a WT control. The y-axis represents the percentage of DAJIN-reported alleles. The colours of the bar represent DAJIN-reported allele types. The compartments partitioned off by horizontal lines in a bar represent the DAJIN-reported alleles. **c** Design of PCR-RFLP to validate LoxP knock-in alleles. The *Asc*I and *Eco*RV restriction sites are adjacent to Left LoxP and Right LoxP, respectively. The arrows represent PCR primers for the digested DNA fragments, including PCR product sizes. **d** PCR results for the detection of the LoxP knock-in allele. The top and bottom panels represent DNA fragments digested with *Asc*I and *Eco*RV, respectively. The numbers on the panel mean barcode IDs. The boxed number represents the samples with LoxP alleles. **e** Design of PCR to validate deletion alleles. The arrows represent PCR primers. **f** PCR results for the detection of deletion alleles. The panels show the PCR products. The number on the panel means barcode IDs. The boxed number represents the samples with deletion alleles. **g** DAJIN’s consensus sequence of the floxed allele in BC14. The red and blue boxes represent left and right LoxP sites, respectively. **h** Sanger sequences for the LoxP sites.

To confirm whether the next generation inherits the mutations, the BC11, BC12, BC13, and BC14, having an ‘Intended flox’ allele were mated with WT. (Additional file 2: Fig. S19). The results showed that the genotype of the first filial generation (F1) mice from BC14 were heterozygous flox/WT, suggesting that BC14 has homozygous floxed alleles in the germline. Moreover, F1 mice from BC11, BC12, BC13, and BC18 had the ‘Intended flox’ allele, which corresponded with DAJIN’s results. Therefore, it provides evidence that DAJIN accurately captured the genotypes of the founder mice. DAJIN detected the ‘Inversion’ allele in five mice (BC02, BC05, BC07, BC10, and BC16; Fig. 6b). To verify the inversion allele, we performed PCR to amplify the genomic region including the inversion junction site. The results revealed the inversion band in the same five samples corresponding to those mentioned in DAJIN’s report (Additional file 2: Fig. S20a,b). Furthermore, the consensus sequence of BC02 revealed a 1-bp substitution at the inversion junction site. Sanger sequencing also detected the same substitution (Additional file 2: Fig. S20c), which suggested that DAJIN can detect complex alleles such as an inversion with single-nucleotide variants.

To confirm that DAJIN is also useful for flox analysis at other loci, we further generated and analysed floxed mice for two genes, *Exoc7* and *Usp46*. In the *Exoc7* project (Additional file 2: Fig. S21a), we obtained 40 founder mice and analysed them using DAJIN. DAJIN identified 11 mice with the ‘Intended flox’ allele, 7 with the ‘Left LoxP’ allele, 16 with the ‘Right LoxP’ allele; and 5 with the ‘Deletion’ allele (Additional file 2: Fig. S21b). To verify DAJIN’s results, we performed PCR-RFLP and standard PCR to detect the LoxP and deletion band, respectively (Additional file 2: Fig. S22a). The PCR-RFLP results agreed with the DAJIN report except for BC13 and BC32, which was shown to have no ‘Left LoxP’ allele by DAJIN, but PCR-RFLP detected this allele (Additional file 2: Fig. S22b). Because the majority of reads in BC13 and BC32 were annotated as ‘Deletion’ allele (Additional file 2: Fig. S21b), the deletion band might be predominantly amplified, and the number of ‘Left LoxP’ reads decreased owing to the PCR bias. In deletion alleles, the PCR genotyping was in agreement with DAJIN’s results (Additional file 2: Fig. S22c,d). Next, to confirm whether the allele determined by using DAJIN was inherited through the next generation, we mated WT with *Exoc7* BC14 that DAJIN reported as heterozygous for ‘Intended flox’ allele (42.4%) and ‘Right LoxP’ allele (45.5%; Additional file 2: Fig. S21b). Of the total 11 F1 mice, 5 were flox/WT and 6 were Right LoxP/WT mice (Additional file 2: Fig. S23), which represented the accuracy of DAJIN’s genotyping.

In the *Usp46* project (Additional file 2: Fig. S24a), we obtained 34 founder mice, and DAJIN reported 4 mice with the ‘Intended flox’ allele, 2 with the ‘Left LoxP’ allele, 2 with the ‘Right LoxP’ allele, and 21 with the ‘Deletion’ allele (Additional file 2: Fig. S24b). DAJIN’s results were validated using PCR-RFLP that detected Left and Right LoxP alleles (Additional file 2: Fig. S25a,b). However, some results were inconsistent. First, PCR-RFLP analysis of BC23 and BC33 indicated that these mice may have flox alleles, but DAJIN did not report it. Second, PCR-RFLP identified ‘Left LoxP’ alleles in BC13 and BC27, but DAJIN did not. Since DAJIN reported that these samples dominantly had the ‘Deletion’ alleles (Additional file 2: Fig. S24b), the mismatch might be caused by PCR bias. In contrast, DAJIN detected the ‘Left LoxP’ allele in BC21, but PCR-RLFP did not. Thus, we conducted PCR again by adjusting the dilution ratio and detected the left LoxP band (Additional file 2: Fig. S25c). For other alleles such as ‘Right LoxP’ and ‘Deletion’ alleles, DAJIN and PCR-RFLP’s genotyping results were consistent (Additional file 2: Fig. S25b,d,e). Notably, the PCR band in six samples (BC07, BC12, BC17, BC23, BC30, and BC33) seemed to be a deletion band, whereas DAJIN reported them as ‘SV’ alleles. Visualisation of the reads revealed that the alleles contained about 30–200 bp indels (Additional file 2: Fig. S26). The result indicated that DAJIN’s annotation is accurate even when distinguishing allele types by PCR band size was difficult. Finally, we investigated the next generation of mice. We obtained F1 progeny by crossing BC10 and BC11 with WT and found floxed and deletion alleles in the F1 mice (Additional file 2: Fig. S27a), which suggested DAJIN’s allele reports of BC10 and BC11 were accurate. Next, to tackle the ‘pseudo-LoxP’ alleles, we implemented DAJIN to detect potential pseudo-LoxP alleles, and DAJIN annotated the flox allele of BC04 as a pseudo-flox allele (Additional file 2: Fig. S27b). Then we evaluated genotype of BC04 in the F1 mice. The results revealed that the genotype of BC04 was not flox but Left-LoxP/Right-LoxP (Additional file 2: Fig. S27b), which indicates BC04 had a pseudo-flox. Lastly, we performed NanoSV and Sniffles to test whether they can detect flox alleles. NanoSV captured the insertion, but it did not report the LoxP sequences. On the other hand, Sniffles could not detect LoxP alleles (Additional file 4). Taken together, these results provide evidence that DAJIN can accurately and comprehensively detect diverse mutations of floxed mice.

## DISCUSSION

Conventional approaches such as short-range PCR, Sanger sequencing, and PCR-RFLP are standard methods to detect on-target mutagenesis induced by CRISPR-Cas and other genome editing tools. Recent studies on on-target variability of edited materials clearly show that the characterisation of genome editing events and selection of animals or cultured cells with intended and unintended mutations require alternative methods with higher sensitivity and broader range to capture mosaic mutagenic events (44,45). In this study, we developed a genotyping method using a novel software, DAJIN, that can be applied for long-read sequencing to validate the quality of genome-edited organisms. Our approach involving DAJIN can handle multiple samples obtained under different editing conditions and identify not only alleles with the desired mutation but also those with a variety of unexpected mutations, including large deletions, in a single run; this refined strategy can contribute to a more precise genome editing.

One of DAJIN’s distinguishing features is its automatic allele clustering and annotation, as well as the utilisation of a long-read sequencer. Genotyping tools similar to DAJIN have been developed previously (46-48); however, they are optimised for short-read sequencing. Because DAJIN uses a long-read sequencer, it can identify cis- or trans-heterozygosity and complex mutant alleles such as unexpected indels and SVs. Besides, although several tools have been developed to detect PMs or SVs using long-read sequencing, DAJIN outperformed them in capturing mutations (Additional file 1: Table. S9, Additional file 2: Fig. S17). It may be because the previous method focuses on monoploid or diploid organisms; however, since genome-edited samples often contain more than 2 alleles, these unpredictable allele numbers might make it challenging for the previous tools to capture mutations correctly. Polyploid phasing, which allows the reconstruction of haplotypes of the polyploid genome, is similar to DAJIN in terms of estimating alleles. WHATSHAP POLYPHASE (49) and H-PoPG (50) are state-of-the-art tools for polyploid phasing, but these tools require prior knowledge of the polyploidy of the target organism. Thus, the current tools for genotyping of genome-edited samples have some limitations.

As long-read sequencing induces base calling errors across a segment and cannot be used as is to validate the genome editing outcomes (22), novel screening techniques and tools need to be developed in order to identify diverse sequence changes in the genome. Short-range PCR amplification and Sanger sequencing confirmed that no additional mutation was detected in ‘Intact’ alleles identified by DAJIN, suggesting that DAJIN validates the consequences of genome editing at the base level (Figs. 2 and 4). We investigated DAJIN’s accurate genotyping in three major genome editing designs, including PM, two-cut KO, and flox KI, by comparing conventional methods such as PCR, RFLP, Sanger sequencing, and NGS. In most cases, DAJIN’s genotyping results were consistent with those obtained using the conventional methods. With regard to PM, DAJIN reported the intended PM plus SV alleles (BC25, BC26), whereas NGS showed only the intended PM allele (Fig 3). For the two-cut KO designs, DAJIN’s genotyping results were in complete agreement with the PCR-based genotyping. In the flox design, DAJIN correctly identified intended flox KI in most cases; however, there were three false negative samples that were caused by PCR amplification bias (discussed in the last paragraph).

DNA double-strand break repair leads to long-range deletion, inversion, and insertion, as well as small indels in zygotes and stem cells (12,13,51-53). In previous knock-in experiments, exogenous repair templates and unwanted mismatches had been identified around the target region (20,54-56). A parallel analysis of short- and long-read sequencing results confirmed that DAJIN was able to identify editing outcomes, including unpredictable large-scale inversion events, in mouse zygotes (Additional file 2: Figs. S15 and S20). Short-range analysis combined with NGS could not detect SV alleles and therefore gave misleading results of founder genotype as a mosaic without SV alleles (Fig. 3b,c). In some cases, it recognised an SV allele as an allele with the intended mutation (Fig. 3d). In addition, it reported separate loxP insertions in *cis* generated using two gRNAs positioned up to 2 kb apart on the same chromosome (Figs. 4 and 5, Additional file 2: Figs. S21 and S24). Detection of mutations and/or integrations in *cis/trans* at a kilobase-scale distance requires a combination of assays and a considerable amount of time and effort. Recently, DNA cleavage in cultured cells and zygotes was shown to induce gene conversions mediated by homologous chromosomes or homologous sequences on the same chromosome (14,15). Genotype assessment using DAJIN facilitates the selection of genome-edited samples with precisely targeted alleles or those with unwanted alleles. DAJIN might contribute to a better understanding of the consequences of editing events at the targeted locus.

Comprehensive mutation analysis might reduce the overall cost of genome editing in not only laboratory mice but also other experimental animals with a higher cost of maintenance or farm animals with a longer generation time. DAJIN is also preeminent in multi-specimen processing due to its PCR-based barcoding, which enables multiplexed sequencing and allows sufficient coverage of numerous samples in a single run (Additional file 1: Table. S5). In this study, BC01–35 of Usp46 shared the same barcode as that of BC01–26 of Prdm14 and BC27–35 of Ddx4, and we analysed 83 samples from three different mouse strains in a single run. It can be undoubtedly applied to samples with a larger number of strains but a smaller number of mice for each strain. Besides, because DAJIN supports parallel processing, we were able to analyse 226 samples (total 5,982,507 reads) in only 15 h using a general-purpose desktop computer (Additional file 1: Table. S10). Thus, DAJIN’s genotyping is considerably time-efficient compared to that of conventional genotyping methods. Multiple alleles may be generated in the edited cell culture pools, but they cannot be segregated as in the case of founder animals. Our results indicate that genotyping with DAJIN can also be applied to the assessment of editing outcomes in cellular experiments (*e*.*g*. CRISPR screening) and cellular therapies. Thus, DAJIN offers a novel strategy to identify multiple genomic changes, including large sequence alterations or unexpected mutations, regardless of the species or type of the material.

DAJIN’s current limitations are false negatives of flox KI samples and pseudo-flox due to PCR. Although PCR enables convenient barcoding and high-level enrichment of target genomic locus, it may cause several issues. The first is PCR bias, a lower efficiency to amplify long reads and GC-rich sequences, known as length bias and GC bias. GC bias can be alleviated using high-grade DNA polymerase, but length bias cannot be removed, affecting the accuracy of DAJIN’s allele percentage. In the *Exoc7* and *Usp46* flox knock-in, the percentage of the ‘Intended flox’ allele was low because the deletion allele might have been preferentially amplified, and three false negatives (*Exoc7* BC13 and *Usp46* BC23, 33) were reported (Additional file 2: Fig. S21 and S24). Next, the ‘pseudo-LoxP’ alleles could be generated if the PCR products, which included one-side LoxP but not another-side LoxP, worked as a PCR primer to anneal WT allele in the next PCR step. In this study, we found the *Usp46* BC04 included pseudo-flox alleles (Additional file2: Fig. S27). Since the samples with the pseudo-flox must be excluded, DAJIN flags samples that may be pseudo-flox. Two recently developed methods may address these issues. First, IDMseq is used for labelling PCR amplicons using unique molecular identifiers, which eliminates PCR bias and allows more quantitative analysis of allele frequencies (57). Second, nCATS enables the enrichment of the genome region without PCR; thereby, it may avoid pseudo-flox (58). Notably, DAJIN can be used for the nanopore sequencing reads from these techniques; thus, combining these techniques with DAJIN can potentially overcome the issues caused by PCR.

## DATA AVAILABILITY

DAJIN is accessible at https://github.com/akikuno/DAJIN under the MIT Licence. The version of DAJIN used in this study to reproduce the analyses can be found at https://github.com/akikuno/DAJIN/tree/manuscript-version. All sequencing data are available in the DDBJ DRA under accession number DRA011971.

## ACKNOWLEDGEMENTS

We would like to thank the staff at the Laboratory Animal Resource Center University of Tsukuba for their help in breeding and rearing of the mice. We are grateful to Tomoyuki Fujiyama for advice on the experimental design. We would also like to thank Ozaki Haruka for fruitful discussions.

## AUTHOR CONTRIBUTIONS

S. A., A. Y., S. T., F. S., and S. M. designed the study with inputs from all other authors. A. K., K. S., and S. S. conducted the bioinformatics analyses. Y. I., K. K., Y. H., T. M., and M. M. conducted molecular experiments. Ke. M., A. W., N. M., T. T., H. D., Ka. M., and M. H. conducted lineage and phenotypic analysis experiments on mice. S. A., M. I., N. I., Y. D., and Y. T. conducted experiments to produce mice. A. K., Y. I., S. A., K. S., and S. M. wrote the manuscript. The authors read and approved the final manuscript.

## Supplementary Information

**Additional file 1**.

Tables S1–S9 (MS Excel).

**Additional file 2**.

Supplementary Figures S1–S21 with their legends (MS Word).

**Additional file 3**.

DAJIN’s consensus sequences (HTML).

**Additional file 4**.

VCF files by NanoSV and Sniffles for Prdm14 and Cables2

## REFERENCES

1. Christian, M., Cermak, T., Doyle, E.L., Schmidt, C., Zhang, F., Hummel, A., Bogdanove, A.J. and Voytas, D.F. (2010) Targeting DNA double-strand breaks with TAL effector nucleases. Genetics, 186, 757–761.

2. Jinek, M., Chylinski, K., Fonfara, I., Hauer, M., Doudna, J.A. and Charpentier, E. (2012) A programmable dual-RNA-guided DNA endonuclease in adaptive bacterial immunity. Science, 337, 816–821.

3. Komor, A.C., Kim, Y.B., Packer, M.S., Zuris, J.A. and Liu, D.R. (2016) Programmable editing of a target base in genomic DNA without double-stranded DNA cleavage. Nature, 533, 420–424.

4. Yeh, C.D., Richardson, C.D. and Corn, J.E. (2019) Advances in genome editing through control of DNA repair pathways. Nat Cell Biol, 21, 1468–1478.

5. Mizuno, S., Dinh, T.T., Kato, K., Mizuno-Iijima, S., Tanimoto, Y., Daitoku, Y., Hoshino, Y., Ikawa, M., Takahashi, S., Sugiyama, F. et al.. (2014) Simple generation of albino C57BL/6J mice with G291T mutation in the tyrosinase gene by the CRISPR/Cas9 system. Mamm Genome, 25, 327–334.

6. Yin, H., Xue, W., Chen, S., Bogorad, R.L., Benedetti, E., Grompe, M., Koteliansky, V., Sharp, P.A., Jacks, T. and Anderson, D.G. (2014) Genome editing with Cas9 in adult mice corrects a disease mutation and phenotype. Nat Biotechnol, 32, 551–553.

7. Smith, C., Gore, A., Yan, W., Abalde-Atristain, L., Li, Z., He, C., Wang, Y., Brodsky, R.A., Zhang, K., Cheng, L. et al.. (2014) Whole-genome sequencing analysis reveals high specificity of CRISPR/Cas9 and TALEN-based genome editing in human iPSCs. Cell Stem Cell, 15, 12–13.

8. Teboul, L., Herault, Y., Wells, S., Qasim, W. and Pavlovic, G. (2020) Variability in Genome Editing Outcomes: Challenges for Research Reproducibility and Clinical Safety. Mol Ther, 28, 1422–1431.

9. Canver, M.C., Bauer, D.E., Dass, A., Yien, Y.Y., Chung, J., Masuda, T., Maeda, T., Paw, B.H. and Orkin, S.H. (2014) Characterization of genomic deletion efficiency mediated by clustered regularly interspaced short palindromic repeats (CRISPR)/Cas9 nuclease system in mammalian cells. J Biol Chem, 289, 21312–21324.

10. Hendel, A., Kildebeck, E.J., Fine, E.J., Clark, J., Punjya, N., Sebastiano, V., Bao, G. and Porteus, M.H. (2014) Quantifying genome-editing outcomes at endogenous loci with SMRT sequencing. Cell Rep, 7, 293–305.

11. Kraft, K., Geuer, S., Will, A.J., Chan, W.L., Paliou, C., Borschiwer, M., Harabula, I., Wittler, L., Franke, M., Ibrahim, D.M. et al.. (2015) Deletions, Inversions, Duplications: Engineering of structural variations using CRISPR/Cas in Mice. Cell Rep, 10, 833–839.

12. Boroviak, K., Fu, B., Yang, F., Doe, B. and Bradley, A. (2017) Revealing hidden complexities of genomic rearrangements generated with Cas9. Sci Rep, 7, 12867.

13. Kosicki, M., Tomberg, K. and Bradley, A. (2018) Repair of double-strand breaks induced by CRISPR-Cas9 leads to large deletions and complex rearrangements. Nat Biotechnol, 36, 765–771.

14. Ma, H., Marti-Gutierrez, N., Park, S.W., Wu, J., Lee, Y., Suzuki, K., Koski, A., Ji, D., Hayama, T., Ahmed, R. et al.. (2017) Correction of a pathogenic gene mutation in human embryos. Nature, 548, 413–419.

15. Javidi-Parsijani, P., Lyu, P., Makani, V., Sarhan, W.M., Yoo, K.W., El-Korashi, L., Atala, A. and Lu, B. (2020) CRISPR/Cas9 increases mitotic gene conversion in human cells. Gene Ther, 27, 281–296.

16. Liang, D., Marti, N.G., Chen, T., Lee Y., Park, S.W., Ma, H. et al.. Frequent gene conversion in human embryos induced by double strand breaks. bioRxiv, 162214. doi: https://doi.org/10.1101/2020.06.19.162214 (2020).

17. Simeonov, D.R., Brandt, A.J., Chan, A.Y., Cortez, J.T., Li, Z., Woo, J.M., Lee, Y., Carvalho, C.M.B., Indart, A.C., Roth, T.L. et al.. (2019) A large CRISPR-induced bystander mutation causes immune dysregulation. Commun Biol, 2, 70.

18. Brinkman, E.K., Chen, T., Amendola, M. and van Steensel, B. (2014) Easy quantitative assessment of genome editing by sequence trace decomposition. Nucleic Acids Res, 42, e168.

19. Birling, M.C., Schaeffer, L., Andre, P., Lindner, L., Marechal, D., Ayadi, A., Sorg, T., Pavlovic, G. and Herault, Y. (2017) Efficient and rapid generation of large genomic variants in rats and mice using CRISMERE. Sci Rep, 7, 43331.

20. Lanza, D.G., Gaspero, A., Lorenzo, I., Liao, L., Zheng, P., Wang, Y., Deng, Y., Cheng, C., Zhang, C., Seavitt, J.R. et al.. (2018) Comparative analysis of single-stranded DNA donors to generate conditional null mouse alleles. BMC Biol, 16, 69.

21. Canaj, H., Hussmann, A.J., Li, H., Beckman, A.K., Goodrich, L., Cho, H.N. et al.. (2019) Deep profiling reveals substantial heterogeneity of integration outcomes in CRISPR knock-in experiments. bioRxiv, 841098; doi: https://doi.org/10.1101/841098.

22. McCabe, V.C., Codner, F.G., Allan, J.A., Caulder, A., Christou, S., Loeffler, J. et al.. (2019) Application of long-read sequencing for robust identification of correct alleles in genome edited animals. bioRxiv, 838193doi: https://doi.org/10.1101/838193.

23. Kaneko, T. and Mashimo, T. (2015) Simple Genome Editing of Rodent Intact Embryos by Electroporation. PLoS One, 10, e0142755.

24. Sato, Y., Tsukaguchi, H., Morita, H., Higasa, K., Tran, M.T.N., Hamada, M., Usui, T., Morito, N., Horita, S., Hayashi, T. et al.. (2018) A mutation in transcription factor MAFB causes Focal Segmental Glomerulosclerosis with Duane Retraction Syndrome. Kidney Int, 94, 396–407.

25. Mizuno-Iijima, S., Ayabe, S., Kato, K., Matoba, S., Ikeda, Y., Dinh, T.T.H., Le, H.T., Suzuki, H., Nakashima, K., Hasegawa, Y. et al.. (2020) Efficient production of large deletion and gene fragment knock-in mice mediated by genome editing with Cas9-mouse Cdt1 in mouse zygotes. Methods.

26. Dobin, A., Davis, C.A., Schlesinger, F., Drenkow, J., Zaleski, C., Jha, S., Batut, P., Chaisson, M., Gingeras, R.T. (2013) STAR: ultrafast universal RNA-seq aligner. Bioinformatics, 29, 15–21.

27. Robinson, J.T., Thorvaldsdottir, H., Winckler, W., Guttman, M., Lander, E.S., Getz, G. and Mesirov, J.P. (2011) Integrative genomics viewer. Nat Biotechnol, 29, 24–26.

28. Yang, C., Chu, J., Warren, R.L. and Birol, I. (2017) NanoSim: nanopore sequence read simulator based on statistical characterisation. Gigascience, 6, 1–6.

29. Li, H. (2018) Minimap2: pairwise alignment for nucleotide sequences. Bioinformatics, 34, 3094–3100.

30. Breunig, M.M., Kriegel, H.P., Ng, T.R., Sander, J. (2000) LOF: Identifying Density-Based Local Outliers. ACM SIGMOD Record, 29, 93–104, https://doi.org/10.1145/335191.335388.

31. McInnes, L., Healy, J., Astels, S. (2017) hdbscan: Hierarchical density based clustering. Journal of Open Source Software, 2, 205, doi:10.21105/joss.00205.

32. Li, H., Handsaker, B., Wysoker, A., Fennell, T., Ruan, J., Homer, N., Marth, G., Abecasis, G., Durbin, R. and Genome Project Data Processing, S. (2009) The Sequence Alignment/Map format and SAMtools. Bioinformatics, 25, 2078–2079.

33. Karolchik, D., Hinrichs, A.S., Furey, T.S., Roskin, K.M., Sugnet, C.W., Haussler, D. and Kent, W.J. (2004) The UCSC Table Browser data retrieval tool. Nucleic Acids Res, 32, D493–496.

34. https://github.com/nanoporetech/medaka

35. Luo, R., Wong, CL., Wong, YS., Tang, CI., Liu, CM., Leung, CM., Lam, TW. (2020) Exploring the limit of using a deep neural network on pileup data for germline variant calling. Nat Mach Intell, 2, 220–227

36. Cretu Stancu, M., van Roosmalen, M.J., Renkens, I., Nieboer, M.M., Middelkamp, S., de Ligt, J., Pregno, G., Giachino, D., Mandrile, G., Espejo Valle-Inclan, J. et al. (2017) Mapping and phasing of structural variation in patient genomes using nanopore sequencing. Nat Commun, 8, 1326.

37. Sedlazeck, F.J., Rescheneder, P., Smolka, M., Fang, H., Nattestad, M., von Haeseler, A. and Schatz, M.C. (2018) Accurate detection of complex structural variations using single-molecule sequencing. Nat Methods, 15, 461–468.

38. Gruning, B., Dale, R., Sjodin, A., Chapman, B.A., Rowe, J., Tomkins-Tinch, C.H., Valieris, R., Koster, J. and Bioconda, T. (2018) Bioconda: sustainable and comprehensive software distribution for the life sciences. Nat Methods, 15, 475–476.

39. McInnes, L., Healy, J., Melville, J. (2020) UMAP: Uniform Manifold Approximation and Projection for Dimension Reduction. 1802.03426v3.

40. Yamaji, M., Seki, Y., Kurimoto, K., Yabuta, Y., Yuasa, M., Shigeta, M., Yamanaka, K., Ohinata, Y. and Saitou, M. (2008) Critical function of Prdm14 for the establishment of the germ cell lineage in mice. Nat Genet, 40, 1016–1022.

41. Zetsche, B., Gootenberg, J.S., Abudayyeh, O.O., Slaymaker, I.M., Makarova, K.S., Essletzbichler, P., Volz, S.E., Joung, J., van der Oost, J., Regev, A. et al.. (2015) Cpf1 is a single RNA-guided endonuclease of a class 2 CRISPR-Cas system. Cell, 163, 759–771.

42. Osawa, Y., Usui, M., Kuba, Y., Le, T.H., Mikami, N., Nakagawa, T. et al.. (2020) EXOC1 regulates cell morphology of spermatogonia and spermatocytes in mice. bioRxiv, 139030, doi: https://doi.org/10.1101/2020.06.07.139030.

43. Gurumurthy, C.B., O’Brien, A.R., Quadros, R.M., Adams, J., Jr., Alcaide, P., Ayabe, S., Ballard, J., Batra, S.K., Beauchamp, M.C., Becker, K.A. et al.. (2019) Reproducibility of CRISPR- Cas9 methods for generation of conditional mouse alleles: a multi-center evaluation. Genome Biol, 20, 171.

44. Mianne, J., Codner, G.F., Caulder, A., Fell, R., Hutchison, M., King, R., Stewart, M.E., Wells, S. and Teboul, L. (2017) Analysing the outcome of CRISPR-aided genome editing in embryos: Screening, genotyping and quality control. Methods, 121-122, 68–76.

45. Burgio, G. and Teboul, L. (2020) Anticipating and Identifying Collateral Damage in Genome Editing. Trends Genet, 36, 905-914.

46. Park, J., Lim, K., Kim, J.S. and Bae, S. (2017) Cas-analyzer: an online tool for assessing genome editing results using NGS data. Bioinformatics, 33, 286–288.

47. Clement, K., Rees, H., Canver, M.C., Gehrke, J.M., Farouni, R., Hsu, J.Y., Cole, M.A., Liu, D.R., Joung, J.K., Bauer, D.E. et al.. (2019) CRISPResso2 provides accurate and rapid genome editing sequence analysis. Nat Biotechnol, 37, 224–226.

48. Iida, M., Suzuki, M., Sakane, Y., Nishide, H., Uchiyama, I., Yamamoto, T., Suzuki, K.T. and Fujii, S. (2020) A simple and practical workflow for genotyping of CRISPR-Cas9-based knockout phenotypes using multiplexed amplicon sequencing. Genes Cells, 25, 498–509.

49. Schrinner, S.D., Mari, R.S., Ebler, J., Rautiainen, M., Seillier, L., Reimer, J.J., Usadel, B., Marschall, T. and Klau, G.W. (2020) Haplotype threading: accurate polyploid phasing from long reads. Genome Biol, 21, 252.

50. Xie, M., Wu, Q., Wang, J. and Jiang, T. (2016) H-PoP and H-PoPG: heuristic partitioning algorithms for single individual haplotyping of polyploids. Bioinformatics, 32, 3735–3744.

51. 51 Xiao, A., Wang, Z., Hu, Y., Wu, Y., Luo, Z., Yang, Z., Zu, Y., Li, W., Huang, P., Tong, X. et al.. (2013) Chromosomal deletions and inversions mediated by TALENs and CRISPR/Cas in zebrafish. Nucleic Acids Res, 41, e141.

52. Shin, H.Y., Wang, C., Lee, H.K., Yoo, K.H., Zeng, X., Kuhns, T., Yang, C.M., Mohr, T., Liu, C. and Hennighausen, L. (2017) CRISPR/Cas9 targeting events cause complex deletions and insertions at 17 sites in the mouse genome. Nat Commun, 8, 15464.

53. Adikusuma, F., Piltz, S., Corbett, M.A., Turvey, M., McColl, S.R., Helbig, K.J., Beard, M.R., Hughes, J., Pomerantz, R.T. and Thomas, P.Q. (2018) Large deletions induced by Cas9 cleavage. Nature, 560, E8–E9.

54. Codner, G.F., Mianne, J., Caulder, A., Loeffler, J., Fell, R., King, R., Allan, A.J., Mackenzie, M., Pike, F.J., McCabe, C.V. et al.. (2018) Application of long single-stranded DNA donors in genome editing: generation and validation of mouse mutants. BMC Biol, 16, 70.

55. Mianne, J., Chessum, L., Kumar, S., Aguilar, C., Codner, G., Hutchison, M., Parker, A., Mallon, A.M., Wells, S., Simon, M.M. et al.. (2016) Correction of the auditory phenotype in C57BL/6N mice via CRISPR/Cas9-mediated homology directed repair. Genome Med, 8, 16.

56. Skryabin, B.V., Kummerfeld, D.M., Gubar, L., Seeger, B., Kaiser, H., Stegemann, A., Roth, J., Meuth, S.G., Pavenstadt, H., Sherwood, J. et al.. (2020) Pervasive head-to-tail insertions of DNA templates mask desired CRISPR-Cas9-mediated genome editing events. Sci Adv, 6, eaax2941.

57. Bi, C., Wang, L., Yuan, B., Zhou, X., Li, Y., Wang, S., Pang, Y., Gao, X., Huang, Y. and Li, M. (2020) Long-read individual-molecule sequencing reveals CRISPR-induced genetic heterogeneity in human ESCs. Genome Biol, 21, 213.

58. Gilpatrick, T., Lee, I., Graham, J.E., Raimondeau, E., Bowen, R., Heron, A., Downs, B., Sukumar, S., Sedlazeck, F.J. and Timp, W. (2020) Targeted nanopore sequencing with Cas9- guided adapter ligation. Nat Biotechnol, 38, 433–438.

