## Additional file 2: Fig. S1 for "Multiplex genotyping method to validate the multiallelic genome editing outcomes using machine learning-assisted long-read sequencing"

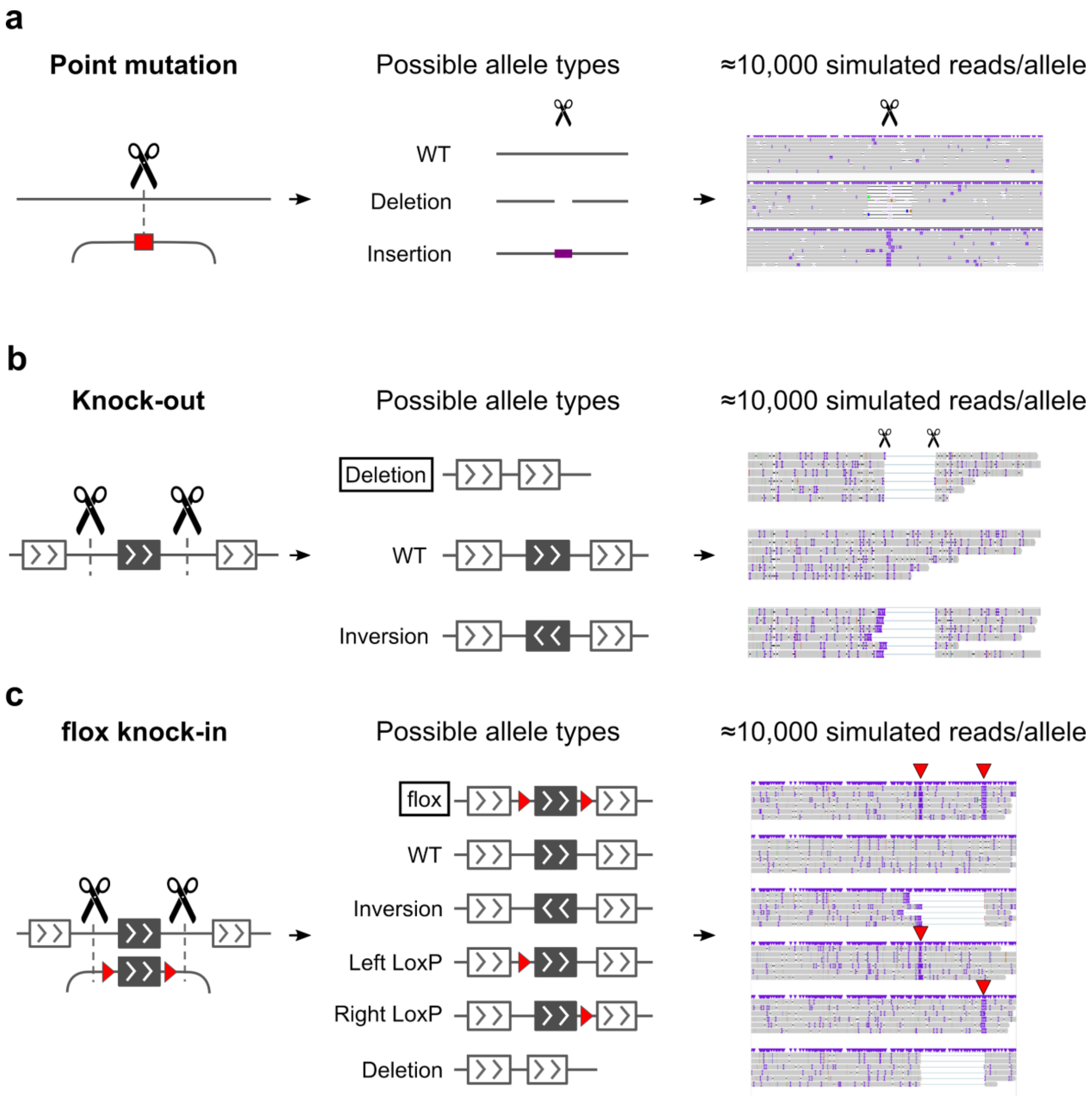


Fig. S1: **Simulated alleles of each genome editing design.**

**a** Point mutation design. Red box represents a target point mutation. Purple bar represents inserted nucleotides. **b** Knock-out design. Black box represents a target exon. Boxed allele type represents the target allele. **c** flox knock-in design. Red triangles represent LoxP sequences.
