## Additional file 2: Fig. S2 for "Multiplex genotyping method to validate the multiallelic genome editing outcomes using machine learning-assisted long-read sequencing"

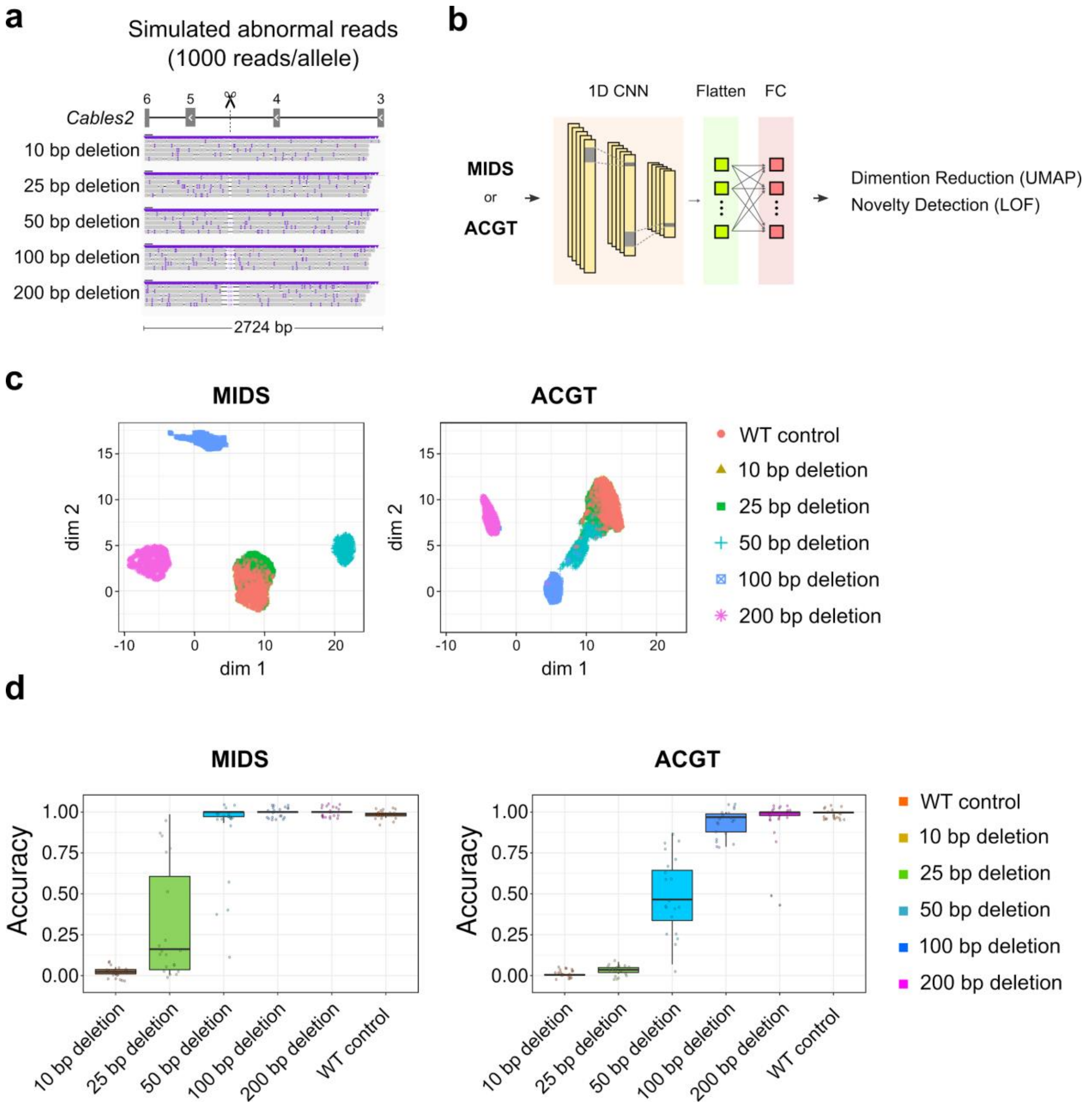


Fig. S2: **Performance evaluation of abnormal allele detection.**

**a** Simulated nanopore sequencing reads of abnormal alleles. The simulated read length was 2724 bp. The integer on the exon represents the exon number. The scissor represents a Cas9-cutting site. **b** Model structure. ‘MIDS’ and ‘ACGT’ mean encoded reads with or without MIDS conversion, respectively. **c** UMAP visualisation of the output vectors from the FC layer. **d** The accuracy of abnormal allele detection with or without MIDS conversion. The 20 dots in each sample on the x-axis represent the iteration of learning and prediction using the deep neural network because the model allowed randomness. In the case of WT control, true positive means a control read is labelled as normal. The accuracy was calculated using the following formula: $accuracy = \frac{TP + TN}{TP + FP + TN + FN}$, where TP, FN, FP, and TN represent the number of true positives, false negatives, false positives, and true negatives, respectively.
