## Additional file 2: Fig. S3 for "Multiplex genotyping method to validate the multiallelic genome editing outcomes using machine learning-assisted long-read sequencing"

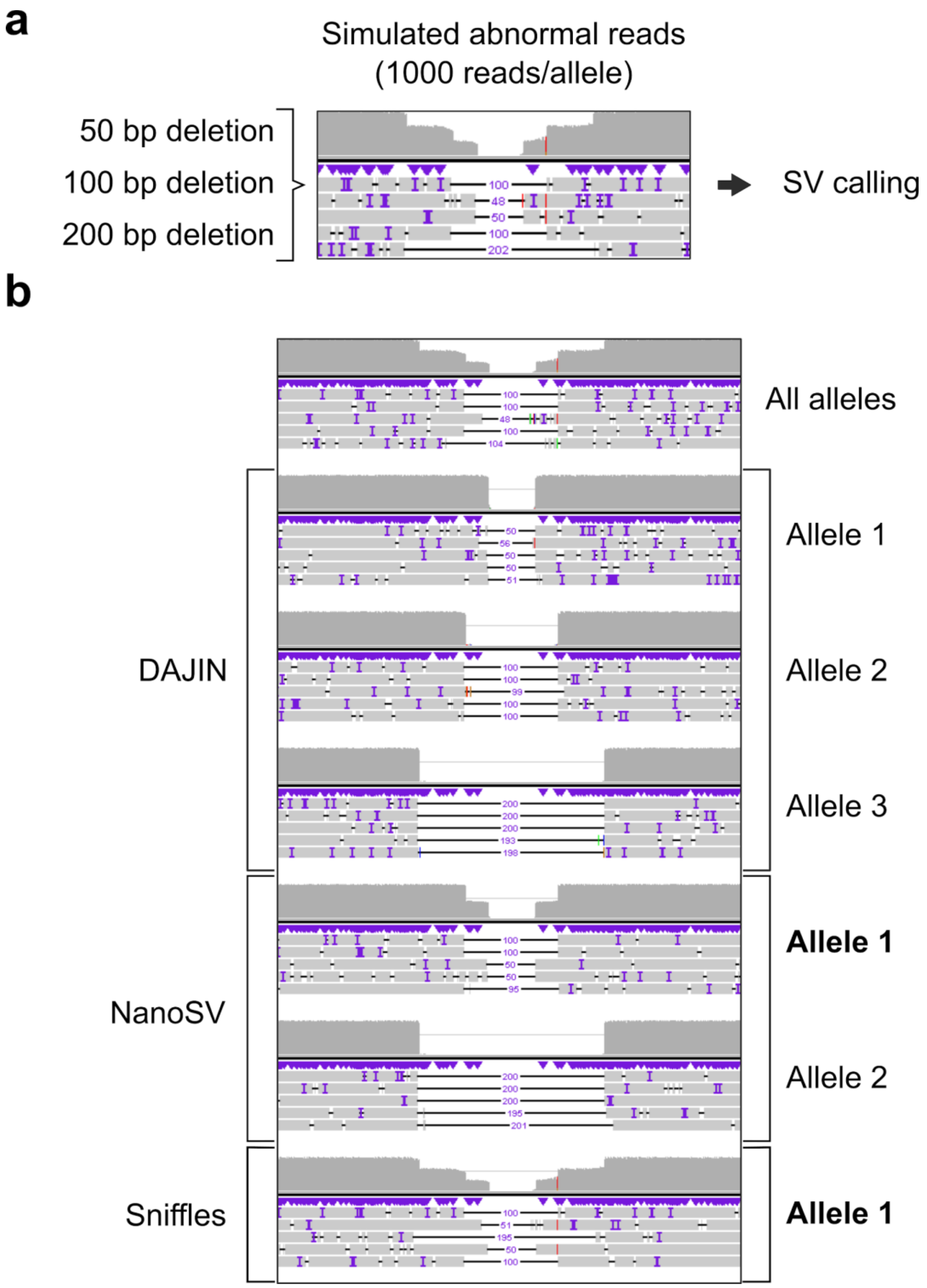


Fig. S3: **Comparison between DAJIN and SV callers**

**a** Three artificial alleles using simulated SV reads. **b** Comparison of DAJIN, NanoSV, and Sniffles. The alleles in bold font represent unclassified alleles.
