## Additional file 2: Fig. S4 for "Multiplex genotyping method to validate the multiallelic genome editing outcomes using machine learning-assisted long-read sequencing"

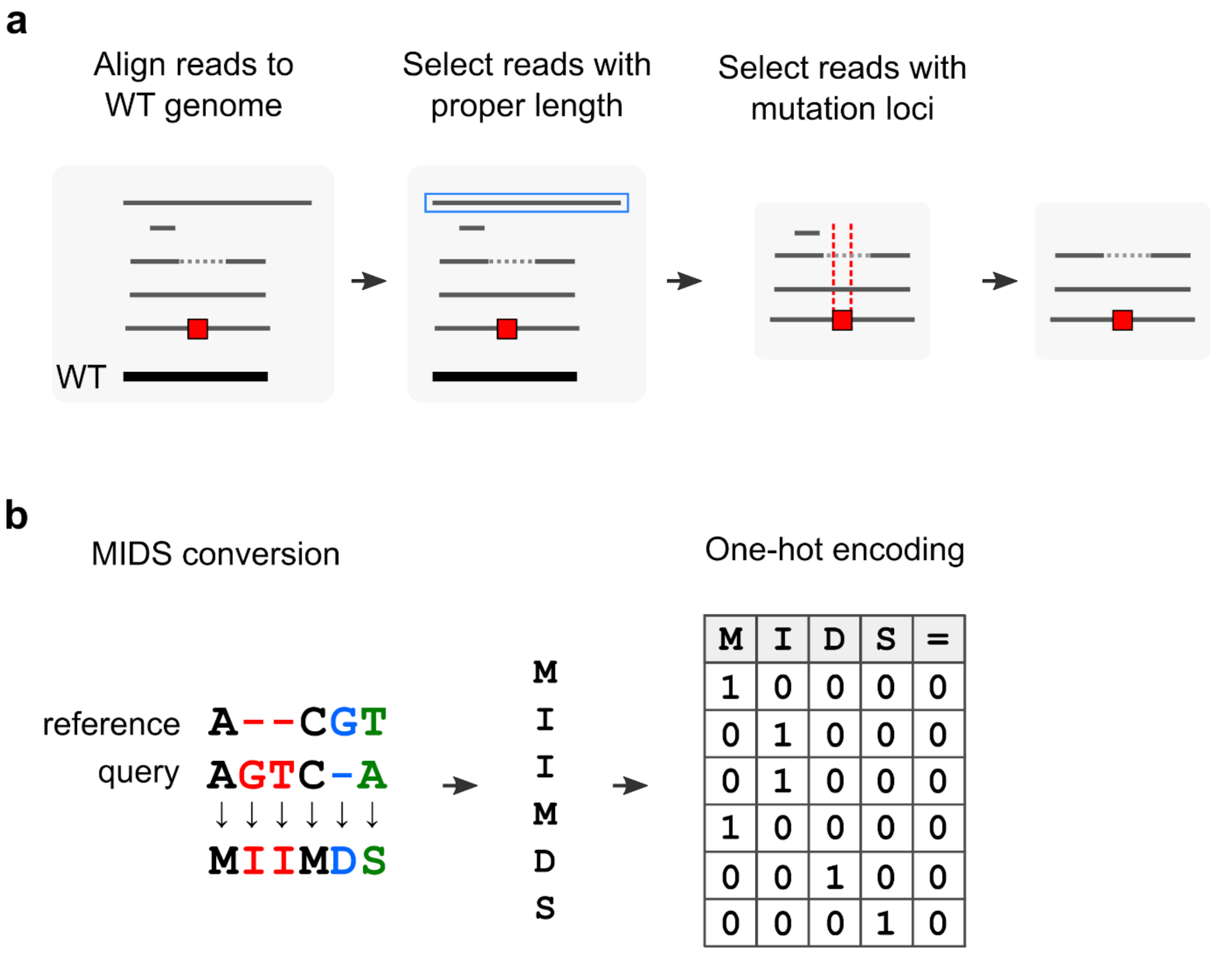


Fig. S4: **Pre-processing**

**a** Screening of reads with proper sequence length and mutation loci. Grey bars represent reads. Dotted bars represent deleted nucleotides. The red box represents the target mutation. The blue boxed bar represents a read exceeding the allowable length. Red dotted vertical lines represent target mutation loci. **b** MIDS conversion and One-hot encoding. The ‘reference’ and ‘query’ mean WT sequence and nanopore reads, respectively.
