## Additional file 2: Fig. S5 for "Multiplex genotyping method to validate the multiallelic genome editing outcomes using machine learning-assisted long-read sequencing"

**a**

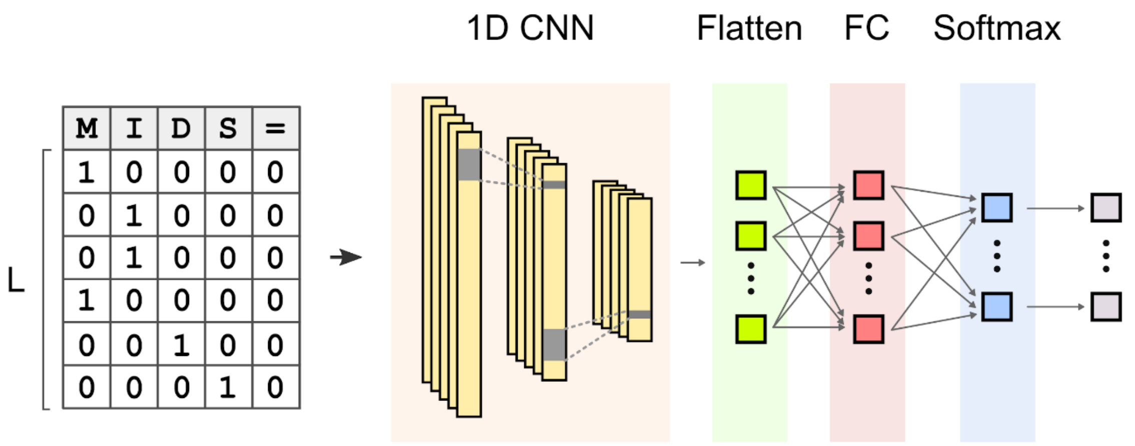


**b**

| Layer name | CNN-1 | CNN-2 | CNN-3 | Flatten | FC | Softmax |
|---|---|---|---|---|---|---|
| Output shape | (L, 32) | (L/12, 32) | (L/64, 32) | L/64 x 32 | 32 | Number of labels |
| Activation | ReLU | ReLU | ReLU | - | ReLU | Softmax |
| Filters | 32 | 32 | 32 | - | - | - |
| Kernel size | 256 | 128 | 64 | - | - | - |
| Max pooling size | 12 | 6 | 3 | - | - | - |

Fig. S5: **The architecture of deep neural network models**

**a** Model structure. The input of the model is the encoded nanopore sequence with length (L). Three layers of a one-dimensional convolutional neural network (1D-CNN) include max-pooling layers and activation functions. The outputs of 1D-CNN layers are joined together into one vector by flattening. Each neuron in the flattened layer is attached to the fully connected (FC) layer. The neurons in the output layer use the softmax function as the activation function, whereas all the neurons in other layers use ReLU as the activation function. **b** Parameter setting for each layer.
