## Additional file 2: Fig. S6 for "Multiplex genotyping method to validate the multiallelic genome editing outcomes using machine learning-assisted long-read sequencing"

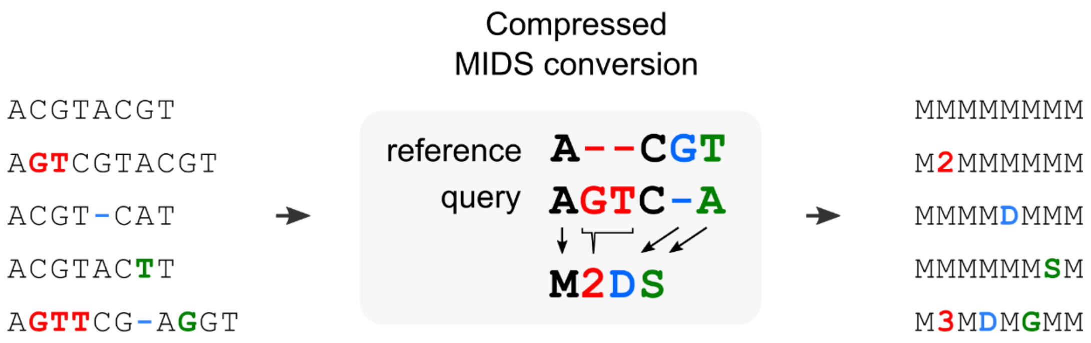


Fig. S6: **Compressed MIDS conversion**

Red and green colours represent insertion and substitution, respectively. Blue colour represents deletion.
