## Additional file 2: Fig. S7 for "Multiplex genotyping method to validate the multiallelic genome editing outcomes using machine learning-assisted long-read sequencing"

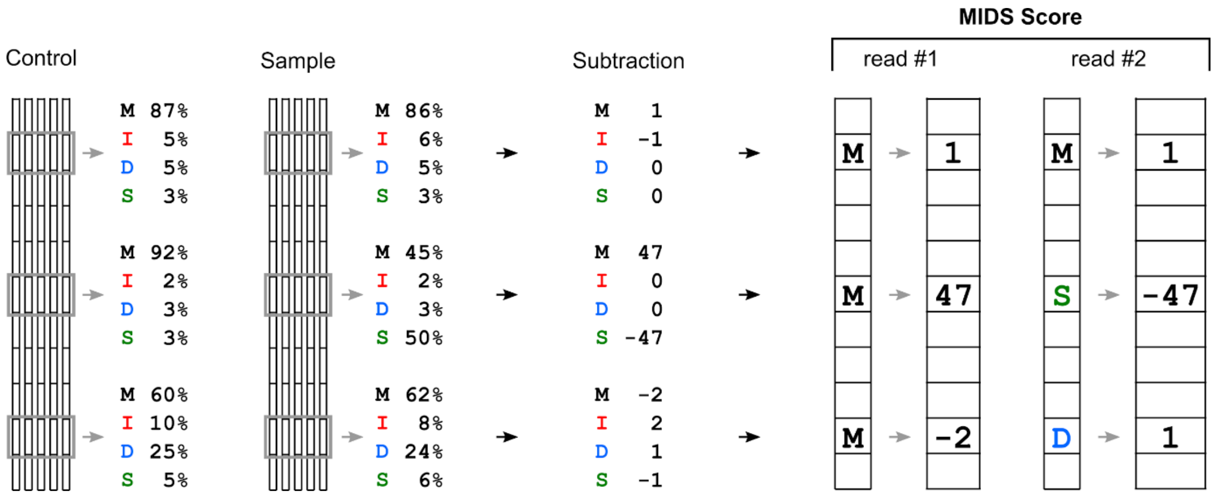


Fig. S7: **MIDS score**

The black boxes represent sequences after compressed MIDS conversion. The grey boxes show representative base positions. MIDS score is calculated by subtracting the relative frequency of MIDS between a control and a sample.
