## Supplementary figures and images for "Multiplex genotyping method to validate the multiallelic genome editing outcomes using machine learning-assisted long-read sequencing"

### Additional file 2: Fig. S8

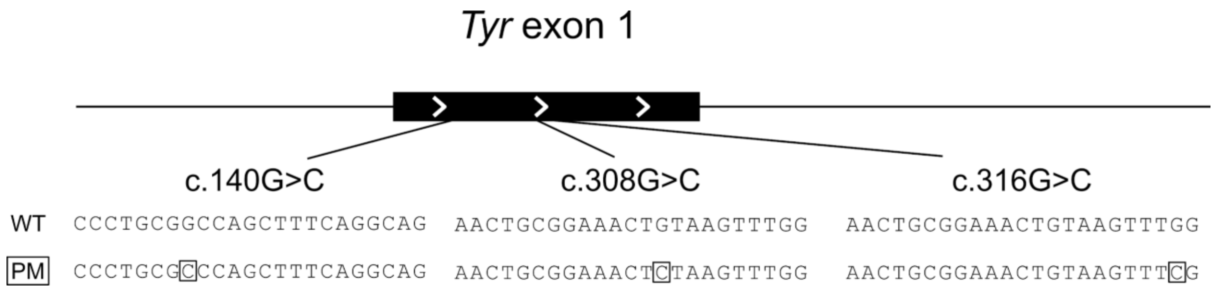


Fig. S8: ***Tyr* c.140G>C, c.316G>C, and c.308G>C point mutation (PM) design**

The boxed nucleotides represent intended PMs.

### Additional file 2: Fig. S23

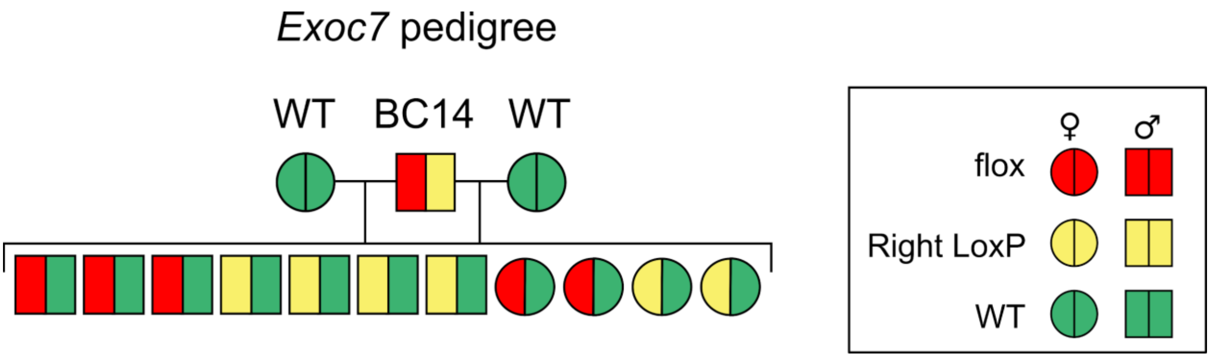


Fig. S23: **Pedigree line of BC14 in *Exoc7* flox knock-in design**
