## Additional file 2: Fig. S9 for "Multiplex genotyping method to validate the multiallelic genome editing outcomes using machine learning-assisted long-read sequencing"

BC21: Allele 2 Intended PM (93.6 %)

```
TGCATTGAAGCAGTTCACCAAAATAACAAAGTAACAAAGTAAGATATCTTTGGAATAATCAATTCAAGATAATCAAGGAAAAATGAGAGGCAACTA
TTTTAGACTGATTACTTTTATAAAATAAATAAGCTCAGCTTAGCCAGATATAAGCAATATTCTGAGTTCTGAAGAAAAATTTTTGACAAAATGAGT
TCTATAAATGTTATTGTCTACTTATGATCTCTAAATACAACAGGCTTGTATTCAGAATCTAGATGTTTCATGACCTTTATTCATAAGAGATGATGT
ATTCTTGATACTACTTCTCATTTGCAAATTCCAATTATTATTAATTTCATATCAATTAGAATAATATATCTTCCTTCAATTTAGTTACCTCACTAT
GGGCTATGTACAAACTCCAAGAAAAAGTTAGTCATGTGCTTTGCAGAAGATAAAAGCTTAGTGTAAAACAGGCTGAGAGTATTTGATGTAAGAAGG
GGAGTGGTTATATAGGTCTTAGCCAAAACATGTGATAGTCACTCCAGGGGTTGCTGGAAAAGAAGTCTGTGACACTCATTAACCTATTGGTGCAGA
TTTTGTATGATCTAAAGGAGAAAATGTTCTTGGCTGTTTTGTATTGCCTTCTGTGGAGTTTCCAGATCTCTGATGGCCATTTTCCTCGAGCCTGTG
CCTCCTCTAAGAACTTGTTGGCAAAAGAATGCTGCCCACCATGGATGGGTGATGGGAGTCCCTGCGGCCAGCTTTCAGGCAGAGGTTCCTGCCAGG
ATATCCTTCTGTCCAGTGCACCATCTGGACCTCAGTTCCCCTTCAAAGGGGTGGATGACCGTGAGTCCTGGCCCTCTGTGTTTTATAATAGGACCT
GCCAGTGCTCAGGCAACTTCATGGGTTTCAACTGCGGAAACTCTAAGTTTGGATTTGGGGGCCCAAATTGTACAGAGAAGCGAGTCTTGATTAGAA
GAAACATTTTTGATTTGAGTGTCTCCGAAAAGAATAAGTTCTTTTCTTACCTCACTTTAGCAAAACATACTATCAGCTCAGTCTATGTCATCCCCA
CAGGCACCTATGGCCAAATGAACAATGGGTCAACACCCATGTTTAATGATATCAACATCTACGACCTCTTTGTATGGATGCATTACTATGTGTCAA
GGGACACACTGCTTGGGGGCTCTGAAATATGGAGGGACATTGATTTTGCCCATGAAGCACCAGGGTTTCTGCCTTGGCACAGACTTTTCTTGTTAT
TGTGGGAACAAGAAATTCGAGAACTAACTGGGGATGAGAACTTCACTGTTCCATACTGGGATTGGAGAGATGCAGAAAACTGTGACATTTGCACAG
ATGAGTACTTGGGAGGTCGTCACCCTGAAAATCCTAACTTACTCAGCCCAGCATCCTTCTTCTCCTCCTGGCAGGTAAGATGCACTATATAGAGAG
AGTTGCAAAGACTGGTACTTCAGCAGCCACATTTTCATGCTCTGTGAGCATCTCTGATAATATCTCAGGGCAGAAAATGTGCCTTACTAACAGATG
TTAATGCTTCTTGATTTCTTTTTCTCTTTTGAGAACTCTTCAAAGTTGTTATTAAACAAATATCTATGTGCTTATTTGTCTTAATATCTAACAGCT
TAGTTAGATTTCTAAGCTGCTATAAACAAGGACTGATTGGTTCACCACTGTATTGTTAGCACCTCCTATGGTATCTGGAATAAACAGTAACTCAGT
GCGGAAACTCTAAAATATAAATCTCTGGCCAAAACCAAGACTTATTATTCAGGATCTTCAAGAGAAAGTGCTGAGATAATTCACTAAGTATCAGAG
ATGACCTTTATTACATGATTGCCTGATAGAAAAAATGATTACACACACACAAAAAAAATCTTCAGTTGCTTAAATTTTAAACGTTGCTGACTCTCAA
ACAGTTAAGTAATAAAAGAGTTAAAGCCTGCTGTGTATTTAGAATATGTGAATACCTATTGAAAGAATTTATTGTACAATTAATATAAACAGACTT
CTATTTTACAGTCATAAGATACTACTTAATTTGTTAAAAATTATTTTTTGATAGCATTGTTGGTAAATAGCAAAGGTGATATTTGCTAATGATTAC
AAGGGCTGTCTGGCTAACTTACGTTATGTTCAGGGAGAAGACAGTCCTTTTTAAGGAATGGGCACTTTCTAACTTTTTTTCTCTAGGATGGAGAAA
AATTAGCCTTCTTCCTACTTTAAAAATGTTAGACATAGAATTAAGGGATTGTTATTTTGAGATTAAATTTTCTTTTCTCCTATTATTTTTCCTCAT
TCTGGAATGGAAGCAAAAGATGAAGAAAGAAATATATGTTAAATTGTTTTCCTTTAAATGAACACAAATGTGAAATATGTTTTTCTGCCTATCTTG
TAAAATTTTCTATTGCAACTATTCTGATTACCAGTTCAAATGGGGAAAAAAGAACATAGGCTACCCCACACTTGAAATTTTGAAATATGAATGTCC
TCTGTCTCTGCTGGTCTAACACTTCCAAAATGGAAACCTTTAAAGGGCCACTGTAAATTACAGCTGCTAATTCCTGGTGCCAATGGTGATAAGTGT
TTACTAAACCTAGTGAGTACTTTATAGCATGGGGCTCTGCTGCGAAGTAACATTGCTGTATATTTTCAGTCATTCTACCTTAATTCATGAACTGCA
AAACTCTCATCTAGCTTTTTACTTCTCTAGCTATTGCTTTAAGTTCTATCAGGCTCAGGTGTGGAATTCTC
```

Insertion Deletion Substitution

G G T T T C A A C T G C G G A A A C T C T A A G T T T G G A T T T G G

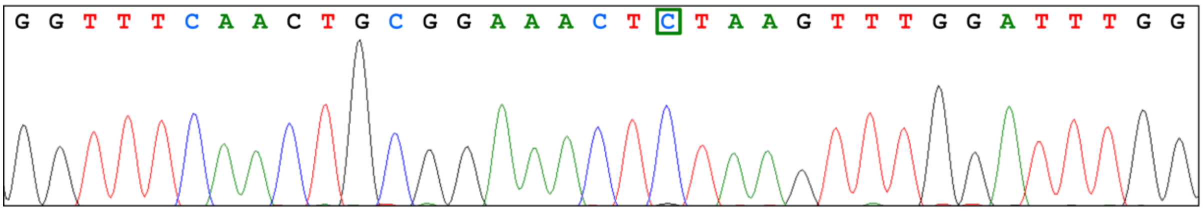


Fig. S9: **DAJIN's consensus sequence and Sanger sequencing of *Tyr* c.308G>C BC21**

The sequence represents the consensus sequence of *Tyr* c.308G>C BC21. The green highlighted nucleotides represent substitution. The boxed sequences in the consensus sequences are captured via Sanger sequencing.
