## Additional file 2: Fig. S10 for "Multiplex genotyping method to validate the multiallelic genome editing outcomes using machine learning-assisted long-read sequencing"

BC31: Allele 2 Mutated WT (93.8%)

TGCATTGAAGCAGTTCACCAAAATAACAAAGTAACAAAGTAAGATATCTTTGGAATAATCAATTCAAGATAATCAAGGAAAAATGAGAGGCAACTA
TTTTAGACTGATTACTTTTATAAAATAAATAAGCTCAGCTTAGCCAGATATAAGCAATATTCTGAGTTCTGAAGAAAAATTTTTGACAAAATGAGT
TCTATAAATGTTATTGTCTACTTATGATCTCTAAATACAACAGGCTTGTATTCAGAATCTAGATGTTTCATGACCTTTATTCATAAGAGATGATGT
ATTCTTGATACTACTTCTCATTTGCAAATTCCAATTATTATTAATTTCATATCAATTAGAATAATATATCTTCCTTCAATTTAGTTACCTCACTAT
GGGCTATGTACAAACTCCAAGAAAAAGTTAGTCATGTGCTTTGCAGAAGATAAAAGCTTAGTGTAAAACAGGCTGAGAGTATTTGATGTAAGAAGG
GGAGTGGTTATATAGGTCTTAGCCAAAACATGTGATAGTCACTCCAGGGGTTGCTGGAAAAGAAGTCTGTGACACTCATTAACCTATTGGTGCAGA
TTTTGTATGATCTAAAGGAGAAAATGTTCTTGGCTGTTTTGTATTGCCTTCTGTGGAGTTTCCAGATCTCTGATGGCCATTTTCCTCGAGCCTGTG
CCTCCTCTAAGAACTTGTTGGCAAAAGAATGCTGCCCACCATGGATGGGTGATGGGAGTCCCTGCGGCCAGCTTTCAGGCAGAGGTTCCTGCCAGG
ATATCCTTCTGTCCAGTGCACCATCTGGACCTCAGTTCCCCTTCAAAGGGGTGGATGACCTTGAGTCCTGGCCCTCTGTGTTTTATAATAGGACCT
GCCAGTGCTCAGGCAACTTCATGGGTTTCAACTGCGGAAACTGTAAGTTTGGATTTGGGGGCCCAAATTGTACAGAGAAGCGAGTCTTGATTAGAA
GAAACATTTTTGATTTGAGTGTCTCCGAAAAGAATAAGTTCTTTTCTTACCTCACTTTAGCAAAACATACTATCAGCTCAGTCTATGTCATCCCCA
CAGGCACCTATGGCCAAATGAACAATGGGTCAACACCCATGTTTAATGATATCAACATCTACGACCTCTTTGTATGGATGCATTACTATGTGTCAA
GGGACACACTGCTTGGGGGCTCTGAAATATGGAGGGACATTGATTTTGCCCATGAAGCACCAGGGTTTCTGCCTTGGCACAGACTTTTCTTGTTAT
TGTGGGAACAAGAAATTCGAGAACTAACTGGGGATGAGAACTTCACTGTTCCATACTGGGATTGGAGAGATGCAGAAAACTGTGACATTTGCACAG
ATGAGTACTTGGGAGGTCGTCACCCTGAAAATCCTAACTTACTCAGCCCAGCATCCTTCTTCTCCTCCTGGCAGGTAAGATGCACTATATAGAGAG
AGTTGCAAAGACTGGTACTTCAGCAGCCACATTTTCATGCTCTGTGAGCATCTCTGATAATATCTCAGGGCAGAAAATGTGCCTTACTAACAGATG
TTAATGCTTCTTGATTTCTTTTTCTCTTTTGAGAACTCTTCAAAGTTGTTATTAAACAAATATCTATGTGCTTATTTGTCTTAATATCTAACAGCT
TAGTTAGATTTCTAAGCTGCTATAAACAAGGACTGATTGGTTCACCACTGTATTGTTAGCACCTCCTATGGTATCTGGAATAAACAGTAACTCAGT
TATTTAAGAATGGATGAGAAACCAGATTATCTTAGTTCATGTTTCTGAGTAATATTAAAATTAATATTAACAGTAAAATCCATAAGTATGCTACTT
TAAAATATAAATCTCTGGCCAAAACCAAGACTTATTATTCAGGATCTTCAAGAGAAAGTGCTGAGATAATTCACTAAGTATCAGAGATGACCTTTA
TTACATGATTGCCTGATAGAAAAAATGATTACACACACACAAAAAAATCTTCAGTTGCTTAAATTTTAAACGTTGCTGACTCTCAAACAGTTAAGT
AATAAAAGAGTTAAAGCCTGCTGTGTATTTAGAATATGTGAATACCTATTGAAAGAATTTATTGTACAATTAATATAAACAGACTTCTATTTTACA
GTCATAAGATACTACTTAATTTGTTAAAAATTATTTTTTGATAGCATTGTTGGTAAATAGCAAAGGTGATATTTGCTAATGATTACAAGGGCTGTC
TGGCTAACTTACGTTATGTTCAGGGAGAAGACAGTCCTTTTTAAGGAATGGGCACTTTCTAACTTTTTTTCTCTAGGATGGAGAAAAATTAGCCTT
CTTCCTACTTTAAAAATGTTAGACATAGAATTAAGGGATTGTTATTTTGAGATTAAATTTTCTTTTCTCCTATTATTTTTCCTCATTCTGGAATGG
AAGCAAAAGATGAAGAAAGAAATATATGTTAAATTGTTTTCCTTTAAATGAACACAAATGTGAAATATGTTTTTCTGCCTATCTTGTAAAATTTTC
TATTGCAACTATTCTGATTACCAGTTCAAATGGGGAAAAAAGAACATAGGCTACCCCACACTTGAAATTTTGAAATATGAATGTCCTCTGTCTCTG
CTGGTCTAACACTTCCAAAATGGAAACCTTTAAAGGGCCACTGTAAATTACAGCTGCTAATTCCTGGTGCCAATGGTGATAAGTGTTTACTAAACC
TAGTGAGTACTTTATAGCATGGGGCTCTGCTGCGAAGTAACATTGCTGTATATTTTCAGTCATTCTACCTTAATTCATGAACTGCAAAACTCTCAT
CTAGCTTTTTACTTCTCTAGCTATTGCTTTAAGTTCTATCAGGCTCAGGTGTGGAATTCTC

Insertion Deletion Substitution

Fig. S10: **DAJIN's consensus sequence of *Tyr* c.230G>T point mutation**

The green highlighted nucleotide represents a substitution mutation.
