## Additional file 2: Fig. S11 for "Multiplex genotyping method to validate the multiallelic genome editing outcomes using machine learning-assisted long-read sequencing"

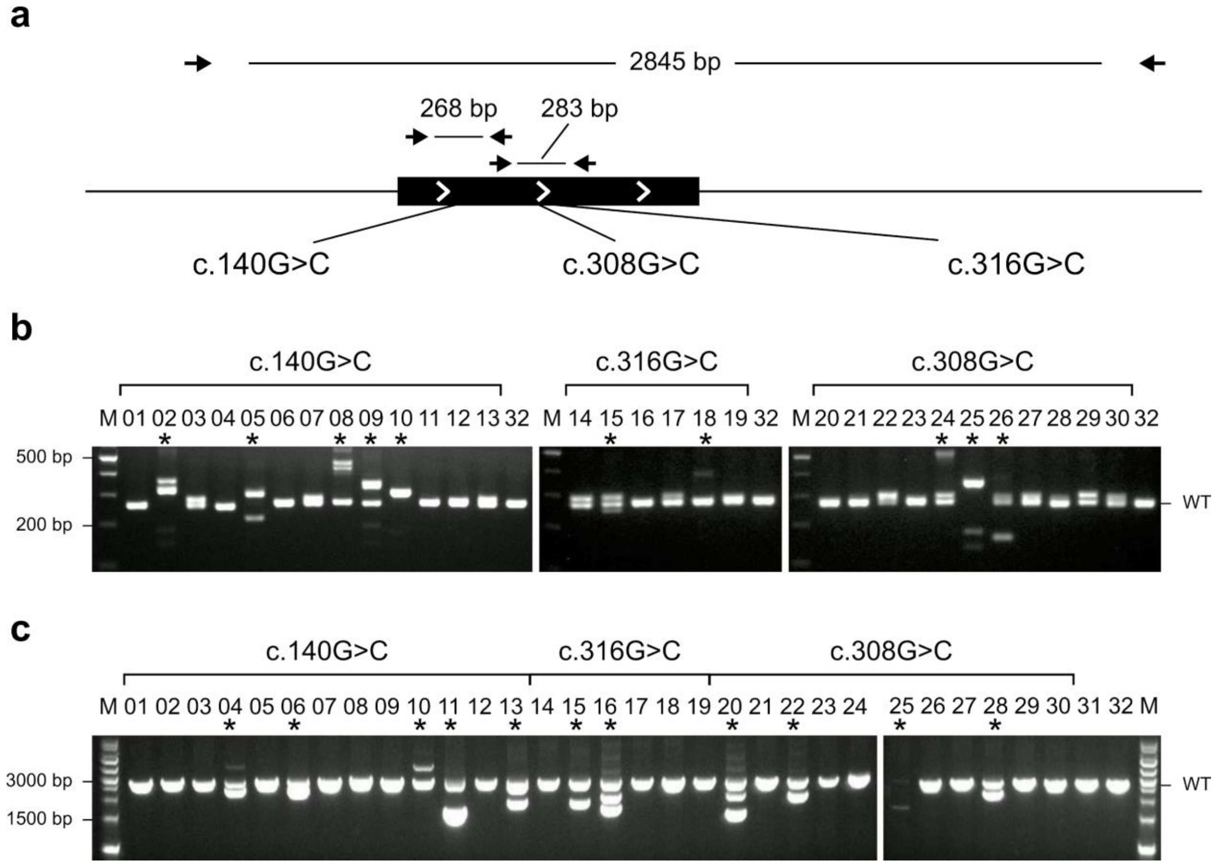


Fig. S11: **PCR-based genotyping of *Tyr* PM design**

**a** Genome editing design. The arrows represent PCR primers for short and long PCR. **b** The short PCR results for the detection of small indel alleles. The numbers on the panel represent barcode IDs. The asterisks represent the samples with small indels. **c** The long PCR results for the SV detection. The numbers on the panel represent barcode IDs. The asterisks represent the samples with SVs.
