## Additional file 2: Fig. S12 for "Multiplex genotyping method to validate the multiallelic genome editing outcomes using machine learning-assisted long-read sequencing"

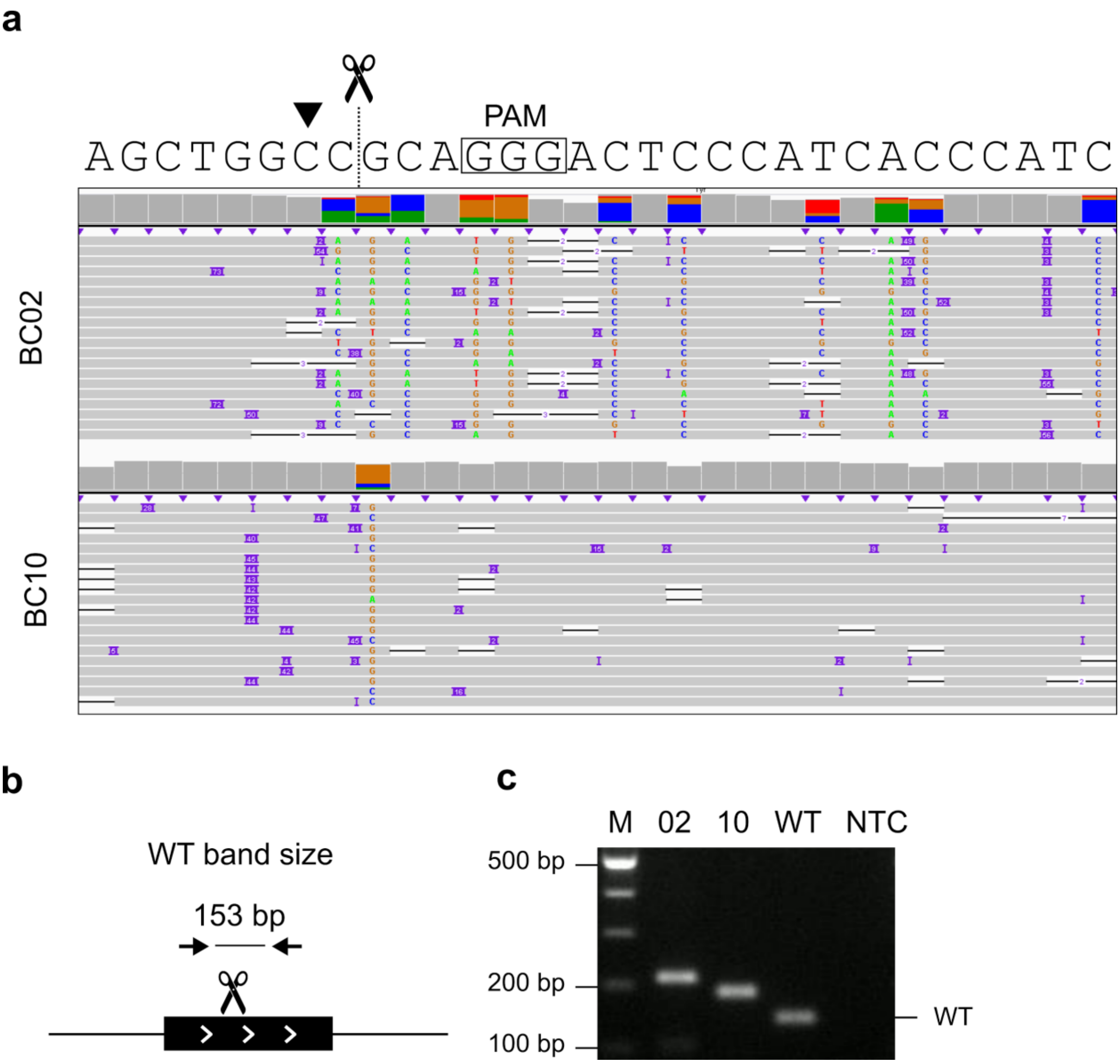


Fig. S12: **Verification of insertion mutation of *Tyr* c.140G>C BC02 and BC10 mice**

**a** Visualisation of nanopore sequencing reads of BC02 and BC10. The scissors and dotted line represent a Cas-cutting site. Arrowhead represents the target nucleotide. **b** PCR design. **c** PCR results for the detection of insertion alleles. NTC means no template control.
