## Additional file 2: Fig. S13 for "Multiplex genotyping method to validate the multiallelic genome editing outcomes using machine learning-assisted long-read sequencing"

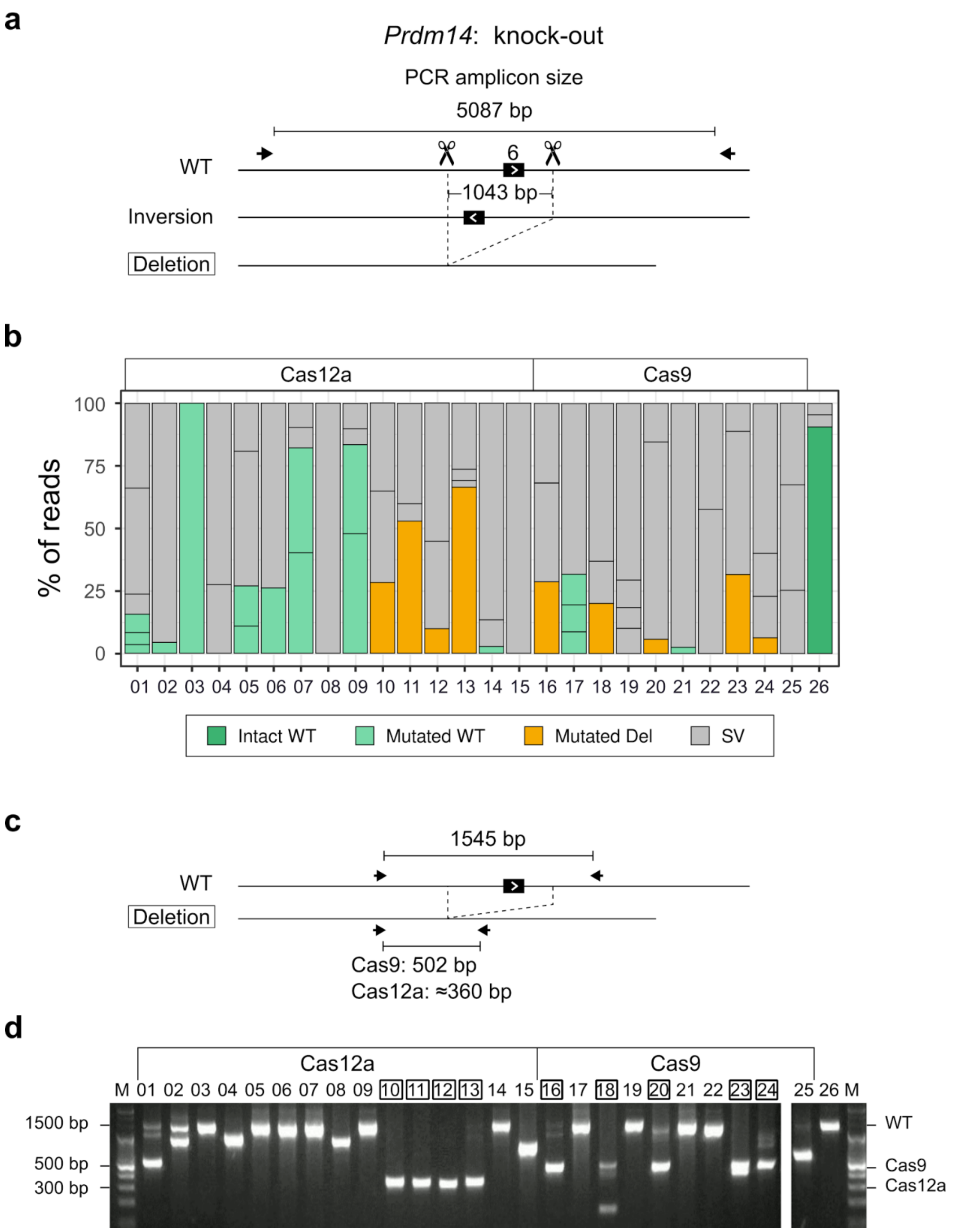


Fig. S13: **DAJIN application to *Prdm14* knock-out design using Cas9 and Cas12a**

**a** Genome editing design for *Prdm14* knock-out using Cas9 and Cas12a. Black boxes represent exon-coding sequences numbered 6. The scissors and dotted lines represent Cas-cutting sites. The arrows represent PCR primers. The boxed allele type represents the target allele. The inversion allele represents a possible by-product. **b** DAJIN's report of the allele percentage. The barcode numbers on the x-axis represent mouse IDs. The BC01–BC15 and BC16–BC25 are treated by Cas12a and Cas9, respectively. The barplot of Cas9 samples is the same plot as shown in Fig. 4b. BC26 is a WT control. The y-axis represents the percentage of DAJIN-reported alleles. The colours of the bar represent DAJIN-reported allele types. The horizontal lines in a bar represent the DAJIN-reported alleles. **c** Design of a short PCR to validate the target deletion allele. The arrows represent PCR primers for the digested DNA fragments, including the size of the PCR products. **d** PCR results for the detection of the target deletion allele. The number on the panel means barcode IDs. The boxed number represents the samples with deletion alleles. 'Cas9' and 'Cas12a' represent expected positions of deletion bands by Cas9 and Cas12a cutting.
