## Additional file 2: Fig. S15 for "Multiplex genotyping method to validate the multiallelic genome editing outcomes using machine learning-assisted long-read sequencing"

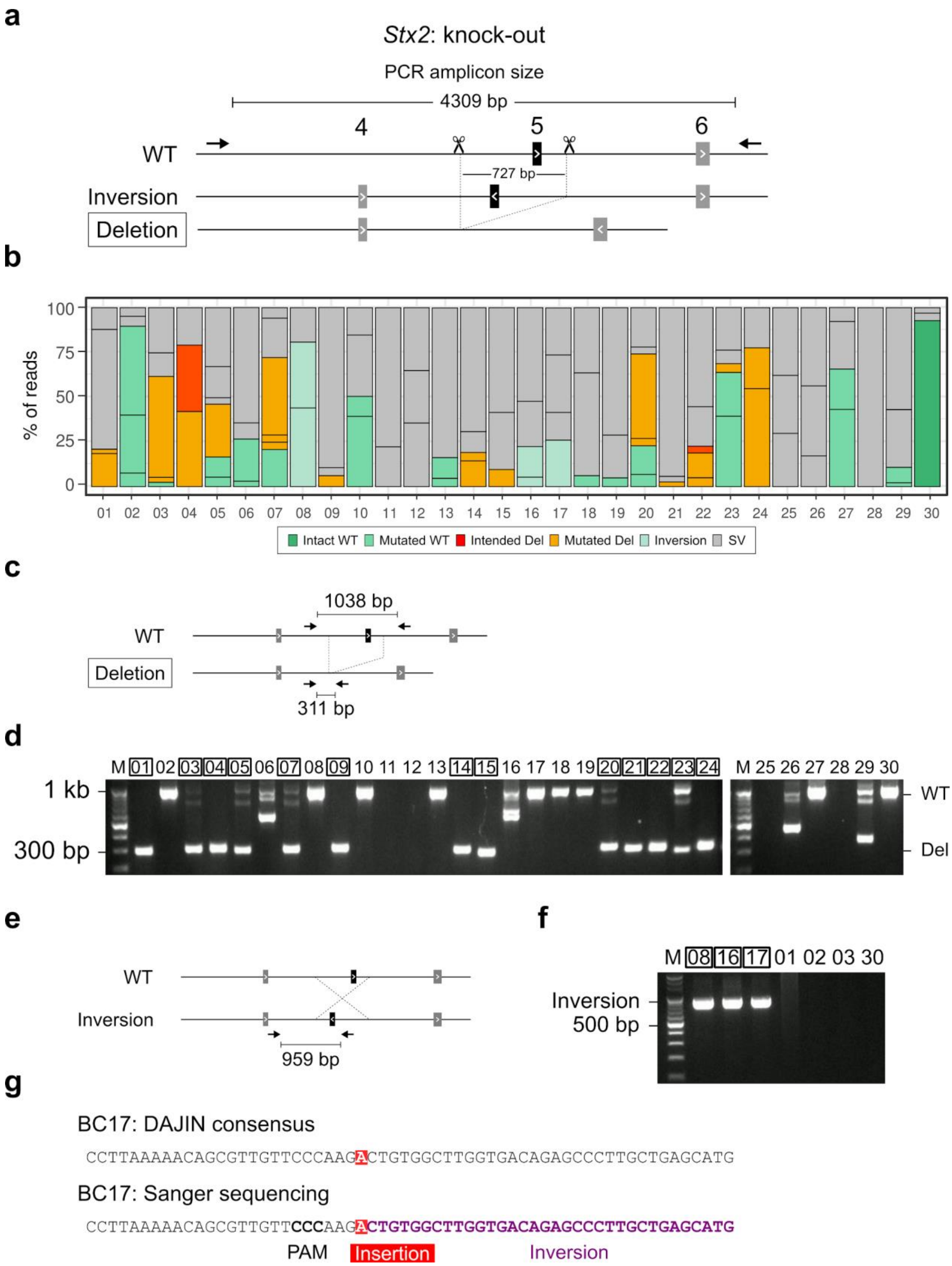


Fig. S15: **DAJIN application to *Stx2* knock-out design**

**a** Genome editing design for *Stx2* knock-out. Shaded and black boxes represent exon-coding sequences numbered 4–6. The scissors and dotted lines represent Cas9-cutting sites. The arrows represent PCR primers. The boxed allele type represents the target allele. The inversion allele represents a possible by-product. **b** DAJIN's report of the allele percentage. The barcode numbers on the x-axis represent mouse IDs. BC30 is a WT control. The y-axis represents the percentage of DAJIN-reported alleles. The colours of the bar represent DAJIN-reported allele types. The horizontal lines in a bar represent the DAJIN-reported alleles. **c** PCR design to validate target deletion allele. The arrows represent PCR primers for the digested DNA fragments, including the size of PCR products. **d** PCR results for the detection of the target deletion allele. The number on the panel represents barcode IDs. The boxed number represents the samples with deletion alleles. **e** PCR design to validate inversion allele. The arrows represent PCR primers for the digested DNA fragments, including the size of PCR products. **f** PCR results for the detection of inversion allele. The number on the panel means barcode IDs. The boxed number represents the samples with deletion alleles. **g** Comparison between DAJIN's consensus sequence and Sanger sequencing of the inversion allele of BC17. The red and purple highlighted nucleotides represent insertion and inversion, respectively.
