## Additional file 2: Fig. S16 for "Multiplex genotyping method to validate the multiallelic genome editing outcomes using machine learning-assisted long-read sequencing"

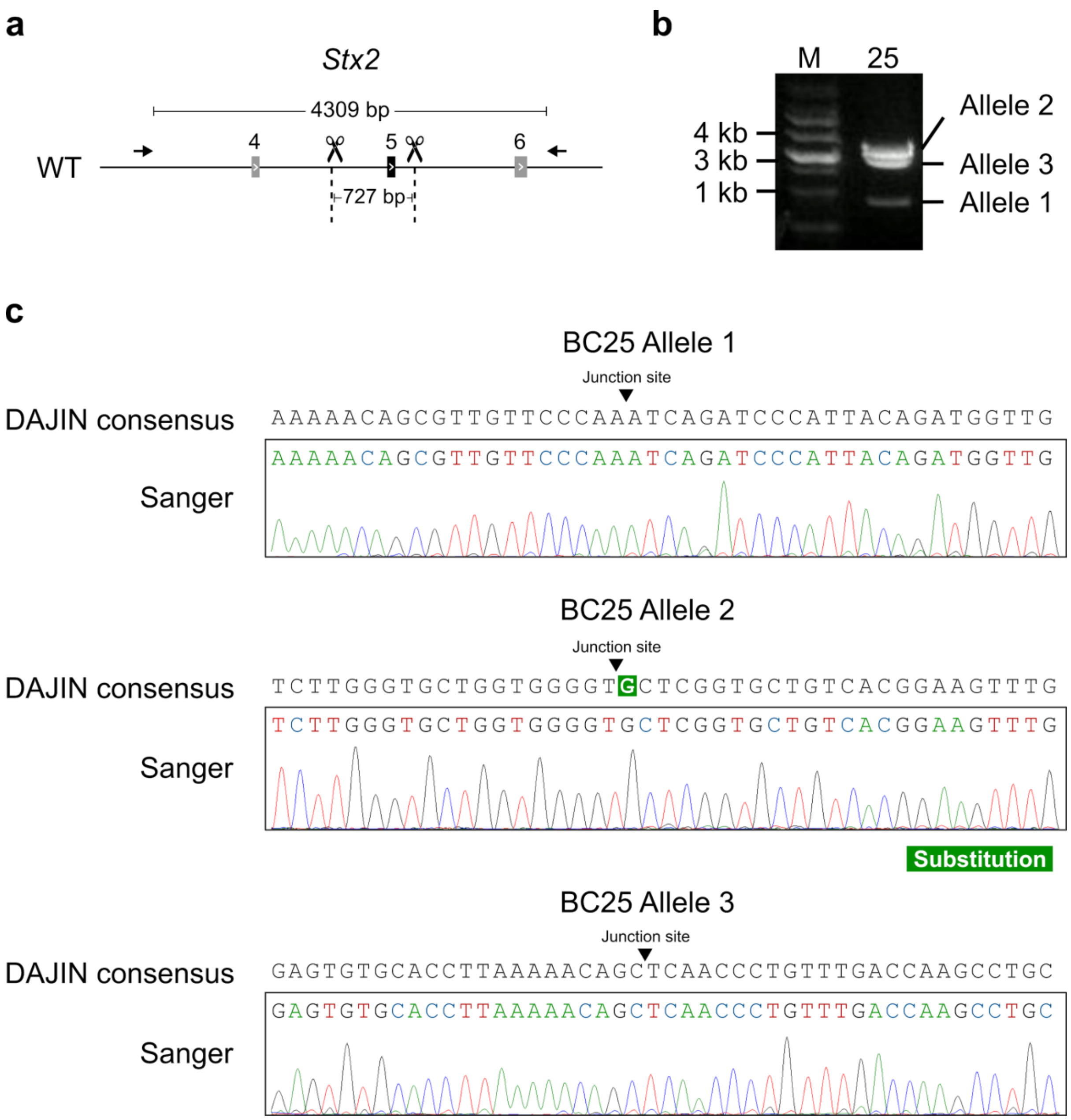


Fig. S16: **Validation of DAJIN-reported SV alleles in *Stx2* BC25**

**a** PCR design to validate SV alleles. The arrows represent PCR primers. **b** PCR results for the detection of SV alleles. The numbers on the panel are the barcode IDs. **c**. Comparison between DAJIN's consensus sequence and Sanger sequencing. The green highlighted nucleotide represents a substitution.
