## Additional file 2: Fig. S17 for "Multiplex genotyping method to validate the multiallelic genome editing outcomes using machine learning-assisted long-read sequencing"

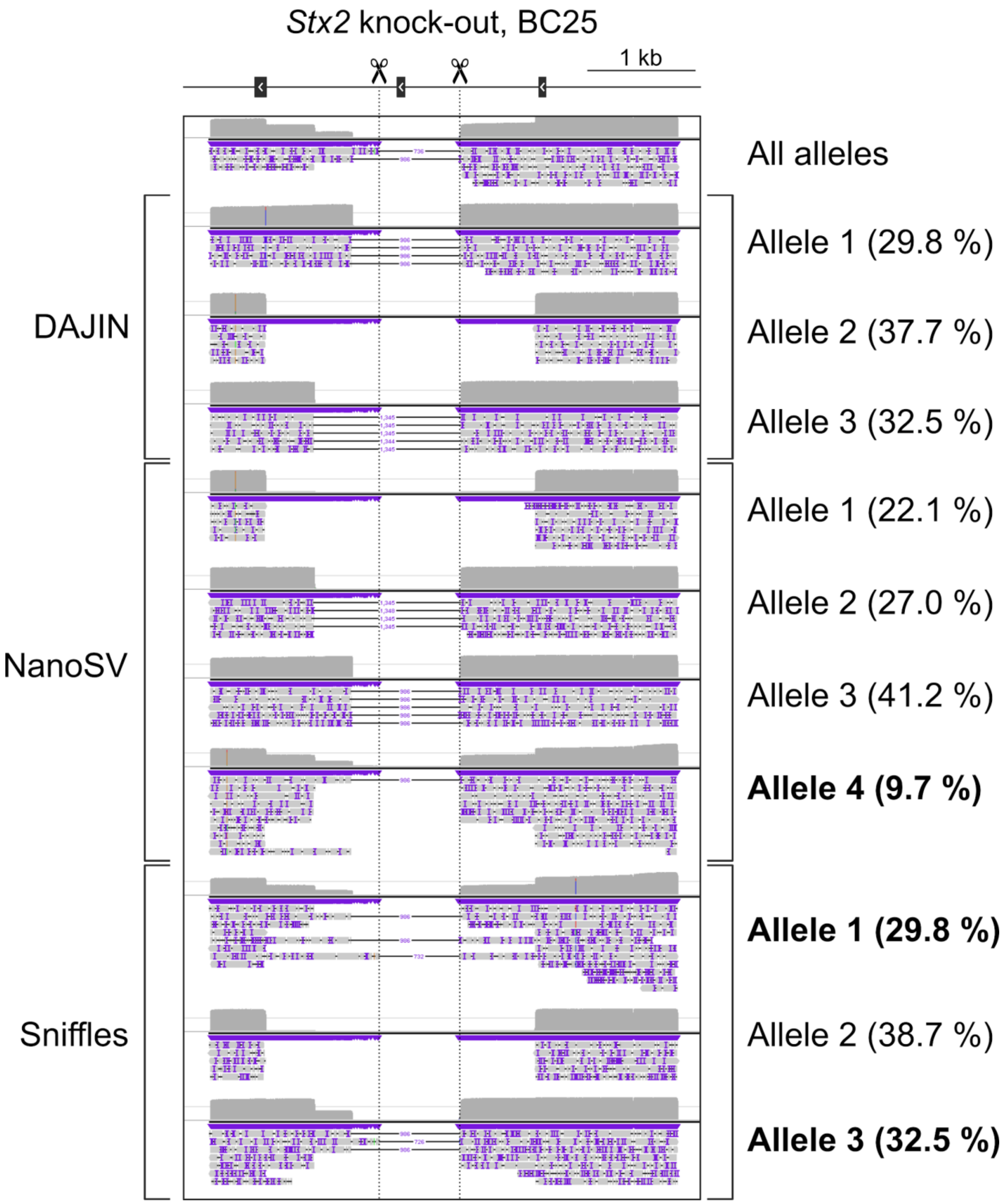


Fig. S17: **Comparison between DAJIN and SV callers**

The alleles in bold font represent misclassified alleles.
