## Additional file 2: Fig. S18 for "Multiplex genotyping method to validate the multiallelic genome editing outcomes using machine learning-assisted long-read sequencing"

### Mixture of intended flox and WT #1

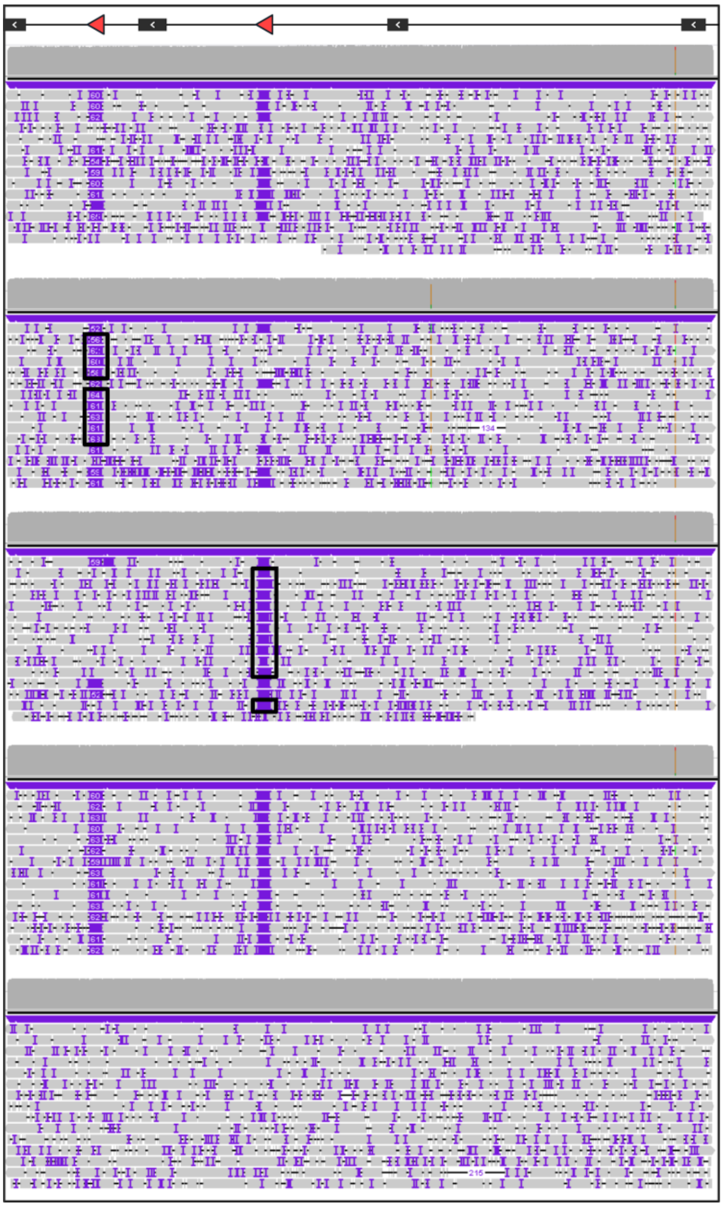


Fig. S18: **Pseudo-LoxP alleles**

Visualisation of simulated reads at *Cables2* locus. The black boxes represent pseudo-LoxPs.
