## Additional file 2: Fig. S19 for "Multiplex genotyping method to validate the multiallelic genome editing outcomes using machine learning-assisted long-read sequencing"

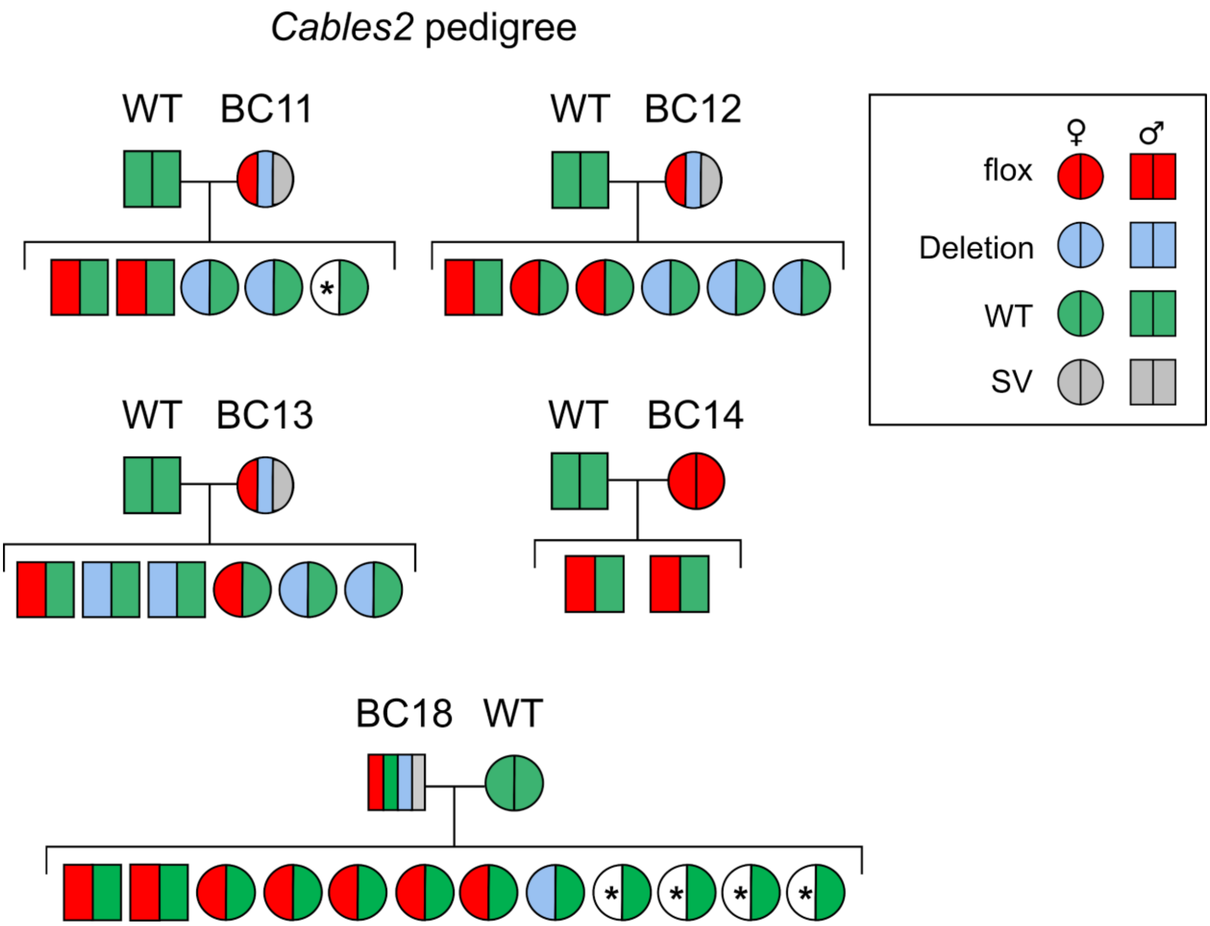


Fig. S19: **Pedigree line of BC11, BC12, BC13, BC14, and BC18 in *Cables2* flox knock-in design**

Alleles that have not been identified are marked with ‘*’.
