## Additional file 2: Fig. S20 for "Multiplex genotyping method to validate the multiallelic genome editing outcomes using machine learning-assisted long-read sequencing"

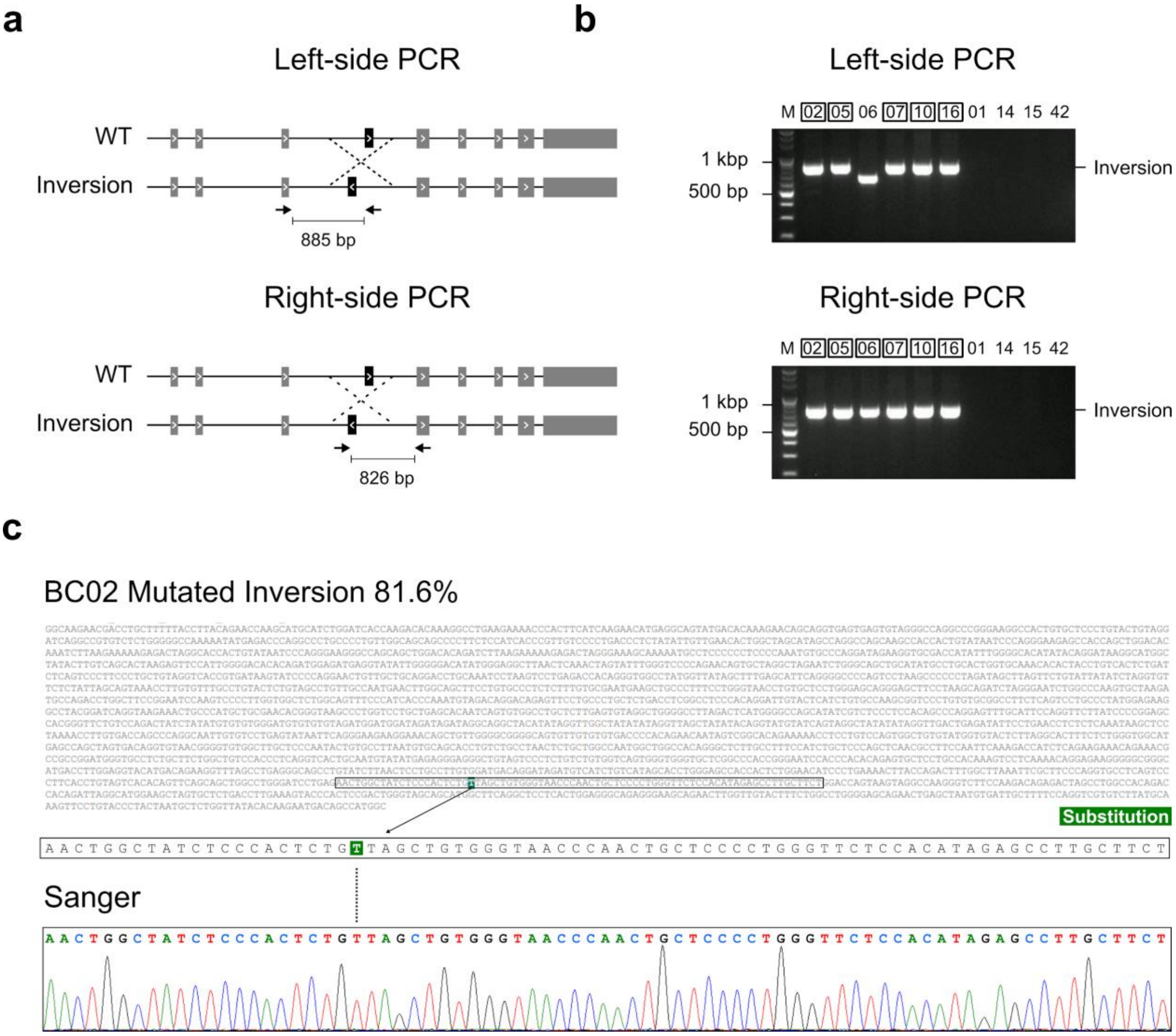


Fig. S20: **Validation of DAJIN-reported inversion alleles**

**a** PCR design to validate inversion alleles. The arrows represent PCR primers for the digested DNA fragments, including the size of PCR products. **b** PCR results for the detection of inversion alleles. The numbers on the panel are the barcode IDs. The boxed number represents the samples with inversion alleles. **c** Comparison between DAJIN's consensus sequence and Sanger sequencing. The sequence represents the consensus sequence of inversion alleles of BC02. The green highlighted nucleotides represent substitution. The dotted lines represent corresponding nucleotides between Sanger and DAJIN's consensus sequences.
