## Additional file 2: Fig. S22 for "Multiplex genotyping method to validate the multiallelic genome editing outcomes using machine learning-assisted long-read sequencing"

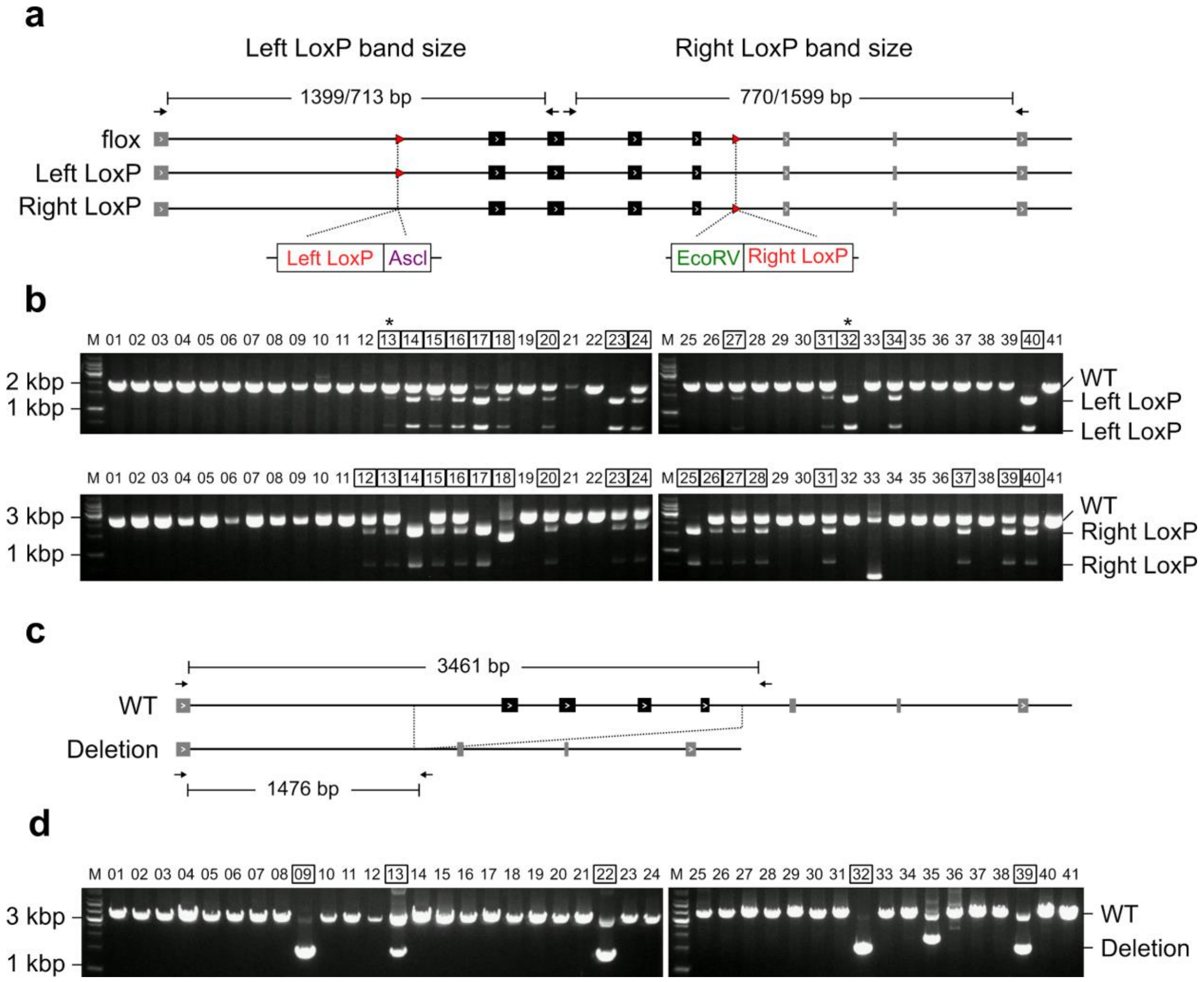


Fig. S22: **PCR-based genotyping of *Exoc7* flox knock-in design**

**a** PCR-RFLP design to validate LoxP knock-in alleles. The *Asc*I and *Eco*RV digest the restriction sites adjacent to Left LoxP and Right LoxP, respectively. The arrows represent PCR primers for the digested DNA fragments, including PCR product sizes. **b** PCR results for the detection of LoxP knock-in alleles. The top and bottom panels represent the DNA fragments digested with *Asc*I and *Eco*RV, respectively. The numbers on the panel are the barcode IDs. The boxed number represents the samples with LoxP alleles. The asterisks represent mismatched samples from DAJIN's genotyping. **c** PCR design to validate deletion alleles. The arrows represent PCR primers for the digested DNA fragments, including the size of PCR products. **d** PCR results for the detection of deletion alleles. The numbers on the panel are the barcode IDs. The boxed number represents the samples with deletion alleles.
