## Additional file 2: Fig. S24 for "Multiplex genotyping method to validate the multiallelic genome editing outcomes using machine learning-assisted long-read sequencing"

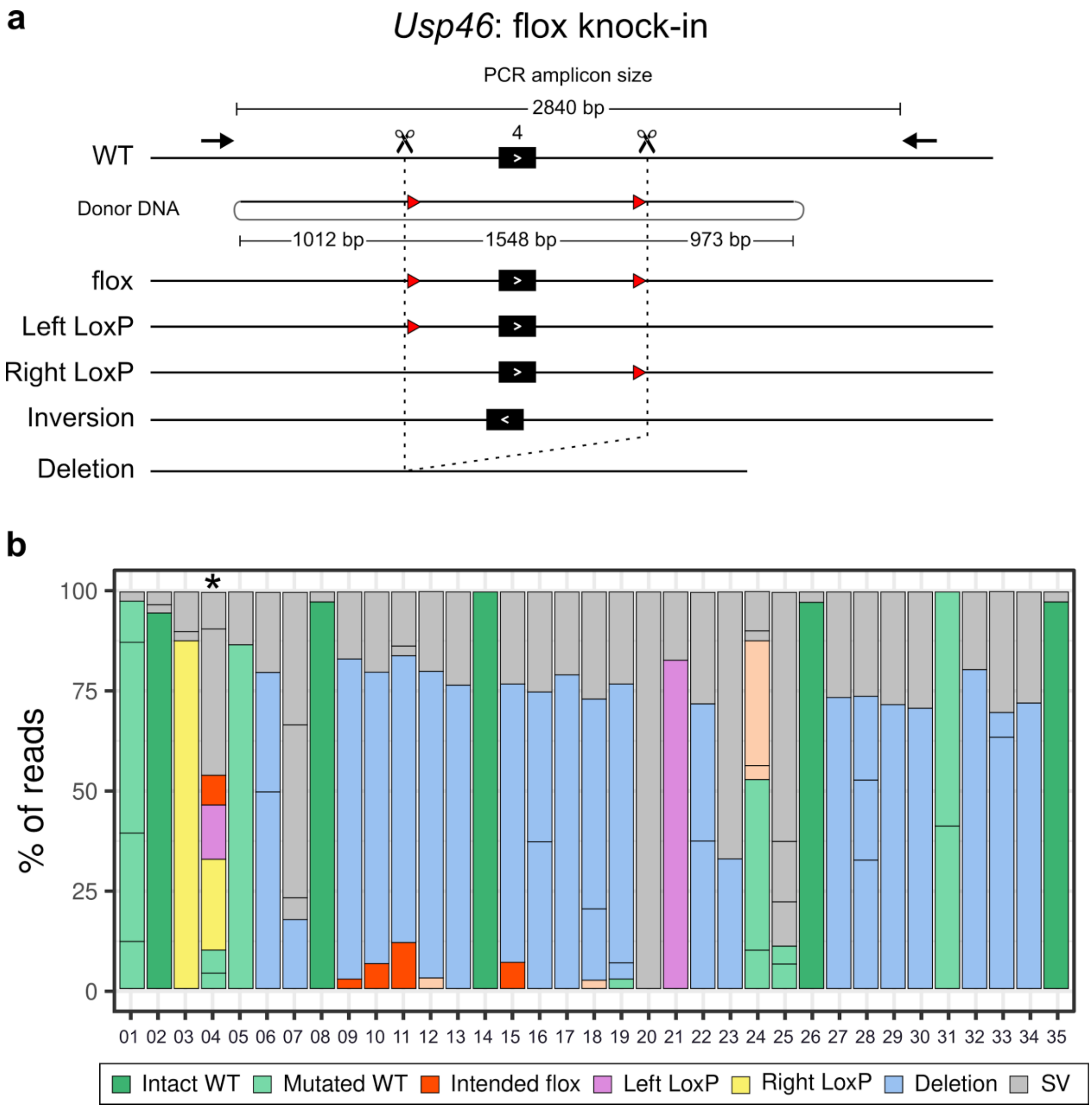


Fig. S24: **DAJIN application to *Usp46* flox knock-in design**

**a** Genome editing design for flox knock-in into the *Usp46* locus. The scissors represent Cas9-cutting sites. The arrows represent PCR primers including the size of the PCR amplicon. The circular DNA represents the donor DNA. The base numbers on the donor DNA describe the size of the left, central, and right arms. The red arrowheads represent LoxPs. The boxed allele type represents the target alleles. The other allele types include Left LoxP and Right LoxP. Inversion and Deletion represents possible by-products. **b** DAJIN's report of the allele percentage. The barcode numbers on the x-axis represent mouse IDs. BC35 is a WT control. The y-axis represents the percentage of DAJIN-reported alleles. The colours of the bar represent DAJIN-reported allele types. The horizontal lines in a bar represent the DAJIN-reported alleles. The asterisk on BC04 represents a pseudo-flox mouse.
