## Additional file 2: Fig. S25 for "Multiplex genotyping method to validate the multiallelic genome editing outcomes using machine learning-assisted long-read sequencing"

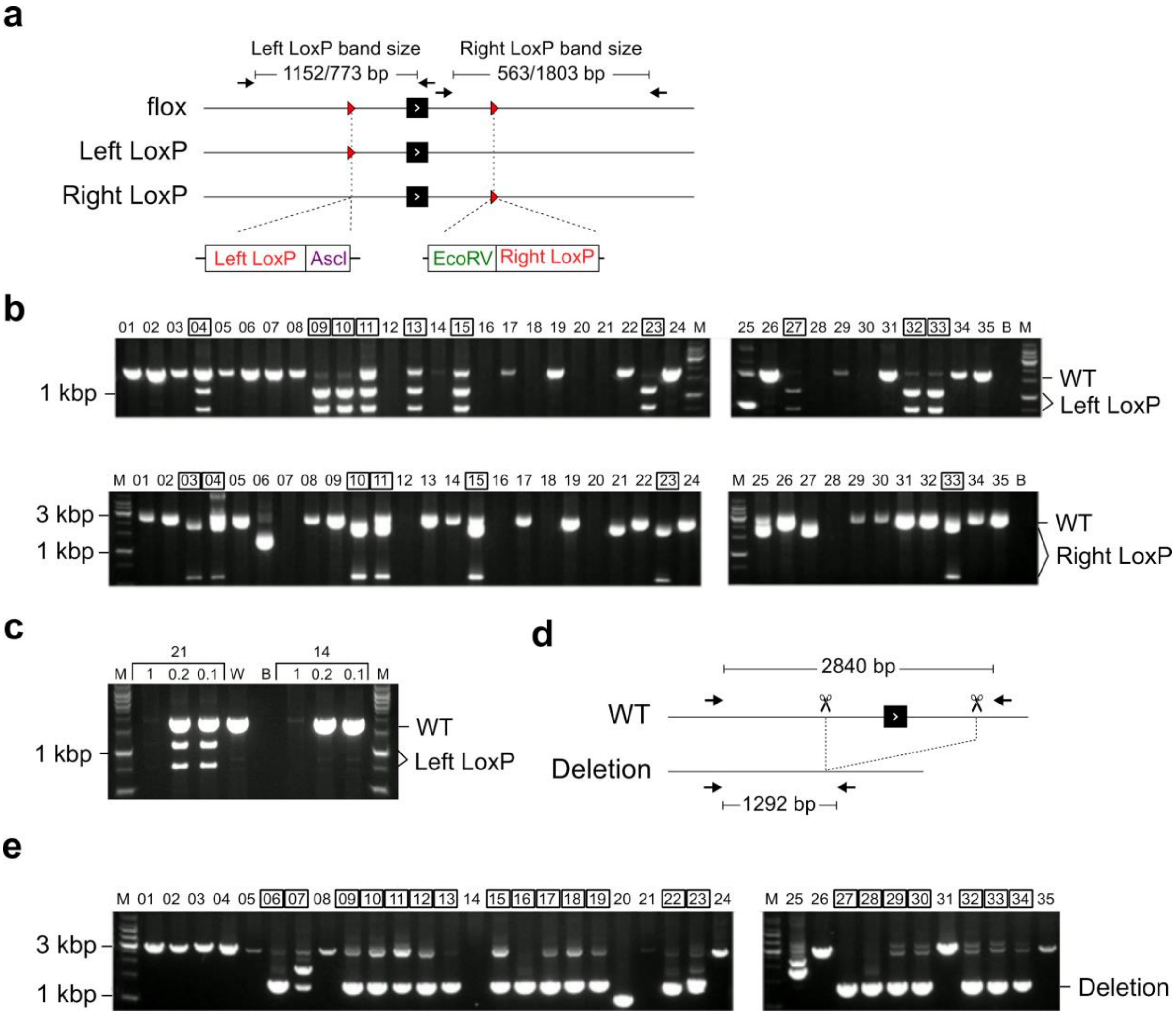


Fig. S25: **PCR-based genotyping of *Usp46* flox knock-in design**

**a** PCR-RFLP design to validate LoxP knock-in alleles. The *Asc*I and *Eco*RV digest the restriction sites adjacent to Left LoxP and Right LoxP, respectively. The arrows represent PCR primers for the digested DNA fragments, including PCR product sizes. **b** PCR results for the detection of LoxP knock-in alleles. The top and bottom panels represent the DNA fragments digested with *Asc*I and *Eco*RV, respectively. The numbers on the panel are the barcode IDs. The boxed number represents the samples with LoxP alleles. ‘M’ and ‘B’ means marker and blank, respectively. **c** PCR results for the detection of DAJIN-reported left LoxP alleles in BC21. The numbers on the panel are the barcode IDs and their dilution condition. The boxed number represents the samples with left LoxP alleles. **d** PCR design to validate deletion alleles. The arrows represent PCR primers for the digested DNA fragments, including the size of PCR products. **e** PCR results for the detection of deletion alleles. The numbers on the panel are the barcode IDs. The boxed number represents the samples with deletion alleles.
