## Additional file 2: Fig. S26 for "Multiplex genotyping method to validate the multiallelic genome editing outcomes using machine learning-assisted long-read sequencing"

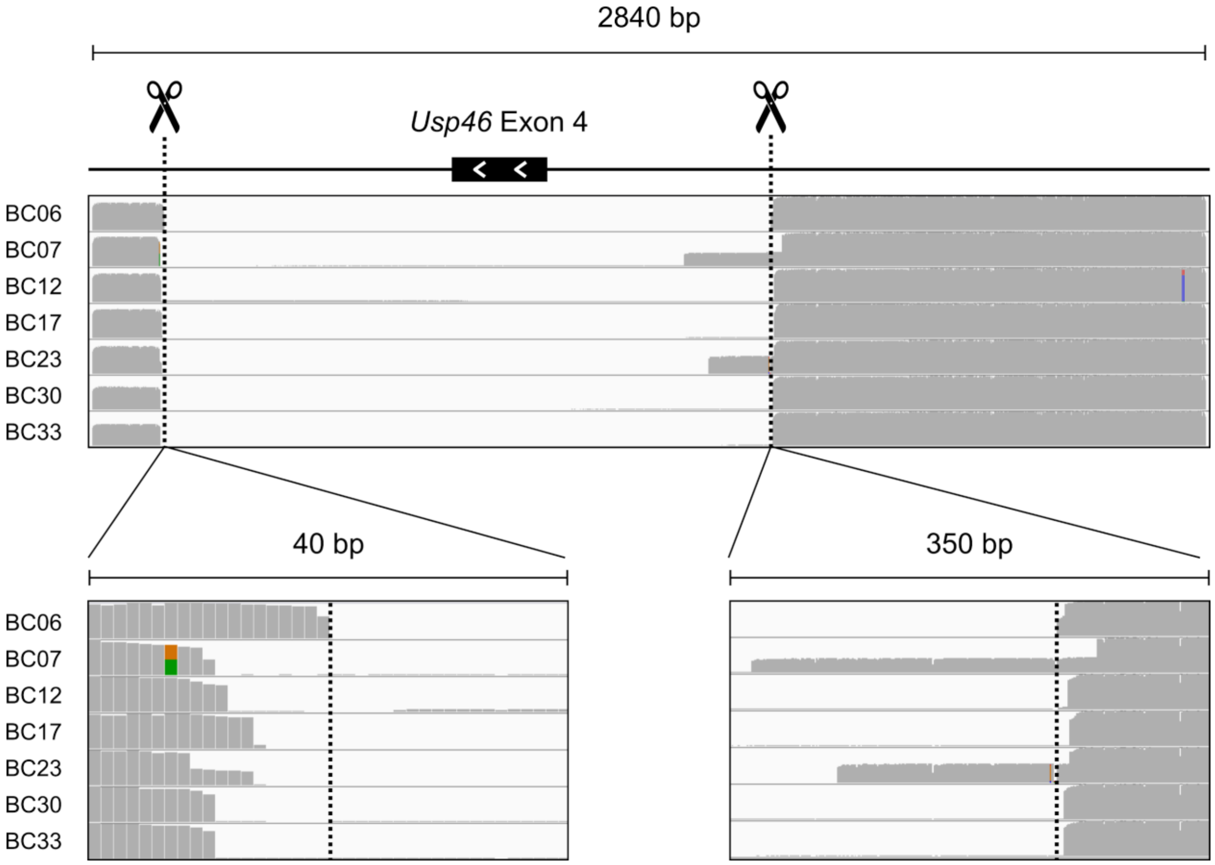


Fig. S26: **DAJIN distinguishes between deletion and SV alleles**

DAJIN labelled the BC07, BC12, BC17, BC23, BC30, and BC33 as ‘SV’. BC06 is a ‘Deletion’ allele as a control.
