## Additional file 2: Fig. S27 for "Multiplex genotyping method to validate the multiallelic genome editing outcomes using machine learning-assisted long-read sequencing"

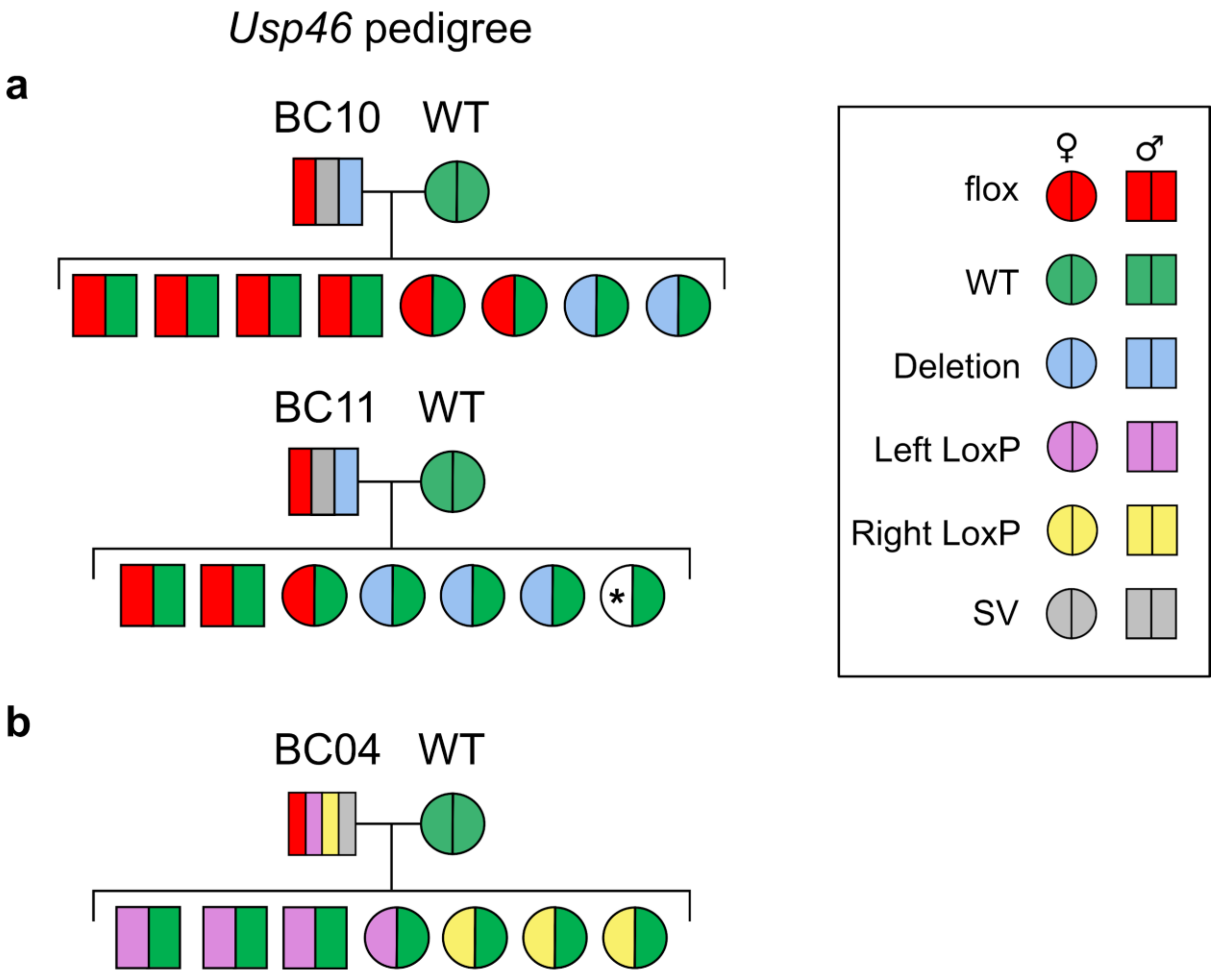


Fig. S27: **Pedigree line of BC10 and BC11 in *Usp46* flox knock-in design**

**a** The flox mouse line which transmitted to its pedigree. **b** The pseudo-flox mouse line.

Alleles that have not been identified are marked with ‘*’.
