## Additional file 3 for "Multiplex genotyping method to validate the multiallelic genome editing outcomes using machine learning-assisted long-read sequencing": Additional_Stx2_barcode25_allele1_SV.html

>barcode25\_allele1\_mutation\_abnormal\_29.8%
GCTCCAGGGTGTCTCATAGTGTTTGAAGGCTCCTAAATTGCCCAGTGTTCAGCTGGGGAAAGACCATCAGCTAGGCAGGATCCAAAGGATAATGAGTGTGCCCCATGGGACTCTGGCTTAGCCGCAGCTCTACCCTAAGCCCACAGTTGAGGGGTAGTGGATCTTGTGTTTGAGGACATTAACAACAGGCTGATTGGGAAAGTAGTGGTTGCTTGATGGAGTTGGGCTAGCGATGGAGGTGAGTGAGTCTGGAGGCCAGTTGTGTGCCGCATACTAACAGAGGAGTAAGCACCAGCTAGATTTTCATTTCCTTAGGATATCTTTCTAAATCTCAGCTCTTTTTAGAAATGTTAAATGCCTTTGTTTAGAAACTTTGAATGTTGGCTGCCACTTACATTAAAAAAAAAAAAAAAAAAAACCTGGAAACTAAGACAGTGTCTGGTCACTCCCACACATCAGTTACTGTGGAGCAGCTTGCTCACCTGTGCTCACTCCACTGCCCTCAAGGAGCTAATCTGCCTGTCTGCTGGCTGCATATTCTGCTTCCTCCTGGGGAGTCTCATAGGGTCTCTGAGATCACACTGTTTCTGGTCACACTGGGGTTTTGTAAGGCCTCCCTTACATCTGCCAGGTTCCAGTTTGCTTCACCTCATTGGGTGTTTTAATGACATCACATCAGATGGCAAGAAAGACTCTGGGTGGAGTCCCCCAGAATCTCCATAGATGCCATAGAGGTATAGACATGGCAGTGGACTGCTGCTGTCAGTTAGCTGCAGCCAGTGACCTTTTACCCCGAGAATGGCTGGGGCACCAAAGTTGGCTCTGAAGGTCTAGAGCCAGGTCTCCAGGAGGGTGACATGCGTGGTATGTCTGATGGAGCAACAGGGACCAGGAAGACAGCGACAGCAGAAGGGACTTCTGGCAGCTTCTTCCTGGGTGTGGTTGCATGTACTCTGTGTCCCAGTGAAGGTTAGCATCAGAGATCTGATTGGGGTGAGCCTGCAGTGCTGGCTCACCCCAGCTGTGTCTGGTAACACAGTTCGGGGTCTACCAGGAGCTGGCCCGGACACGGCACTGGTGGCTGGGGAGGTGAACTTGGACAGGCACTGAGAAAAGAAGGCTGGTTGCTCCTTCTCTGTGCTTGGCCTGCATGCCTGTATCTGTAGTGTAAAGTGGTCCATCTCACATGCTTCATTTCCCCCTTTTAAAGAAATAAAAGAAGAGCTGGAGGACCTGAACAAAGAGATCAAGAAAACTGCTAACAGGATCCGGGGCAAGCTGAAGTGTAAGTTTGCCTTCTTGGGTGCTGGTGGGGTGTGGAAGTTGGTACCTTTTTCCTTAGAGTTTTATTATTATATTTCCCAAAATGAAAATTGAGACTGCTTCTAAGTAGGTTGGTTATCTGTAATCCCGTGCTATATCTCATTGTGACTCAGTAGGTGACATTGGGCAGGTGTTTGAGAAAGTGGGGATAGCTTTCCTAGTGACTCGCCAAGCTCTCAGGCGGGCCTCTTAGCTAGAATGTTCTTTCCAGCCCCTAGAAGCTCAGCTGAGATCATGGTCCTCTGTGTTCAGGATGACGCCTTGGCTGGGTTGAGGGTTGTGGCTTTTTGCACAGTGCTTAAACAGAAGTTGCTTTCCTTGATCTGTGAGGAAGACATATGTTTAACTTTTTTATTTTTTTCTGAGACAGGGTTTCTCTGTGTAGCCCTGGCTGTCCTAGAACTCACTTTGTAGACCAAGTTGGCCTCGAACTCAGAAATCCGCCTGCCTCTGCCTCCCGAGTGCTGGGATTAAAGGCATGCTCCACCACTGCCCAGCTGACATGTGTTTTAACTTACAGAGACACATCAAAAGGAATAAATCTCATCTCCGCACATACCCTCTCCTCAGGAACAGTGACTCCCAGGTAGCAGGCACTCTGCCCAGATCCTAGTAGGCTCCTGGCACTCCTCACTGGGCTCAAGGCAGACCTTAGAGAGTGTGCACCTTAAAAACAGCGTTGTTCCCAAGAGCGCAACCACTTTATGTCTGAGTCAGACTCATTACAGTGATTGGGTGATTAAAAATACCCGGCACACCCCATCTGTACACACAATATACTTACACAAGTGCACACACATGCAAATGAACCAGAGCAACATAGAGACATGTGGTTACACACACACACACACAAACTCAGCATATGAATCCATAAAAACAAACACGTAAGGCCCATGTTCAGTCACATACCTATATGCATATTTGTGCATGATTTTGTTTTTGTGTGTTTTTCTGTATGCATGTGCCTATGATTTTGGTATGCATGTACACATAAAACATGTTTGTGATTGTCCATCCACCTGTCTGTCATTGTCCACAAGTGCGGACAGTGTCATAAGCATTCCTTATGATGGCTGGACTTTAGAGGACAATTTCCAGAGGTCACCTATTCTTTAGTTTTGTGGGGAAAGTTGAGGCCATCCCTGAGACCAATGTCGTATCTTGGTGACTCTGTCCTGCTTCCTTCCAGCTATTGAGCAGAGCTGTGATCAGGACGAGAATGGGAACCGAACTTCAGTGGATCTGCGGATACGAAGGACCCAGGTTGGCCTTCCAGGCTCAGTTAAAGCGATGTGGAAGAGCAGCATTATGCTTTGTCCAGACTCAGAGCAATTTCAGTTTTTATTTTTTTATTGTGCTGGAAGTTGAGGCCAGTATCCTGCACATGCTCAGCAAGGGCTCTGTCACCAAGCCACAGCCGAGCATCACATCTGCCCTTATGTAATGTCTGTCTGCAGGGGGTGGGGGTGGGGCTGGGGGGCTGCATCTCTTCCATGTGAAGAGATGACTCAGCGACTCGGCTACCTATATGCTCTGTGCCTCAGAGTAGAGGACTAGCCAGACCACCTAGATGGTAGTGGTTTGGTTTTAATCAGTTTTTTTTTTTAAAGATTTATTTATTTATTTTATGTATATGTGTACACGGTCGCTGTCTTCAGACACACCAAAAGAGGGCATCAGATCCCATTACAGATGGTTGTGAGCCACCATGTGGTTTGCTGGGAATTGAACTCAGGTCCTCTGGAACAGCAGTCCATGCTCTTAACCACTGAGCCATCTCTCTAGCTGGGTTTTCAACCTTCCTAATGCTGTGACCCTTTAATATATAGTTTCTCATGTTGTAATGACCCCAATCATAATATTATCTTCATTGCTACTTTATAACTGTAACTTTGCTTGTTATGAATTGTAATATAAATGTCTGACATGCAGGATATCTGCTATTTGACCCCAAAGTGGTTGTTACCAAAAGCCTTAACTATTAGCCTTAACTATTCAGGGTTTATGGTTGAAACTGGGATGCTCAACCCTGTTTGACCAAGCCTGCCAGCTTAGGGCAGGCAGTCCACGTGCCTGCCAGCTGGATTCAACCTGTGATAACGCCCTCCCTCTAGAGGTCAGCTGGCTGCTCCGGGCTCCAACTTATTTGTTGTTGTTATTGGTGATGGTGGTATTTTGTTTTGTTGAGCCAGAATCTTACTCTGTATCCCTGGCTGGCCTGGAGTGAGCTTGTTACATAGACCAGGTCGACCTTGAACTCACAGAGATTTATTTCCCTCTCCCTCTGTATTGGAATTGAAGGCATGCGCCACCGTCCTGGTGGCCCAGGTCATGCACCTCCCATTGCCGCTCTTTCTCTTACTTTCTTCTGGGATGGAGTCATGGTTGGGGCTTCTTAGTCGTCAGTTTTACCTCTTTCAAAGGAAGGGGTTGCCCCCACATAGTAACACGGTATTCTCCGTGTTCTAGCACTCGGTGCTGTCACGGAAGTTTGTGGACGTCATGACAGAATACAATGAAGCGCAGATCCTGTTCCGGGAGCGAAGCAAAGGCCGCATCCAGCGCCAGCTGGAGATCAGTGAGTAGGGCGCATGCGGGAGACGTGCTTCAGGACCCATCAGATGGCAAGCGCCCCACTTCTAACTGCAGAGTAAACCGAAAGGCCCTTTGCTGCAGCCTGTGCCTAGAGTCAGGCAGATGTGTTATGTGTCGTAAGCTTCAGAGTTCTGAGTGCGGGGCTGTAGTGGCTTGCGATTGCTGGTGCTGCTATTTCACGCATCCCGAAATCTTGTGGTCCTGATTTCTTAAGGCATTTTCGTGTCCTGCAGCTGCTCCTACCTGGCTGGGAGAGGATCATCCTCATGTGCCTCCTTCAGCTTAACTCCCTCTGGCTAGAAGCCCTGTGGGACCCGAGCATAGCTGCTCCTCTGTAACACCAGGGTAGGGTTGTGACACATTGCATCACATTTCTGCCACATTACTGTGT

---

Insertion Deletion Substitution
