## Additional file 3 for "Multiplex genotyping method to validate the multiallelic genome editing outcomes using machine learning-assisted long-read sequencing": Fig2_Tyr_barcode08_allele4_mutation_target.html

>barcode08\_allele4\_mutation\_target\_61.4%
TGCATTGAAGCAGTTCACCAAAATAACAAAGTAACAAAGTAAGATATCTTTGGAATAATCAATTCAAGATAATCAAGGAAAAATGAGAGGCAACTATTTTAGACTGATTACTTTTATAAAATAAATAAGCTCAGCTTAGCCAGATATAAGCAATATTCTGAGTTCTGAAGAAAAATTTTTGACAAAATGAGTTCTATAAATGTTATTGTCTACTTATGATCTCTAAATACAACAGGCTTGTATTCAGAATCTAGATGTTTCATGACCTTTATTCATAAGAGATGATGTATTCTTGATACTACTTCTCATTTGCAAATTCCAATTATTATTAATTTCATATCAATTAGAATAATATATCTTCCTTCAATTTAGTTACCTCACTATGGGCTATGTACAAACTCCAAGAAAAAGTTAGTCATGTGCTTTGCAGAAGATAAAAGCTTAGTGTAAAACAGGCTGAGAGTATTTGATGTAAGAAGGGGAGTGGTTATATAGGTCTTAGCCAAAACATGTGATAGTCACTCCAGGGGTTGCTGGAAAAGAAGTCTGTGACACTCATTAACCTATTGGTGCAGATTTTGTATGATCTAAAGGAGAAAATGTTCTTGGCTGTTTTGTATTGCCTTCTGTGGAGTTTCCAGATCTCTGATGGCCATTTTCCTCGAGCCTGTGCCTCCTCTAAGAACTTGTTGGCAAAAGAATGCTGCCCACCATGGATGGGTGATGGGAGTCCCTGCGCCCAGCTTTCAGGCAGAGGTTCCTCTGCCAGGATATCCTTCTGTCCAGTGCACCATCTGGACCTCAGTTCCCCTTCAAAGGGGTGGATGACCGTGAGTCCTGGCCCTCTGTGTTTTATAATAGGACCTGCCAGTGCTCAGGCAACTTCATGGGTTTCAACTGCGGAAACTGTAAGTTTGGATTTGGGGGCCCAAATTGTACAGAGAAGCGAGTCTTGATTAGAAGAAACATTTTTGATTTGAGTGTCTCCGAAAAGAATAAGTTCTTTTCTTACCTCACTTTAGCAAAACATACTATCAGCTCAGTCTATGTCATCCCCACAGGCACCTATGGCCAAATGAACAATGGGTCAACACCCATGTTTAATGATATCAACATCTACGACCTCTTTGTATGGATGCATTACTATGTGTCAAGGGACACACTGCTTGGGGGCTCTGAAATATGGAGGGACATTGATTTTGCCCATGAAGCACCAGGGTTTCTGCCTTGGCACAGACTTTTCTTGTTATTGTGGGAACAAGAAATTCGAGAACTAACTGGGGATGAGAACTTCACTGTTCCATACTGGGATTGGAGAGATGCAGAAAACTGTGACATTTGCACAGATGAGTACTTGGGAGGTCGTCACCCTGAAAATCCTAACTTACTCAGCCCAGCATCCTTCTTCTCCTCCTGGCAGGTAAGATGCACTATATAGAGAGAGTTGCAAAGACTGGTACTTCAGCAGCCACATTTTCATGCTCTGTGAGCATCTCTGATAATATCTCAGGGCAGAAAATGTGCCTTACTAACAGATGTTAATGCTTCTTGATTTCTTTTTCTCTTTTGAGAACTCTTCAAAGTTGTTATTAAACAAATATCTATGTGCTTATTTGTCTTAATATCTAACAGCTTAGTTAGATTTCTAAGCTGCTATAAACAAGGACTGATTGGTTCACCACTGTATTGTTAGCACCTCCTATGGTATCTGGAATAAACAGTAACTCAGTTATTTAAGAATGGATGAGAAACCAGATTATCTTAGTTCATGTTTCTGAGTAATATTAAAATTAATATTAACAGTAAAATCCATAAGTATGCTACTTTAAAATATAAATCTCTGGCCAAAACCAAGACTTATTATTCAGGATCTTCAAGAGAAAGTGCTGAGATAATTCACTAAGTATCAGAGATGACCTTTATTACATGATTGCCTGATAGAAAAAATGATTACACACACACAAAAAAATCTTCAGTTGCTTAAATTTTAAACGTTGCTGACTCTCAAACAGTTAAGTAATAAAAGAGTTAAAGCCTGCTGTGTATTTAGAATATGTGAATACCTATTGAAAGAATTTATTGTACAATTAATATAAACAGACTTCTATTTTACAGTCATAAGATACTACTTAATTTGTTAAAAATTATTTTTTGATAGCATTGTTGGTAAATAGCAAAGGTGATATTTGCTAATGATTACAAGGGCTGTCTGGCTAACTTACGTTATGTTCAGGGAGAAGACAGTCCTTTTTAAGGAATGGGCACTTTCTAACTTTTTTTCTCTAGGATGGAGAAAAATTAGCCTTCTTCCTACTTTAAAAATGTTAGACATAGAATTAAGGGATTGTTATTTTGAGATTAAATTTTCTTTTCTCCTATTATTTTTCCTCATTCTGGAATGGAAGCAAAAGATGAAGAAAGAAATATATGTTAAATTGTTTTCCTTTAAATGAACACAAATGTGAAATATGTTTTTCTGCCTATCTTGTAAAATTTTCTATTGCAACTATTCTGATTACCAGTTCAAATGGGGAAAAAAGAACATAGGCTACCCCACACTTGAAATTTTGAAATATGAATGTCCTCTGTCTCTGCTGGTCTAACACTTCCAAAATGGAAACCTTTAAAGGGCCACTGTAAATTACAGCTGCTAATTCCTGGTGCCAATGGTGATAAGTGTTTACTAAACCTAGTGAGTACTTTATAGCATGGGGCTCTGCTGCGAAGTAACATTGCTGTATATTTTCAGTCATTCTACCTTAATTCATGAACTGCAAAACTCTCATCTAGCTTTTTACTTCTCTAGCTATTGCTTTAAGTTCTATCAGGCTCAGGTGTGGAATTCTC

---

Insertion Deletion Substitution
