## Additional file 3 for "Multiplex genotyping method to validate the multiallelic genome editing outcomes using machine learning-assisted long-read sequencing": Fig3_Prdm14_barcode18_allele3_mutation_target.html

>barcode18\_allele3\_mutation\_target\_15.7%
CTTGTACTCAAAACCTTCTGCCCCAACTCCCCAAGTGCTCCTGACTGTTGAACTGTGTGTGAGTCTGTGTGCGTGTGTCTATGGCTGTGTGTGTGTGTGTGTGTGTGTGTGTGTGTGTGTACACCTGGAGATCTTGTAGAGAGGCTAGCTAGTTCTTAGCACAGGTGGGGCTGGCACTATGACTATGCGCCTTGCTCAGCTTCCAGGGGTTGCTGCTGGTCCCTGGATGATCACATCTGAGTGGCTAGGAGGCAGAGGACAATCAACTCACCATAGGAATAGGTCTTCAGATTTGGTGTTTTTAAGGAAAGAGTGACGCTGAGCCCTCAGACTATGCTAGAATTTTCAGGTTTGTGTTTTTTTGTTTTTTTCTTCTTTGTTCCCTGTCTCAAGTAGTCCAGTATGGATGGCTGGACTGCTCTTCCTACTTCCGACCCCCAAGTTCTAGGATTATAAGCATGTAAAGCTATACCCAGCTTTAACTTGTTTGTTTGTTTGTTTGTTTGTTTTTTGACACCTGGTTTCACTAAGTAGCTGTTCTGAAACTCACTGTGTAGACAGGGTTGAACTCCTGACTTCCTCTGGCTCCTGGATGCTGAGGTTTAAAGTGTGTACCATCGCACTAAGCTTGCAATCTACTCCTAAATATTACCTGTCCAACTGGTTTGTATACATAAACATTTGGAAACTACTGACTCAGTAGGTGAGGAAAAATATAAACTACTGTGTGTATAAAAGGAAGACATCATCACTACAGTGTAATAAGCAGCCCAAAGAGCAGGTAAAACAACACATACATGTTATCTGTCACACTATGGTGATGCTCAGTGTGCCCAGCCTGGTCTACACAGTGAGTTCCAGGACAGCCAGGGCTACAAAGAGAAACCCTGTCTCGAAAAACAAAACAAAACAAACAAACAAACAAAAAGTATGCACACCACACTCTGCCCGTGATACCATTTCTGACAAGGTGTGAAAGGCAGCTTTTAAAGATTAATTTATATTGAGAAGTCTTACAAGTGACTTCCTATGATGACTGTTAATATTGAGTGTCTAGTTTTGTTCCATTGCTGTGTTGAGGATTAAATCAAACTCGGGTCCTGACTCCTGCCAGGCCAGTGCTCTGCTGCTGATCCACACCCCAGCATAAGGTCAATGTTTTTGGAGATGATTAATAATAAAAGAAAAAAATAGTGGTTGTGTGTGTGTAATCTTGATACTTGCAAGGTTGAGGAAGAAGAATTTTAAATTCAAATCTAGTGCGGGTTACTCAGGGAGATCCTGCGTCAAACAAAAACCCACAGCATTTTGGTAGGGTTGGTGGGTGCCTTCCCTCACAAGGTCTGGGGCTGGAAAGGCGGACTCTTGACTTAGTGTCATAAGTGGACTGTTTGAACAGACCAGGCATGCCTGGGGATGAAGCAGGAATGACTCACAGACAACAGCTGCGTCACTATAACCCCAGCCTAGGCTATATCTCACAAAGCAGGGAAACCTGGAGCACACTGTACTCTCAACAGACTGGAGAGTGTCCTTTCCATGTGCCTCAGTTGGTCTAAACCTCTTCCAGGCAGCTCACCTGGTTCCTGCCTCTTCCAGGTCTGCTCTTTGCAGCCTCCTGCCCTCTGAGAAGGGCTCTTAGCTTTGATTGCTTGCTCTCACAAGGAGTTACCTAGTGAATCCAGTTAGTTTCAGGGATCCCTGAAAGTATTTGGAGTTTTGGTTTATTTTTTGCTTTTTGAGACAGGTTTCTCTTTGAAGCCCTCCCTGGCTGTCCTGGAACTTGCTCTGTAGACCAGGGTGACCTTGAAGCCTGAGATCTGCCTGCCCCTGTCTCCCAATGTGTTAGAATTATACTGTTTTGTATTGTGACATTTACATGAGATCATGGTGATAAACAAAAAAGGGAAGGCTACGGTTTAACAGTTATGTGTTTGCCAAGTAGATGGGGGTCTGTGGTGTGGCACCATCACCCAGCCTTCTGCAGATTTCTACTTACGCTCCCTCCTGGTTCCTCTCCCTTGAGGGCAGACCGTAAGGTCTCTGGTTCTAGTATCCAGGAACATCTTCAGGCTCTAGATAGTCACTGGTGCAGAAGAAACACTGACATGGTCCAAAGAATGGAAGGCTACATCACAGTCTGTGTTCTTTCTTTTTTTTTTTTTAAGATTTATTTATTTATTATATGTAAGTACACTGTAGCTCCAGAAGAGGGCGTCAGATCTTGTTACAGATGGTTGTGAGCCACCATATGGTTGCTGGGATTTGAACTCTGGACCTTTGGAAGAACAGTCGGGTGCTCTTACCCACTGAGCCATCTCACCAGCCCCGCAGTCTGTGTTCTTTATGAGGATGAGAGACGACAGTGATGACAGGAAGCTGTTCTCTGCCAGCCCAGACACCCATTGCTCTGCCAGGCTTGGTCAAAAGGGCAAAACCAAGTTTGTCCTTAAAAGACAAGATGGAGTTCTGTGTTTTGTTACAGTGTTTATAGTTCTGTGTACATGTGTGCGTTGTGCATGTGTGAGTAGAGTGCCCCTGAGAGGACAGAAGTGGCCACCAGAGCCCCTGGATTTGGGGTTGCAGATGGTTGTGAGCCACCATGGTTGTGCTGGAAACTGAAGTGCAGTTCATGCATTTAGCCACCAGGTCGGCTCGCCAGCTCCTTGTCATACATAGACTCTTATAGTACATCTTATACTAGTGTTACACTAGTGTATAATATGCAGCGTCATATGCCAGTGCTAACATTTATAGTTTTCCACATTTTAATATCTCTAAATTGATCTGTGTTCTACAACTGGGCCATAGTTTTTTTCTTTTTTTTTTTCTTTCTTGTTTTTGTTTTAATGTTTTTAAAATCAGGGGGTTTTTTTTTGTTTTGTTTTTTGTTTTCTTTGGTAAAGGATACATTTATTTTTATTTTATGTGTTTTGCATGCATGAATGCACACCATGTACATTCAGTTGCTTACAGAGACCAGAAGAAGGCATCAGGTCCCTTGGAACTAGAGTTGCTACTGACAGTCAGTTGTGAGTCACCATCTGGATGCGGGGAACCAAACGTGGGTCCTCAGCTAGAACAGCAATTGCTCTTCAGTCAGCTCTCCAGGCCCTAAATTCTGGGTTGCTTGTTTGTTTGTTGTTGTTGTTGTTGTTACCTAATTTATGTGGTGGTGTGTATGTGTGTATTGTTGTTGTTGTACATATCTGTGTATACCTCTTTGGAGTCTTTCCTCTTTATCACTTTCTGCCATAGTTTTTTGAGAAAGGGTCTGTCTTAGTCAGGGTTTCTATTCCTGCACAAACATCATGACCAAGAAGCAAGTTGGGGAGGAAAGGGTTTATTCGGCTTACACTTCCATACTGCTGTTCATCACCAAGGAAGTCAGGACTGGAACTCAAGCAGGTCAGAAAGCAGGAGCTGATGCAGAGGCCATGGAGGGATGTTCTTTACTGGCTTGCCTCCCCTGGCTTGCTCAGCCTGCTCTCTTATAGATCCCAAGACTACCAGCCCAGAGATGGTCCCACCCACAAGGGGCCTTTCCCCCTTGATCACTAATTGAGAAAATGCCTGACAGTTGTATCTCATGGAGGCATTTCCTCAACTGAAGCTCCTTTCTCTGTGGTAACTCCAGCTGTGTCAAGTTGACACAAAACTAGCCAGTACAGGGTCTTTCACTAAAGCTGGAGTTAGGCTGGCAGTCTACAAACCCCAGCAATCCTCTTGTCTCTCCTTCCCAGAGCCCTGGGGTGCAAGTCGTGGTTGGCCGTGCTTGCTTTTTAATTAAATGGCTGCTGGGGGTTGAATACAGGTCATGCTTTTGCATCCACTGAGCCACCCACCCAGCCATCGGAAAGTGCTTGGGTCCCTTCTCTCTAGTTTACAAGGTCATTTATTTTGTCCCTCATAGTAACAAAAGCTCTGCTTAACCCTGGGTTTAAAATCTCTCCTGAACACAAGCAGCACCTATCCACCGAAGCTGCTCCAAGAGTGCCTCACCCTGAATTTCTTGTGTGCTTTCCCTAACAGTCATGTCCAGAG

---

Insertion Deletion Substitution
