## Additional file 3 for "Multiplex genotyping method to validate the multiallelic genome editing outcomes using machine learning-assisted long-read sequencing": Fig4_1bp_deletion.html.html

>barcode33\_allele3\_mutation\_target\_55.4%
GGCAAGAACGACCTGCTTTTTACCTTACAGAACCAAGCATGCATCTGGATCACCAAGACACAAAGGCCTGAAGAAAACCCACTTCATCAAGAACATGAGGCAGTATGACACAAAGAACAGCAGGTGAGTGAGTGTAGGGCCAGGCCCGGGAAGGCCACTGTGCTCCCTGTACTGTAGGATCAGGCCGTGTCTCTGGGGGCCAAAAATATGAGACCCAGGCCCTGCCCCTGTTGGCAGCAGCCCCTTCTCCATCACCCGTTGTCCCCTGACCCTCTATATTGTTGAACACTGGCTAGCATAGCCAGGCCAGCAAGCCACCACTGTATAATCCCAGGGAAGAGCCACCAGCTGGACACAAATCTTAAGAAAAAGAGACTAGGCACCACTGTATAATCCCAGGGAAGGGCCAGCAGCTGGACACAGATCTTAAGAAAAAGAGACTAGGGAAAGCAAAAATGCCTCCCCCCTCCCCAAATGTGCCCAGGATAGAAGGTGCGACCATATTTGGGGCACATATACAGGATAAGGCATGGCTATACTTGTCAGCACTAAGAGTTCCATTGGGGACACACAGATGGAGATGAGGTATATTGGGGGACATATGGGAGGCTTAACTCAAACTAGTATTTGGGTCCCCAGAACAGTGCTAGGCTAGAATCTGGGCAGCTGCATATGCCTGCACTGGTGCAAACACACTACCTGTCACTCTGATCTCAGTCCCTTCCCTGCTGTAGGTCACCGTGATAAGTATCCCCAGGAACTGTTGCTGCAGGACCTGCAAATCCTAAGTCCTGAGACCACAGGGTGGCCTATGGTTATAGCTTTGAGCATTCAGGGGCCCCAGTCCTAAGCCCCCCTAGATAGCTTAGTTCTGTATTATATCTAGGTGTTCTCTATTAGCAGTAAACCTTGTGTTTGCCTGTACTCTGTAGCCTGTTGCCAATGAACTTGGCAGCTTCCTGTGCCCTCTCTTTGTGCGAATGAAGCTGCCCTTTCCTGGGTAACCTGTGCTCCTGGGAGCAGGGAGCTTCCTAAGCAGATCTAGGGAATCTGGCCCAAGTGCTAAGATGCCAGACCTGGCTTCCGGAATCCAAGTCCCCTTGGTGGCTCTGGCAGTTTCCCATCACCCAAATGTAGACAGGACAGAGTTCCTGCCCTGCTCTGACCTCGGCCTCCCACAGGATTGTACTCATCTGTGCCAAGCGGTCCCTGTGTGCGGCCTTCTCAGTCCTGCCCTATGGAGAAGGCCTACGGATCAGGTAAGAAACTGCCCATGCTGCGAACACGGGTAAGCCCTGGTCCTGCTGAGCACAATCAGTGTGGCCTGCTCTTGAGTGTAGGCTGGGGCCTTAGACTCATGGGGCCAGCATATCGTCTCCCTCCACAGCCCAGGAGTTTGCATTCCAGGTTCTTATCCCCGGAGCCACGGGTTCTGTCCAGACTATCTATATGTGTGTGGGATGTGTGTGTAGATGGATGGATAGATAGATAGGCAGGCTACATATAGGTTGGCTATATATAGGTTAGCTATATACAGGTATGTATCAGTAGGCTATATATAGGTTGACTGAGATATTCCTGAACCTCTCTCAAATAAGCTCCTAAAACCTTGTGACCAGCCCAGGCAATTGTGTCCTGAGTATAATTCAGGGAAGAAGGAAACAGCTGTTGGGGCGGGGCAGTGTTGTGTGTGACCCCACAGAACAATAGTCGGCACAGAAAAACCTCCTGTCAAACATAACTTCGTATAGCATACATTATACGAAGTTATGGCGCGCCCAGAGTGGGAGATAGCCAGTTCTCAGGATCCCAGGCCAGCTGCTGAACTGTGTGACTACAGGTGAAGGGACTGAGGCACCTGGGAAGCGAATTTAAGCCAAAGTCTGGTAAGTTTTCAGGGATGTTCCAGAGTGGTGGCTCCCAGGTGCTATGACAGATGACATCTATCCTGTCATCCACAAGGCAGGAGTTAAGATACAGGCTGCCCTCAGGCTAAACCTTCTGTCATGTACCTCCAAGGTCATGCCCGCCCCCTTCTCCTGTTTTGAGGACTTTGTGGCAGGAGCACTCTGTGTGGGTGGATTCCCGGTGGGCCGAGCACCCACCCACTGACCACAGACAGAGGGACTACAGCCCTCCCTCTCATATACATTGCAGTGACCTGAGGGTGGACAGCCAGAAGCAGAGGCACCCATCCGGCGGCGTTTCTGTTTCTTCTGAGATGGTCTTTGAATTGGAAGGCGTTGAGCTGGGAGCAGATGGAAAGGCAAGAGCCCTGTGGCCAGCCATTGGCCAGCAGAGTTAGGCAGACAGGTGCTGCACATTAAGGCACAGTATTGGGAGCAAGCCACACCCCGTTACACCTGTCACTAGCTGGCTCATGCCACCCAGAGAAAGTGCCTAAGAGTACACCATACACAGCCACTGCGCGCCAATTCGATATCAAGCTATAACTTCGTATAGCATACATTATACGAAGTTATGCGGCCCTAGCTGTGGGTAACCCAACTGCTCCCCTGGGTTCTCCACATAGAGCCTTGCTTCTGGACCAGTAAGTAGGCCAAGGGTCTTCCAAGACAGAGACTAGCCTGGCCACAGACCACAGATTAGGCATGGAAGCTAGTGCTCTGACCTTGAAAGTACCCACTCCGACTGGGTAGCAGCATGGCTTCAGGCTCCTCACTGGAGGGCAGAGGGAAGCAGAACTTGGTTGTACTTTCTGGCCTGGGGAGCAGAACTGAGCTAATGTGATTGCTTTTCCAGGTCGTGTCTTATGCAAAGTTCCTGTACCCTACTAATGCTCTGGTTATACACAAGAATGACAGCCATGGC

---

Insertion Deletion Substitution
